# Adaptive fitness in *APC*-mutant cells requires chromosome rearrangements

**DOI:** 10.1101/2020.09.18.303016

**Authors:** Yoshihiro Kawasaki, Tomoko Hamaji, Koji Owada, Akiko Hayashi, Yuping Wu, Taisaku Nogi, Miwa Okada, Shoko Sakai, Naoko Tokushige, Shota Sasagawa, Hidewaki Nakagawa, Yuta Kouyama, Atsushi Niida, Koshi Mimori, Toshihiko Kuroda, Takao Senda, Miho Ohsugi, Katsumi Fumoto, Akira Kikuchi, Per O. Widlund, Kazuyuki Kiyosue, Norio Yamashita, Masahiko Morita, Hideo Yokota, Satya N. V. Arjunan, Wei-Xiang Chew, Koichi Takahashi, Wesley R. Legant, Bi-Chang Chen, Eric Betzig, Ron Smits, Riccardo Fodde, Hiroko Oshima, Masanobu Oshima, M. Mark Taketo, Tetsu Akiyama, Yuko Mimori-Kiyosue

**Affiliations:** Laboratory for Molecular and Cellular Dynamics, RIKEN Center for Biosystems Dynamics Research, Kobe 650-0047, Japan; Laboratory of Molecular and Genetic Information, Institute of Molecular and Cellular Biosciences, The University of Tokyo, Tokyo 113-0032, Japan; Laboratory for Cancer Genomics, RIKEN Center for Integrative Medical Sciences, Yokohama 230-0045, Japan; Department of Surgery, Kyushu University Beppu Hospital, Beppu 874-0838, Japan; Division of Health Medical Computational Science, Health Intelligence Center, Institute of Medical Science, The University of Tokyo, Tokyo 108-8639, Japan; Nico-tama Coloproctology Clinic, Tokyo 158-0094, Japan; Department of Anatomy, Graduate School of Medicine, Gifu University, Gifu 501-1194, Japan; Department of Life Sciences, Graduate School of Arts and Sciences, The University of Tokyo, Tokyo 153-8902, Japan; Department of Molecular Biology and Biochemistry, Graduate School of Medicine, Osaka University, Osaka 565-0871, Japan; Center for Infectious Disease Education and Research, Osaka University, Osaka 565-0871, Japan; Department of Microbiology and Immunology, University of Gothenburg, Gothenburg 405 30, Sweden; Biomedical Research Institute, National Institute of Advanced Industrial Science and Technology, Osaka 563-0026, Japan; Image Processing Research Team, Center for Advanced Photonics, RIKEN, Saitama 351-0198, Japan; Laboratory for Biologically Inspired Computing, RIKEN Center for Biosystems Dynamics Research, Osaka 565-0874, Japan; Department of Physics, Faculty of Science, University of Malaya, 50603 Kuala Lumpur, Malaysia; Janelia Research Campus, Howard Hughes Medical Institute, Ashburn, VA 20147, USA; Department of Gastroenterology and Hepatology, Erasmus MC Cancer Institute, University Medical Center Rotterdam, 3015 CN Rotterdam, The Netherlands; Department of Pathology, Erasmus MC Cancer Institute, Erasmus University Medical Center, 3000 CA Rotterdam, The Netherlands; Division of Genetics, Cancer Research Institute, Kanazawa University, Ishikawa 920-1192, Japan; Colon Cancer Project, Kyoto University Hospital-iACT, Graduate School of Medicine, Kyoto University, Kyoto 606-8501, Japan; Department of Molecular Genetics, Institute of Biomedical Science, Kansai Medical University, Osaka 573-1010, Japan

## Abstract

Cancer cells tolerate copy number alterations (CNAs) of genomic regions that are lethal to non-cancer cells. Certain CNAs are preferentially associated with specific cancer types and lineages, but the mechanisms underlying the emergence and selection of specific CNAs remain unclear. *Adenomatous polyposis coli* (*APC*) mutations induce mitotic errors, but their impact on tumor evolution remains elusive. We investigated APC function in cultured cells and tumors and found that its loss led to β-catenin accumulation at centrosomes, suppressing its maturation through inhibition of key centrosome regulators, including Aurora kinase A (AURKA) that promotes tumor growth. These defects collectively reduced cellular fitness, leading to impaired mitotic fidelity and delayed cell cycle progression. However, in *APC*-mutant tumors, AURKA activity was maintained, at least in part, through the amplification of chromosomes harboring *AURKA* and its activator genes, yet this alone was insufficient to fully restore proliferation: aberrant chromosomal reorganization also emerged and contributed to the adaptive fitness of *APC*-mutant cells. Such a process of adaptive CNA selection provides a framework for understanding how specific CNAs are selected to counteract disadvantages imposed by genetic alterations during tumor progression, providing one key insight into how specific CNAs are selected in this context.

## Introduction

Chromosome instability (CIN) induces aneuploidy, a hallmark of cancer (1). However, the extent to which aneuploidy itself contributes to tumor initiation and progression remains unclear. Advances in genome characterization technologies have revealed the genomic landscape of cancer, identifying recurrent copy number alterations (CNAs), which include whole chromosome-, arm-level, and focal CNAs (2,3). Certain CNAs are preferentially associated with specific cancer types and lineages (2,4–6), raising critical questions regarding the mechanisms underlying the selection of these tumor-specific CNAs and their functional consequences. While chromosome gains and losses typically reduce cellular fitness, cancer cells can tolerate such alterations, although the mechanisms enabling this adaptation remain largely unknown (7–9).

The tumor suppressor adenomatous polyposis coli (APC) is well recognized for its role in the WNT/β-catenin pathway as a key component of the β-catenin destruction complex. Its primary function is to regulate β-catenin levels by promoting its degradation (10–12). Loss of APC function leads to aberrant nuclear accumulation of β-catenin, hyperactivation of Wnt signaling, and uncontrolled proliferation, thereby driving tumorigenesis (10–12).

Beyond its role in Wnt signaling, APC is also a critical regulator of mitosis, and its dysfunction induces CIN (13–16). APC inactivation in various cell lines and mouse embryonic stem (ES) cells often results in chromosome mis-segregation due to premature mitotic exit (13–15,17). The CIN phenotype of *APC*-mutant cells is characterized by abnormal chromosome numbers and inhibition of apoptosis in both mouse and human tumors (18,19). However, the molecular mechanisms underlying APC-driven CIN remain controversial due to the complexity of its interactions with multiple mitotic regulators (20). Moreover, it is largely unknown whether and how CIN resulting from APC mutations influences tumor progression, potentially through the selection of specific CNAs.

Here, through a comparative analysis of APC function in cultured cells and tumors, we elucidated a functional relationship between APC deficiency and tumor-specific CNAs. First, we examined the global impact of APC deficiency on mitotic regulation in cultured cells. Second, we investigated the correlation between *APC* mutations and chromosomal abnormalities in mouse tumors and human cancers. Finally, we analyzed the process by which *APC* mutations lead to chromosomal reorganization and subsequent changes in cell fitness, allowing cells to mitigate the proliferative stress associated with aneuploidy. These findings provide new perspectives on how CNAs are selected in *APC*-mutant cancers.

## Results

### APC deficiency impairs centrosome maturation

APC cooperates with various mitotic regulators, including spindle assembly checkpoint proteins (Bub1/R1) and microtubule end-binding (EB) family proteins, to ensure accurate chromosome segregation (13–15) (Fig. S1A). Notably, detailed observations of HeLa cells have shown that APC depletion does not significantly disrupt mitotic spindle formation but leads to chromosome misalignment due to premature mitotic exit (15). The precise mechanism underlying this process remains unclear.

To detect subtle phenotypic differences in spindle architecture caused by APC depletion in HeLa cells, we employed lattice light-sheet microscopy (21) combined with GFP-fused EB1 (EB1-GFP), a marker of newly assembled microtubule ends (22). This approach enabled three-dimensional (3D) imaging of entire spindles with high spatiotemporal resolution (23) (Fig. 1A, B; Fig. S2A, B; and Movie 1–3). Following APC depletion, microtubule growth probability increased predominantly in the outer equatorial plane of the spindle, while it significantly decreased around centrosomes (Fig. 1C). These findings indicate that the primary effect of APC depletion is centrosome immaturity.

**Fig. 1.**
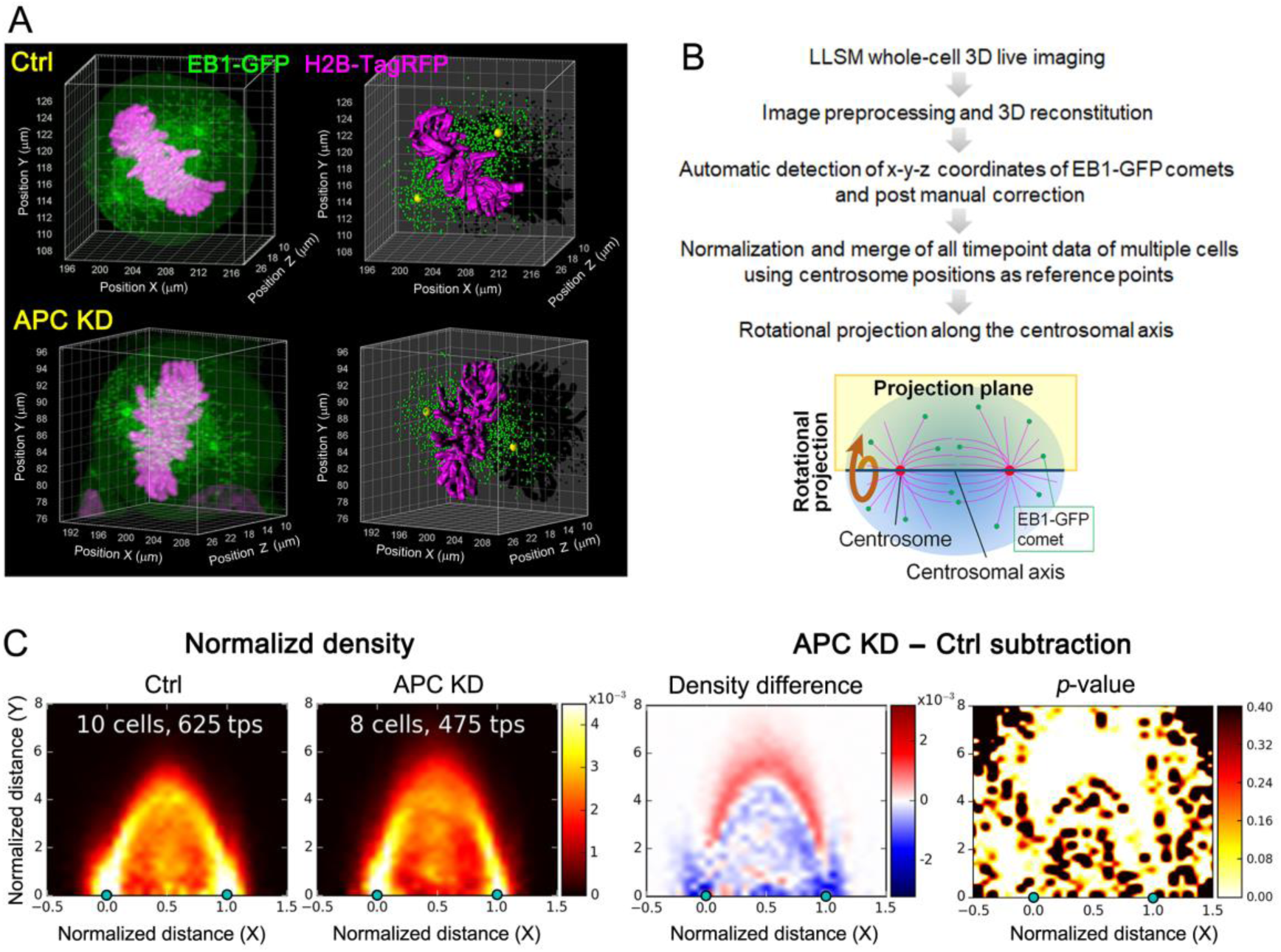
Identification of the site affected by APC depletion. (A) Three-dimensional representation of control (Ctrl) and APC-depleted (APC KD) mitotic HeLa cells expressing EB1-GFP and H2B-TagRFP as visualized by lattice light sheet microscopy. Original images (left) and examples of automated detection of EB1-GFP positions and chromosome surfaces (right) are shown. See also Fig. S2A and Movies S1–S3. (B) Procedure of quantitative analysis of the timelapse 3D data. The rotational projection method is shown diagrammatically in the bottom. (C) Rotationally projected density map of control and APC-depleted cells (left). The centrosome positions are indicated by green dots at positions 0.0 and 1.0 on the x-axis. The number of cells and time points (tps) in time lapse videos used are indicated in the panel. Density difference map (middle), and p-value map (right) are shown.

Consistent with this finding, γ-tubulin accumulation at centrosomes and the activation of the central mitotic kinases AURKA (autophosphorylated at T288) and polo-like kinase 1 (PLK1; phosphorylated at T210), which cooperatively regulate centrosome maturation (24,25), were significantly reduced in APC-depleted cells (Fig. 2A).

**Fig. 2.**
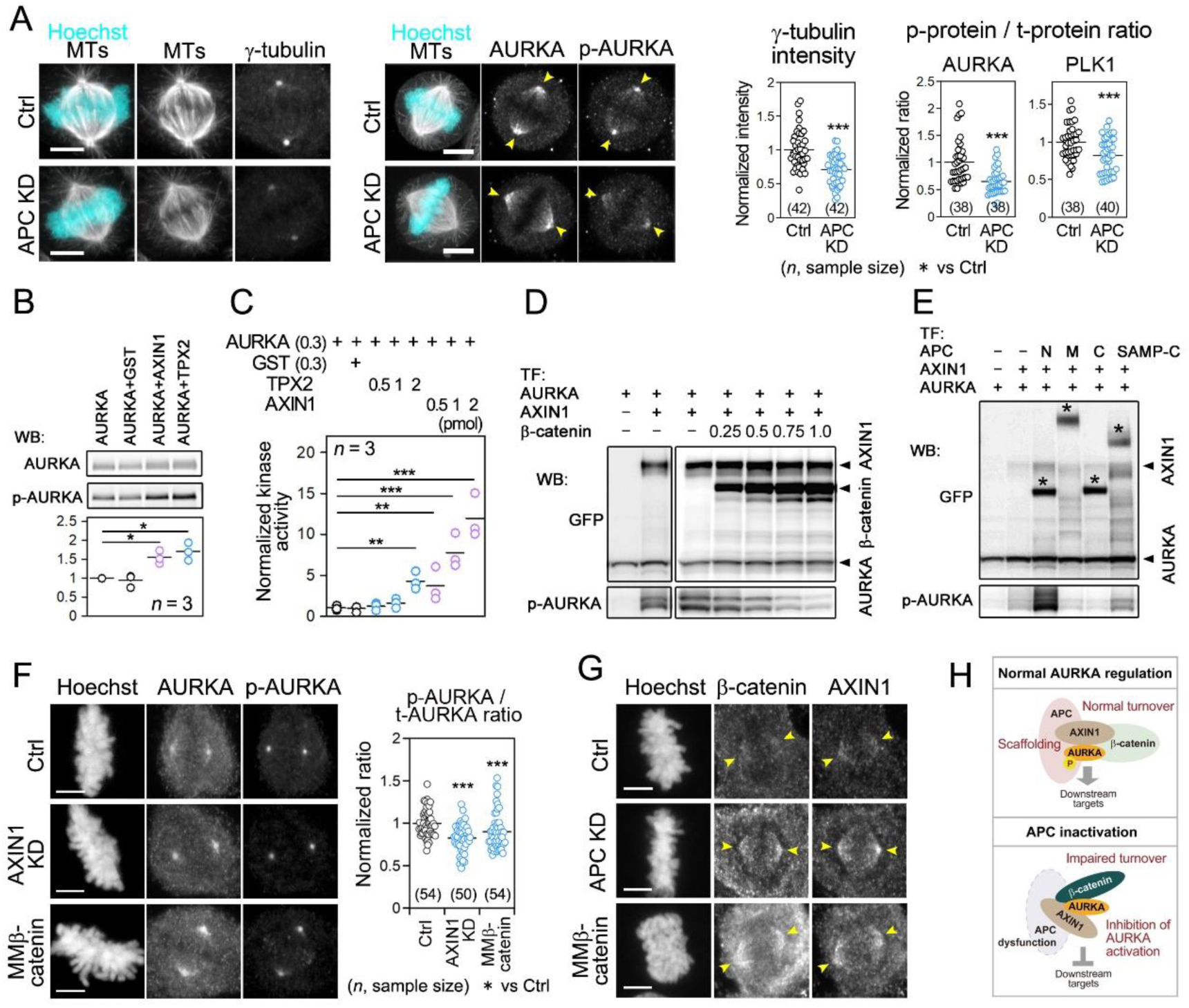
Regulation of AURKA by APC, AXIN1 and β-catenin. (**A**) Immunostaining of γ-tubulin, α-tubulin, AURKA and autophosphorylated AURKA (p-AURKA) with Hoechst 33342 to stain DNA (left). Arrowheads indicate spindle poles. Thereafter, see Table S1 for all representative images. Fluorescence intensities of γ-tubulin at centrosomes and ratios of fluorescence intensities (phosphorylated vs. total protein) of AURKA (T288) and PLK1 (T210) are shown (right). (**B**) In vitro AURKA autophosphorylation at T288 detected by western blotting. (**C**) In vitro kinase assay of AURKA autophosphorylation. (**D, E**) HEK293T cells were transfected with GFP-fused AURKA, AXIN1, β-catenin, and APC fragments (see Fig. S3A) as indicated and then subjected to western blotting using the indicated antibodies. APC bands are marked with asterisks. Note that the phosphorylated AURKA band always appeared as multiple bands. AXIN1 and AURKA levels were increased when coexpressed with binding APC fragments and AXIN1, respectively, probably due to stabilizing effect. Thereafter, see Table S2 for uncropped versions of blots. (**F**) AURKA status in AXIN1 depleted- and MMβ-catenin-expressing HeLa cells. (**G**) Distribution of β-catenin and AXIN1 in APC depleted- and MM-catenin-expressing HeLa cells. Arrowheads indicate spindle poles. (**H**) APC positively regulate AURKA together with AXIN1 and β-catenin (up). When APC is dysfunctional, its ability to activate AURKA reduced and stabilized β-catenin inhibits normal AURKA behavior and its activation (bottom). Scale bars: 5 μm. * *P* < 0.05;** *P* < 0.01; *** *P* < 0.001 (Student’s t-test in **A**, Tukey–Kramer-test in **B, C, F**).

### APC together with AXIN1 and β-catenin regulate AURKA activity

To elucidate the molecular mechanism by which APC regulates centrosomes, we screened centrosomal proteins involved in centrosome maturation for their association with APC. APC was found to bind AURKA, PCNT (pericentrin), and ch-TOG through its N- and/or C-terminal regions (Fig. S3). Since AURKA functions upstream of these interactors in centrosome maturation (26,27), we further examined APC’s role in AURKA regulation.

We did not detect direct promotion of AURKA activation by APC in vitro. However, given that β-catenin and AXIN1, a core scaffold protein in the β-catenin destruction complex (10), bind centrosomal proteins such as AURKA and PLK1 and regulate mitosis (28–30), and that Drosophila APC2 and Axin cooperate to promote mitotic fidelity (31), we investigated whether AXIN1 and β-catenin influence AURKA activation. In vitro and HEK293T cell-based assays revealed that the middle portion of AXIN1 promoted AURKA activation, similar to TPX2, a well-characterized AURKA activator (32) (Fig. 2B, C; Fig. S4A, B). Elevated β-catenin levels inhibited AXIN1-mediated AURKA activation (Fig. 2D), consistent with the competitive interaction of AURKA and β-catenin with AXIN1 (33). Furthermore, the N-terminal region of APC enhanced AXIN1-mediated AURKA activation, suggesting that APC facilitates scaffold formation between AXIN1 and AURKA (Fig. 2E; Fig. S3A, B).

Technical limitations prevented us from expressing a full-length APC fragment lacking only the C-terminal region. Instead, we utilized mouse embryonic fibroblasts derived from *Apc^1638T^* mice, in which the C-terminal half of APC is deleted (Fig. S1B) (34). The truncated APC protein in these cells lacks two of the three AXIN-binding domains and the C-terminal region that interacts with AURKA. These fibroblasts exhibited reduced total AURKA levels and a lower p/T-AURKA ratio, indicating that the APC C-terminal region also contributes to AURKA activation (Fig. S1B; Fig. S4C, D).

Consistently, in HeLa cells, AXIN1 depletion and expression of a dominant stable form of β-catenin (MMβ-catenin), in which multiple GSK phosphorylation sites required for degradation were mutated (10), reduced the active AURKA ratio (Fig. 2F; Fig. S2C). Stabilized β-catenin accumulated abnormally around centrosomes along with AXIN1, suggesting disrupted turnover of these molecules (Fig. 2G; Fig. S2C). These findings indicate that under normal conditions, APC and AXIN1 function as positive regulators of AURKA (Fig. 2H). Conversely, APC inactivation impairs AURKA activation, at least partially through β-catenin stabilization. However, since β-catenin and AXIN1 interact with multiple centrosomal factors, β-catenin stabilization likely has broader effects on the dynamics of these interacting factors.

### Upregulation of AURKA in *APC*-mutant tumors

The above results seem to contradict the traditional view that AURKA, which promotes cell cycle progression at centrosomes, is hyperactive in various tumor types and functions as an oncogene driving tumor growth (27). To examine AURKA activation in *APC*-mutant tumors, we analyzed *Apc^Δ716^*mice, which carry a heterozygous *Apc* truncation mutation leading to severe polyposis after loss of the wild-type allele, mimicking early tumor formation in humans (35) (Fig. S1B). Comparison of multiple polyps and crypts revealed that, unexpectedly, the active AURKA-to-total AURKA ratio was higher in many polyps than in crypts (Fig. 3A–C), despite the increase in β-catenin levels (Fig. 3C, E). Detailed analysis of individual polyps showed that active and total AURKA levels varied between polyps, exhibiting some degree of intra-tumor heterogeneity (Fig. 3C). Additionally, regions with exceptionally high AURKA expression were often observed within portions of the polyps (Fig. 3D). Consistently, in early-stage human *APC*-mutant tumors, increased AURKA activity was accompanied by enhanced proliferative activity and heterogeneity (Fig. S5). APC deficiency induces chromosomal instability (CIN) due to a combination of defects in mitosis and apoptosis (18). Given this, we considered whether AURKA-activated cells might arise as a consequence of copy number alterations (CNAs) involving AURKA-activating genes.

**Fig. 3.**
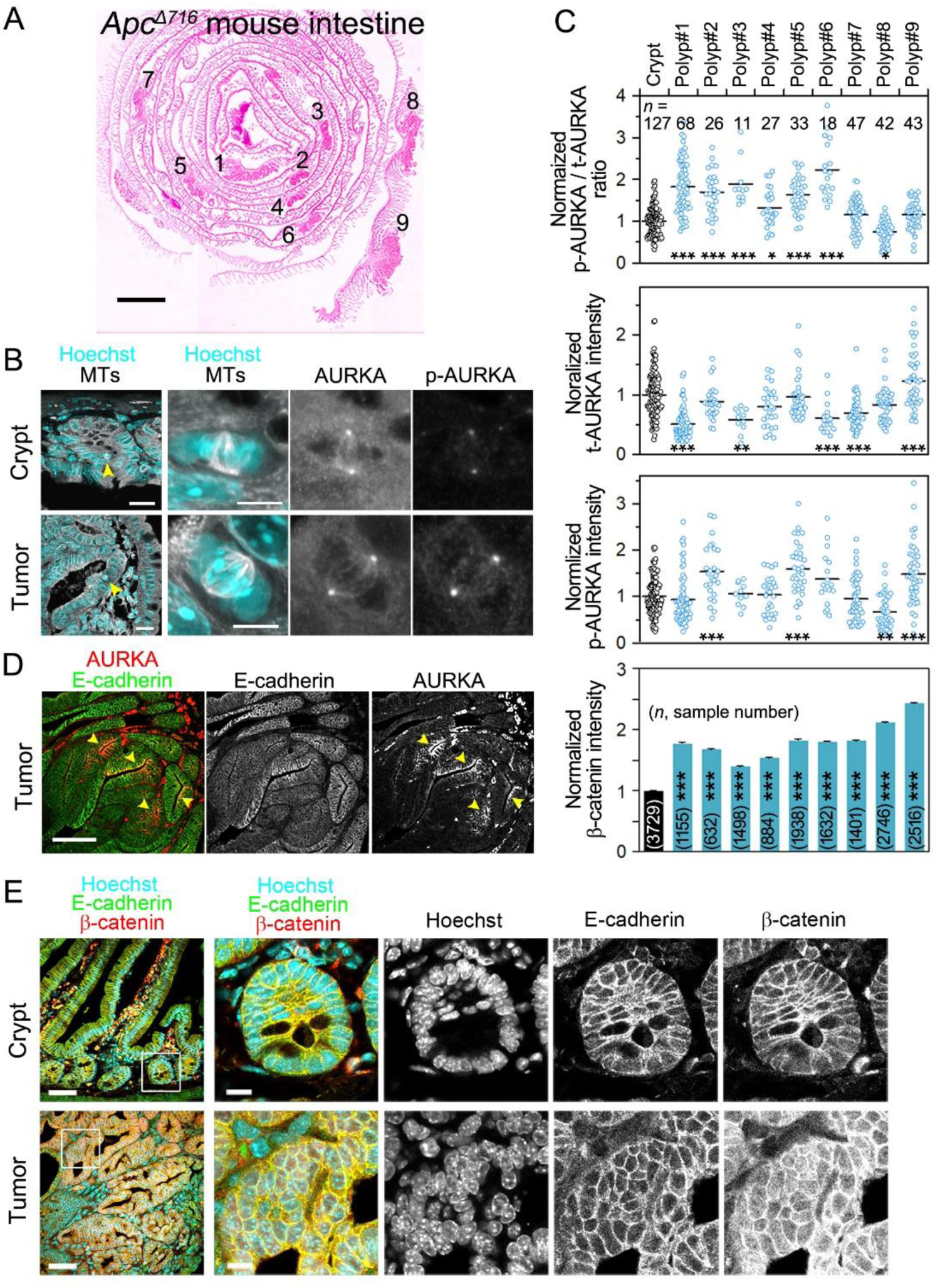
Upregulation of AURKA in mouse tumors. (**A**) HE-stained section of the intestines of a 16-weeks old *Apc^Δ716^* mouse. Polyps analyzed in this study are indicated by numbers. (**B**) Immunostaining of a formalin-fixed, paraffin-embedded *Apc^Δ716^* mouse intestine tissue section for microtubules (MTs), AURKA and p-AURKA with nuclear DNA stained by Hoechst 33342 to visualize crypt cells (top) and tumor cells (bottom) is shown. Low magnification images are shown on the left, and magnified images of mitotic cells indicated with arrowheads are shown in the three panels on the right. (**C**) Quantitative analysis of immunofluorescence signals for AURKA/p-AURKA and β-catenin in crypts and tumor tissues of nine polyps. (**D**, **E**) Immunostaining of a formalin-fixed, paraffin-embedded *Apc^Δ716^* mouse intestine section for AURKA, E-cadherin and β-catenin with nuclear DNA stained by Hoechst 33342 to visualize patchy region in tumors overexpressing AURKA (D) and crypt cells (top) and tumor cells (bottom) (E). In (E), Low magnification images are shown on the left and magnified images of the boxed areas are shown in the four panels on the right. Scale bars: 2 mm (A); 100 μm (D); 50 μm (E, low magnification, left), and 10 μm (E, high magnification, right). \**P* < 0.05; ** *P* < 0.01; *** *P* < 0.001 (Tukey–Kramer-test).

### Recurrent amplification of *AURKA* loci in human and mouse

To investigate the possibility of *AURKA* amplification, CNAs in human and mouse *APC*-mutant tumors were examined and compared. To investigate CNAs in *APC*-mutant human colorectal cancers, we interrogated The Cancer Genome Atlas (TCGA) dataset (3) for *APC* mutations and identified recurrent gain/loss of certain chromosomes in *APC*-mutant colorectal cancers (Fig. 4A). The differences in 20q, 13q and 7q gains and 18p, 18q, 14q, 4q, and 17p losses were significant, and there were also differences in 7p and 19q gains and 9q, 20p, and 1p losses in *APC*-mutant colorectal cancers. Although gains/losses of these chromosomes were also observed in cancers without detectable *APC* mutations, it is possible that APC expression is suppressed by methylation (36).

**Fig. 4.**
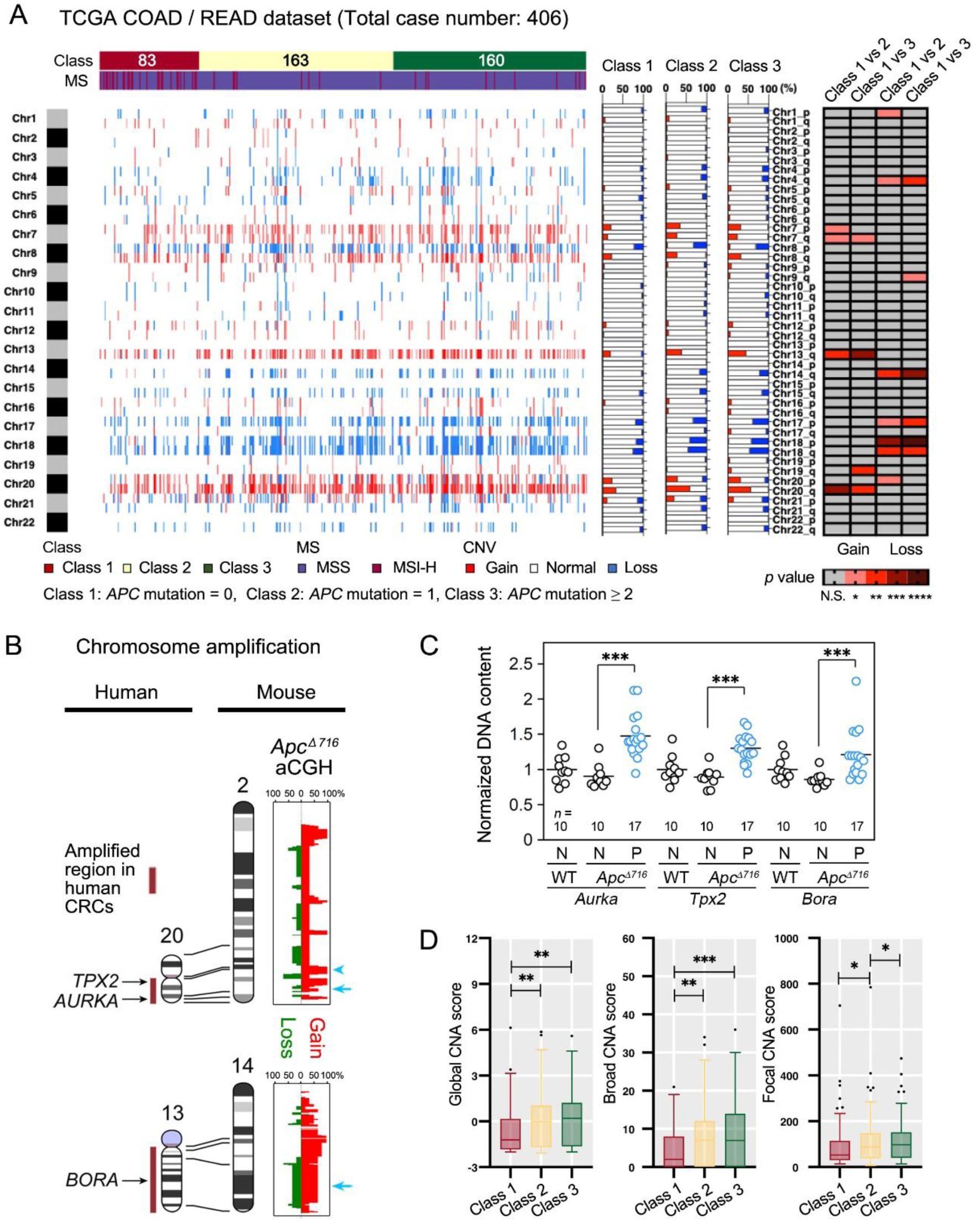
Recurrent CNAs in *APC*-mutant human colorectal cancers and mouse tumors. (**A**) The Cancer Genome Atlas (TCGA)-colon adenocarcinomas (COAD) and rectum adenocarcinomas (READ) samples are arranged along the x-axis and ordered in accordance with their *APC* mutations groups (Class 1: *APC* mutation = 0; Class 2: *APC* mutation = 1; Class 3: *APC* mutation ≥2). The y-axis numbered with 1–22 denotes chromosome numbers. The intensity of red and blue matches the frequency of copy number gain and loss, respectively. The ratio of gain to loss for each class is displayed as a stacked graph with the *p*-value of the Fisher’s exact test for each class on the far right. * *P* < 0.05; ** *P* < 0.01; *** *P* < 0.001; **** *P* < 0.0001. (**B**) Synteny alignment of human chromosomes 20 and 13, and mouse chromosomes 2 and 14, and aCGH analysis of *Apc^Δ716^* mouse polyps shown as frequency plots. Red and green indicate regions of gain and loss, respectively. Positions of mouse *Tpx2* (arrowhead, top), *Aurka* (arrow, top), and *Bora* (arrow, bottom) are shown. The three examined polyps all exhibited amplification of *Tpx2*. Amplification of *Aurka* and *Bora* loci was found in two polyps. See also Fig. S6. (**C**) Relative DNA content of *AURKA*, *TPX2*, and *BORA* genes in normal tissues (N) of WT and *Apc^Δ716^*mice and polyps (P) of *Apc^Δ716^* mice. Seventeen polyps from four *Apc^Δ716^* mice were analysed. (**D**) Global CNA score (GCS), broad CNA score (BCS), and focal CNA score (FCS) distribution by each class obtained from TCGA dataset analysis shown in (A). Wilcoxon rank-sum test significance is shown as * *P* < 0.05; ** *P* < 0.01; *** *P* < 0.001 (Tukey–Kramer-test in **C**, Wilcoxon rank-sum test in **D**).

We also performed high resolution array-based comparative genomic hybridization (aCGH) analysis to detect CNAs in individual polyps from *Apc^Δ716^* mice (Fig. S6). The structures of human and mouse chromosomes are different; therefore, we examined the gain or loss regions of chromosomes that overlap between the two, and found that human chromosome 20 and part of mouse chromosome 2, and human chromosome 13 and part of mouse chromosome 14 are commonly gained (Fig. 4B and Fig. S6). It is noteworthy that human chromosome 20q contains *AURKA* and *TPX2* at 20q13.2 and 20q11, respectively, and chromosome 13q contains *BORA* at 13q21.33, which encodes another AURKA-activating factor (37). Furthermore, qPCR analysis of the relative DNA content of *Aurka*, *Tpx2*, and *Bora* in multiple polyps from *Apc^Δ716^*mice showed that these genes were frequently increased in copy number (Fig. 4C). The syntenic comparison of human and mouse chromosomes, which shows cross-species overlap in the amplified regions, indicates that these AURKA-related loci were selected by functional relatedness rather than structural factors.

### CNAs induced by CIN in *APC*-mutant tumors

In the TCGA dataset analysis, global CNA scores, which provide an overall assessment of the CNA burden, were significantly higher in colorectal cancers with *APC* mutations than in colorectal cancers without *APC* mutations (Fig. 4D). In particular, in *APC*-mutant colorectal cancers, the frequency of broad CNAs, which are defined as whole-chromosome-level alterations, was markedly increased compared with focal CNAs, which are alterations of limited size that range from a few kilobases to part of a chromosome arm (38). Furthermore, the most frequently altered chromosomes showed a high frequency of the microsatellite stable phenotype, which is characterized by CIN (Fig. S7A).

It is important to note that the CNA scores for cases with *CTNNB1* (*β-catenin*) mutations were significantly lower than for cases without *CTNNB1* mutations (Fig. S7B) and no distinctive recurrent CNA pattern similar to those seen for *APC* mutations was observed (Fig. S7C), indicating that the effects of *APC* and *CTNNB1* mutations on CNA formation are distinct.

### APC-deficient MCF10A undergo tetraploidisation and maintain chromosome 20 trisomy

To investigate the effect of human chromosome 20 amplification, we used MCF10A, a nearly diploid, noncancerous p53-proficient human mammary epithelial cell line that spontaneously develops chromosome 20 trisomy (39,40) (Fig. S8A and Table S3). We isolated MCF10A subclones with chromosome 20 disomy [MCF10A(20di)] or trisomy [MCF10A(20tri)] (Table S3). Contrary to the expectation that AURKA amplification promotes cell proliferation, MCF10A(20tri) cells exhibited a longer doubling time than MCF10A(20di) cells (Fig. 5A, B and Movie S4). This is consistent with the observation that the gain of an extra chromosome often leads to poor proliferation due to aneuploidy-induced fitness deficits (41). Although AURKA levels at centrosomes were higher in MCF10A(20tri) cells than in MCF10A(20di) cells as expected, the proportion of activated AURKA was lower in MCF10A(20tri) cells, indicating that chromosome-level amplification of AURKA alone is insufficient for enhanced proliferation (Fig. 5C, D). This suggests that additional regulatory mechanisms are required to balance AURKA activity for optimal cell fitness.

**Fig. 5.**
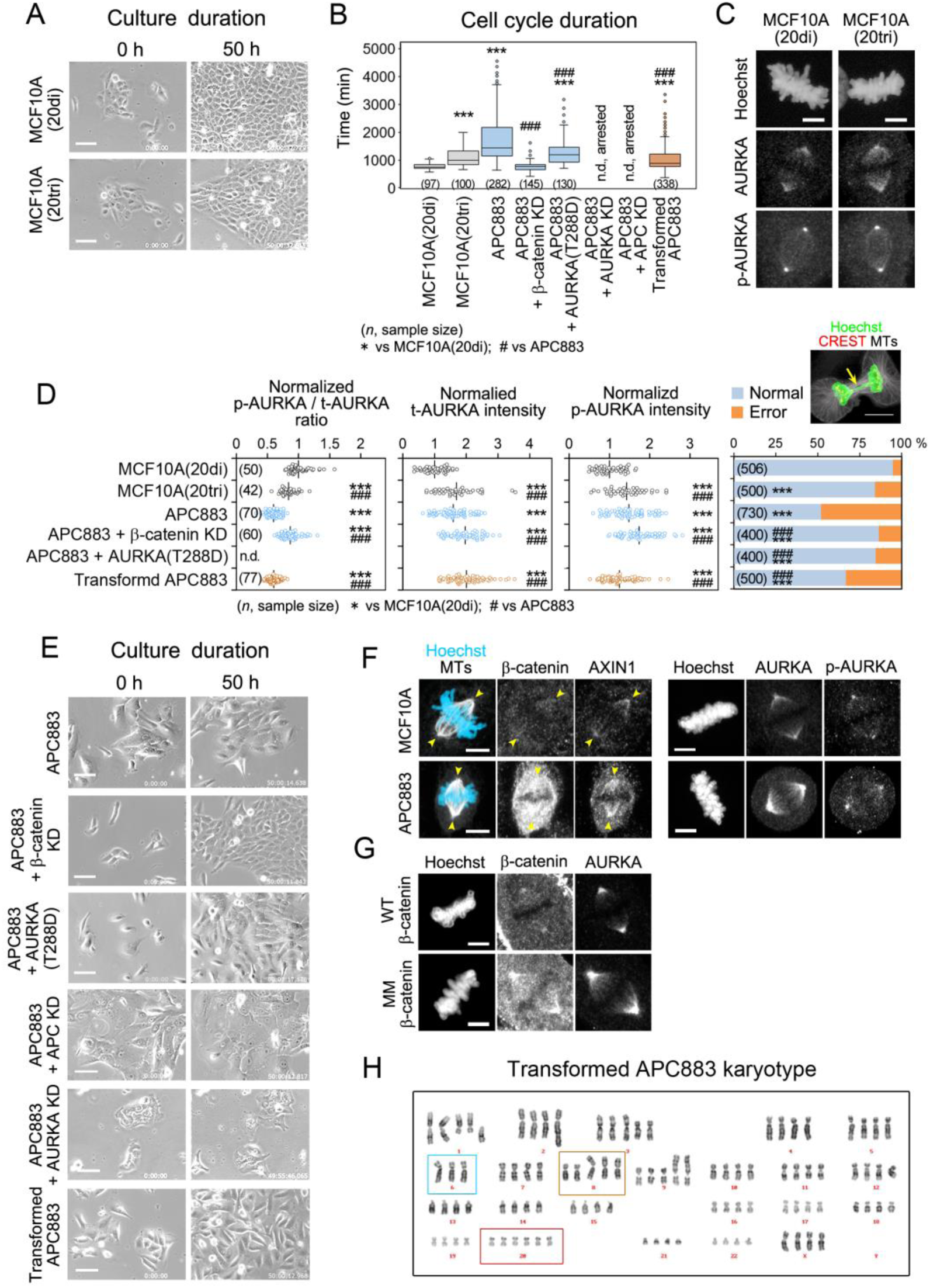
Proliferation and AURKA activities of MCF10A derived cell lines. (**A**) Selected images from phase contrast timelapse videos of MCF10A(20di) and MCF10A(20tri) cells. See also Fig. S8A and Movie S4, 5. (**B**) Doubling time of MCF10A- and APC883-derived cell lines. Note that no data are available for AURKA or APC depleted APC883 cells, as their proliferation is arrested. (**C**) Immunostaining of MCF10A(20di) and MCF10A(20tri) cells for AURKA and p-AURKA. (**D**) AURKA status at centrosomes obtained from immunofluorescence images. Total AURKA protein levels (t-AURKA), autophosphorylated AURKA levels (p-AURKA), the auto-phosphorylation ratio (p-AURKA/t-AURKA ratio), and mitotic error rate in MCF10A- and APC883-derived cell lines are shown as indicated. The values were normalized to 1 using MCF10A(20di) cells as a reference. An example of a chromosome separation error (arrow) is shown in the image on the right. (**E**) Selected images from phase contrast timelapse videos of the indicated cell lines. See also Movie S4, 5. (**F**) Immunostaining of MCF10A and APC883 cells as indicated. Position of spindle poles are indicated with arrow heads. β-catenin was up-regulated in the whole cytoplasm and at the spindle pole in APC883 cells, but only faintly detected on the spindles in MCF10A cells. In APC883 cells, AXIN1 and AURKA were also increased, whereas the proportion of p-AURKA is low. (**G**) Immunostaining of MCF10A expressing WTβ-catenin or MMβ-catenin for AURKA, showing accumulation of MMβ-catenin and AURKA around the centrosomes. (**H**) Representative karyotype of spontaneously transformed APC883 cells that contained a near tetraploid karyotype with six copies of chromosome 20 (red box) and other common numerical imbalances (cyan and brown boxes). Scale bars: 5 μm (C, F, G); 20 μm (A, E). *** *P* < 0.001; ^##^ *P* < 0.01; ^###^ *P* < 0.001 (Tukey–Kramer test).

Next, we analyzed APC883 cells, a MCF10A-derived cell line carrying a homozygous *APC* mutation leading to a short, truncated APC protein lacking all domains involved in β-catenin regulation (Fig. S1A, B). This cell line is near-diploid but consistently exhibits chromosome 20 trisomy (Fig. S2D, S8B and Table S3). APC883 cells displayed an even longer doubling time than MCF10A(20tri) cells (Fig. 5B, E and Movie S4). Notably, β-catenin accumulated around the centrosome with AURKA (Fig. 5F, G), and despite high AURKA levels at centrosomes, the proportion of activated AURKA was low (Fig. 5D). The requirement for AURKA activity in these cells was evident, as moderate knock-down of AURKA, at a level that did not inhibit MCF10A cell proliferation (Fig. S2F, Movie S5), led to immediate growth arrest. Similarly, knock-down of truncated APC, which retains AURKA-activating ability, also resulted in growth arrest (Fig. 5B, Fig. S2E, F and Movie S4). In contrast, the expression of active AURKA (T288D) or β-catenin knock-down, both of which increase AURKA activation, partially rescued proliferation and mitotic fidelity (Fig. 5B, D, E and Fig. S2D, Movie S4). Despite these interventions, chromosome 20 trisomy was maintained (Fig. S8C, D). These findings indicate that in APC883 cells, chromosome 20 trisomy alone is insufficient to support proliferation due to inadequate AURKA activation caused by high β-catenin levels. This results in heightened sensitivity to AURKA perturbations, necessitating the maintenance of chromosome 20 trisomy. Importantly, the effect of chromosome 20 amplification on proliferation differs between MCF10A and APC883 cells; chromosome 20 trisomy provides a fitness advantage only in the context of APC mutation.

Additionally, the expression of stable MMβ-catenin in MCF10A cells did not slow proliferation (Fig. S2D, F, Movie S5), indicating that functional APC counteracts the negative effects of β-catenin. This may explain why β-catenin-mutant human colorectal cancers (CRCs) do not exhibit a CIN phenotype or generate a CNA pattern similar to that seen with APC mutations (Fig. 4A, Fig. S7C).

### Chromosomal alterations in *APC*-mutant cells with extended passage

As APC883 cells were passaged over time, a subpopulation with an increased growth rate gradually emerged and became dominant in the culture (“transformed APC883,” Fig. 5B, D, E, and Movie S4). These cells exhibited newly acquired karyotypic abnormalities, including a near-tetraploid chromosome content with recurrent numerical imbalances (Fig. 5H; Fig. S8E and Table S3). Notably, chromosome 20 was present at the highest copy number (six copies per cell) in transformed APC883 cells (Fig. 5H; Fig. S8E). This coincided with an increase in total AURKA levels, but the proportion of activated AURKA remained low compared to MCF10A(20di) cells, indicating that AURKA activation alone does not drive increased proliferation (Fig. 5D). These findings suggest that an increase in AURKA levels or activity alone is insufficient to fully restore proliferation. Instead, maintaining a balanced chromosomal composition is necessary to support the growth of *APC*-mutant cells, highlighting the importance of chromosomal alterations in adapting to fitness impairments. However, despite these chromosomal changes, the proliferation rate of transformed APC883 cells remained lower than that of MCF10A(20di) cells (Fig. 5B). Instead of directly enhancing proliferation, these chromosomal changes may serve as an adaptive mechanism that enables *APC*-mutant cells to survive and continue growing under selective pressure.

### Increased sensitivity of *APC*-mutant tumor cells to AURKA inhibition

Finally, we confirmed the necessity of AURKA activity for tumor growth in *APC*-mutant tumors by inhibiting AURKA. In *Apc^Δ716^* mice treated with the AURKA inhibitor alisertib (MLN8237), the proportion of Ki67-positive tumor cells, a marker of proliferation, was reduced, whereas Ki67 levels in crypts remained unchanged (Fig. 6A). These results demonstrate that *APC*-mutant tumor cells are more dependent on AURKA activity, consistent with observations in cultured cells. Polyp counting further showed that AURKA inhibition reduced the total number of polyps (Fig. 6B), indicating that AURKA inhibition slows tumor growth rather than inducing complete regression. Together with our findings in MCF10A cells, these results suggest that AURKA activity is a necessary but not sufficient factor for sustaining proliferation in *APC*-mutant cells, and that additional chromosomal alterations may play a role in tumor adaptation.

**Fig. 6.**
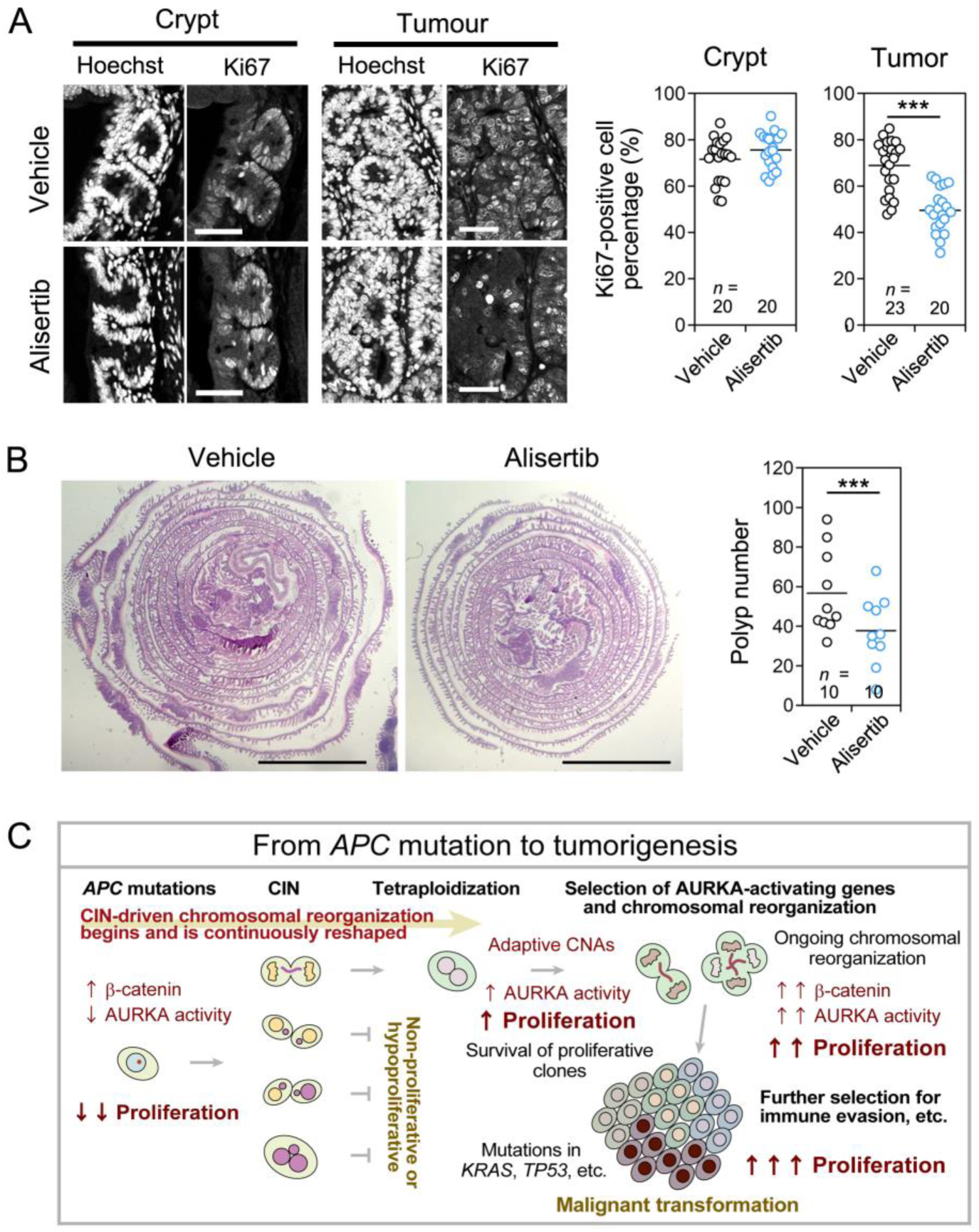
Requirement of AURKA activity for tumor cell growth. (**A**) Effect of AURKA inhibitor on the proliferative activity of mouse intestinal tissues. *Apc^Δ716^* mice were treated with the vehicle or AURKA inhibitor alisertib (20 mg/kg) for 14 consecutive days. Representative images of Ki67-labelled tissue sections from *Apc^Δ716^* mice (left). Analysis of the Ki67-positive cell population (right). (**B**) Effect of AURKA inhibitor on polyp development in *Apc^Δ716^*mice treated with the vehicle or AURKA inhibitor for 28 consecutive days. Examples of HE-stained sections of intestines from vehicle- or alisertib-treated *Apc^Δ716^* mice (left). Analysis of the polyp number and size distribution in the intestines of vehicle- or alisertib-treated *Apc^Δ716^* mice (right). Scale bars: 50 μm (A); 5 mm (B). \*\*\**P* < 0.001 (Student’s t-test). (**C**) Model showing the steps of malignant transformation during early tumorigenesis. *APC* mutations induce CIN partly due to AURKA inactivation and growth retardation. Some cells improve fitness via AURKA-activating CNAs, adapting to promote tumor growth, which may contribute to the formation of cancer-specific CNAs in *APC*-mutant tumors.

Tetraploid or near-tetraploid cells have been observed in early-stage cancers, including *APC*-mutant tumors (18,19,42,43), and are thought to serve as a transitional state during the acquisition of CNAs (43,44). Consistent with this, our data show that near-tetraploid *APC*-mutant MCF10A cells exhibited improved proliferation compared to their parental counterparts, whereas amplification of only chromosome 20 in parental MCF10A cells was associated with reduced fitness. These findings support a model in which tetraploidization facilitates the emergence of complex CNAs, including those that influence AURKA activity (Fig. 6C).

## Discussion

Cells must maintain a proper chromosomal composition to sustain proliferative fitness, yet *APC*-mutant cells take advantage of chromosomal imbalances to acquire adaptive fitness for tumor formation. APC normally cooperates with AXIN1 to enhance AURKA activation; however, when APC is dysfunctional, stabilized β-catenin plays a major role in inhibiting AURKA function. In *APC*-mutant cells, as well as in cultured cells expressing stable mutant MMβ-catenin, β-catenin accumulated near the centrosome. Phosphorylated β-catenin is rapidly degraded (10–12); therefore, stable β-catenin is no longer removed from the centrosome and may persistently block AURKA. Our findings suggest that this sustained β-catenin accumulation disrupts mitotic fidelity and cell cycle progression, in contrast to its well-established role in promoting proliferation and differentiation during normal development. However, in the presence of functional APC, the effects of MMβ-catenin expression were minimal, indicating that APC also regulates β-catenin through mechanisms beyond phosphorylation-mediated degradation. This additional level of regulation may underlie the distinct CNA patterns observed in *APC*-mutant versus *β-catenin/CTNNB1*-mutant cancers. Further research is required to elucidate APC’s role in pathways beyond WNT/β-catenin signaling.

The mechanisms driving distinct CNA patterns across different cell and cancer types remain largely unknown. By reanalyzing the TCGA dataset, we identified a strong correlation between *APC* mutations and specific chromosomal gains and losses, demonstrating how a single molecular alteration can shape CNA formation. Notably, syntenic comparisons between mice and humans, despite their differing chromosomal architectures, revealed shared amplification loci in *APC*-mutant contexts. This suggests that *APC* mutations drive CNA selection through functional cellular changes rather than through structural constraints imposed by chromosome organization. Indeed, our experimental data indicate that CNA selection in *APC*-mutant cells is linked to the restoration of AURKA activity. These findings highlight a mechanistic basis for the correlations observed in TCGA analyses, suggesting that specific CNAs provide a selective advantage by compensating for the proliferation deficits caused by *APC* loss.

*APC* mutations negatively impact proliferation, necessitating compensatory mechanisms for tumor cell survival. Our findings indicate that *APC*-mutant cells experience strong selection pressure for CNAs that restore AURKA activity, as insufficient AURKA function is detrimental to their viability. Since AURKA plays a central role in mitotic fidelity and cell cycle progression, its loss reduces cellular fitness, creating a disadvantage for *APC*-mutant cells. However, our results also show that increasing AURKA activity alone is insufficient to fully restore proliferation. Instead, maintaining a balanced chromosomal composition is required for optimal cell fitness in *APC*-mutant cells.

Although the selection for CNAs that enhance AURKA function was observed in both our in vitro culture model and in vivo *APC*-mutant tumors, the specific CNA patterns differed. This discrepancy may be attributed to the fact that MCF10A cells were originally established with a stable but imbalanced karyotype (39,45), making it unsurprising that different chromosomal patterns emerged. Nonetheless, even in MCF10A cells, chromosome 20 was never lost following *APC* mutation, suggesting a strong selective pressure to retain this chromosome. Importantly, amplification of the *AURKA* locus was consistently observed across MCF10A cells, mouse tumors, and human cancers, indicating that in the context of *APC* mutations, *AURKA* amplification represents a common adaptive strategy to improve cellular fitness, which may subsequently contribute to tumor progression. MCF10A cells, unlike colorectal cancer cell lines, allow us to isolate the effects of *APC* mutations on CNA selection without the confounding influences of pre-existing genomic instability. This enables us to extend beyond intestinal tumorigenesis, potentially contributing to a broader understanding of how APC loss influences chromosomal alterations in cancer.

Moreover, although reducing β-catenin can enhance proliferation in vitro, *β-catenin/CTNNB1* mutations are rare in *APC*-mutant tumors, likely because β-catenin remains essential for tumorigenesis (46,47). We observed that AURKA activation not only improves proliferative capacity but also enables β-catenin to exert its tumorigenic effects, demonstrating that both factors must act in concert to drive tumor progression. Thus, CNAs that enhance AURKA function not only mitigate the disadvantages caused by *APC* mutations but also facilitate β-catenin-dependent tumor evolution.

Our TCGA analysis also revealed significant correlations between *APC* mutations and chromosomal losses, particularly on chromosomes 18, 14, 4, and 17—alterations frequently detected in early colorectal cancer (4). These chromosomal deletions may contribute to tumorigenesis through yet unidentified pathways or by altering gene dosage and regulatory networks, potentially activating key proliferative factors such as AURKA. Mechanisms underlying AURKA activation could involve transcriptional upregulation, RNA stabilization, or other post-transcriptional modifications. Alternatively, these deletions may be selected in response to the additional selective pressures required for tumor growth in vivo, such as immune evasion (48,49) and hypoxic adaptation (50). Our findings demonstrate the necessity of CNAs that enhance AURKA activation in *APC*-mutant cells, but further investigation is required to elucidate how other selective pressures contribute to the acquisition of tumor type- or context-dependent CNA patterns in tumor evolution.

Although CNAs are common in cancers, they typically impose a fitness burden in normal cells (51). While it has long been unclear why tumor cells tolerate such fitness-impairing CNAs, our findings reveal that *APC* mutations shape CNA selection by influencing cellular fitness. We found that APC-mutant cells display distinct aneuploidy preferences compared with APC-active cells. In APC-active MCF10A cells, chromosome 20 amplification slowed proliferation, whereas APC-mutant MCF10A cells overcame their proliferative disadvantage by maintaining chromosome 20 while transitioning toward near-tetraploidy. While the specific CNA patterns observed in MCF10A cells differ from those in human tumors, MCF10A remains a valuable model for experimentally dissecting the selective pressures that drive chromosome 20 amplification and polyploidization. Despite differences in overall CNA patterns, the transition toward near-tetraploidy observed in *APC*-mutant MCF10A cells suggests a conserved mechanism by which polyploidization mitigates the fitness costs of chromosomal instability. Polyploidy, frequently observed in early-stage cancers (42,52,53), has been proposed to buffer against fitness defects arising from chromosomal imbalances (8,54). However, polyploid cells are inherently unstable and often serve as an intermediate state leading to aneuploidy and ultimately tumor progression (43,44). Our observations in *APC*-mutant cells support the idea that polyploidization mitigates the fitness costs of chromosomal instability, thereby promoting tumor evolution.

In summary, our findings suggest a functional relationship between *APC* mutations, tumor-promoting CNAs, and adaptive fitness. *APC* mutations initially reduce AURKA activity and proliferative potential, but *APC*-mutant cells adapt by exploiting compensatory CNAs that arise from mitotic errors, recovering AURKA activity and facilitating fitness adaptation during tumor evolution. This underscores how genetic background influences CNA formation and enables tumor cells to acquire fitness traits that support their progression. While AURKA remains a promising therapeutic target, *APC*-mutant cells’ ability to acquire CNAs suggests they may rapidly develop drug resistance. Thus, rather than targeting AURKA, effective colorectal cancer treatments should focus on preventing chromosomal rearrangements to constrain tumor adaptation and maintain therapeutic efficacy.

## Materials and Methods

### Cell culture and treatment

All cell lines tested negative for mycoplasma contamination. Parental HeLa cell line was authenticated by Bio-Synthesis Inc (Cell Culture STR Profiling and Comparison Analysis (Cat. No.:CL1003). Other cell lines used were obtained from reliable distributors. HeLa cells and HeLa cells expressing EB1-GFP and H2B-TagRFP (clone A1) were maintained as previously described (55,56). A parental MCF10A cell line and an MCF10A cell line with the *APC* gene knocked out (MCF10A cells APC^−/−^) were obtained from Sigma-Aldrich-Aldrich (CLLS1069, St. Louis, MO, USA), and cultured in as described previously (57). The APC^−/−^ cells, which were generated by zinc finger nuclease (ZFN) technology, are homozygous for a 17 bp deletion (GCAGCCCAGATTGCCAA) in the endogenous *APC* gene at 2650-2666 bp of transcript variant 3 (NM_000038). This mutation results in a truncated 883-amino acid APC protein followed by an unrelated 21 amino acid sequence at its C-terminus (SHGRSVSHSYLSGRQKFWVYH). In this paper we refer to this cell line as APC883. The HEK293-based Lenti-X 293T (HEK293T) cell line was obtained from Takara Bio (Shiga, Japan) and was cultured in DMEM (10566024, Thermo Fisher Scientific/Gibco, Waltham, MA, USA) supplemented with 10% FCS at 37°C in a 5% CO_2_ atmosphere. Colorectal cancer cell lines SW480 (CCL-228), HT-29 (HTB-38), and HCT116 (CCL-247) were obtained from ATCC (Manassas, VA, USA). SW480 was cultured in Leibovitz’s L-15 Medium (11415064, Thermo Fisher Scientific/Gibco) supplemented with 10% FCS at 37°C without CO_2_. HT-29 and HCT116 were cultured in McCoy’s 5a Medium (16600082, Thermo Fisher Scientific/Gibco) supplemented with 10% FCS at 37°C in a 5% CO_2_ atmosphere. DLD-1 (JCRB9094) was obtained from JCRB Cell Bank (Osaka, Japan) and cultured in RPMI1640 medium (11875093, Thermo Fisher Scientific/Gibco) with 10% FCS at 37°C in a 5% CO_2_ atmosphere. Mouse embryonic fibroblasts (MEFs) of wild-type C57BL/6 mice and homozygous *Apc^1638T^* mice were isolated from embryonic day (E) 13.5 embryos, and were cultured in DMEM supplemented with 10% FCS in a 5% CO_2_ atmosphere. The number of cells was counted using an automated cell counter (Countess; Invitrogen, Carlsbad, CA, USA).

### Lattice light sheet microscopy and data analysis

Three-dimensional live imaging by lattice light sheet microscopy and quantitative data analysis were performed as described previously (21,23). Briefly, timelapse images of HeLa cells that stably expressed EB1-GFP and H2B-TagRFP (clone A1) (56) were collected at 0.755-s intervals for EB1-GFP only single channel imaging, 1.550-s intervals for EB1-GFP and H2B-TagRFP dual channel imaging in fast scanning mode (3D voxel pitch: 0.100 × 0.100 × 0.217 μm), or 10.000-s intervals for EB1-GFP and H2B-TagRFP dual channel imaging in high resolution mode (3D voxel pitch: 0.100 × 0.100 × 0.195 μm). The former data were used for quantitative analyses and the latter data were used to visualize mitosis progression. The data were displayed in 3D and processed using Imaris software (Bitplane, Zurich, Switzerland). Supplementary video editing of projected images exported from Imaris and image processing were performed using ImageJ software, QuickTime software (Apple. Cupertino, CA, USA), and Adobe Photoshop software (Adobe, San Jose, CA, USA).

To interpret multidimensional data, images were processed for visualization and mathematical computation. The XYZ coordinate positions of EB1-GFP comets were determined automatically. The results are presented as spots or density, or analysed mathematically using a program developed with MATLAB (MathWorks, Natick, MA, USA) as described previously (23) and Python using libraries NumPy, Math, and Matplotlib. An EB1-GFP comet density map (Fig. 1C) was created from the 2D rotational projected comet numbers and the volume associated with the 2D grid. To create a 2D rotational projection of all comets, the coordinates of comets and centrosomes were first transformed so that the spindle axis (vector that connected the two centrosomes) aligned with the x-axis of the Cartesian coordinate. Then, all comet points were rotated around the x-axis and projected onto an arbitrary 2D plane with its normal vector orthogonal to the spindle axis. The result was a half spindle that consisted of comets and centrosomes in 2D coordinates. Before assembling comet points from all cells, the x-coordinates of 2D comets were rescaled so that the inter-centrosomal distance was one. Then, a regular 2D grid that covered all comet spots was created and the number of comets in each grid element was counted. Average comet density of each element was calculated by normalizing the number of comets with i) the timelapse duration and ii) calculation of the 3D volume that corresponded to each grid element. The latter is a function of distance from the spindle axis and is expressed as *v*=π*h^2^w*(2*n*-1) where *n*=1,2,3,…,*Ny* with *Ny* that represents the total number of grid elements perpendicular to the spindle axis. *w* and *h* are the width and height of the elements, respectively. For cell type comparison, comet density was normalized so that the summed density of all elements was one. Normalized density was then rescaled by the global maximum to produce a value between zero and one. Rescaled normalized comet density of each element and interpolated density values of in-between grid elements were coded by color and plotted. Because the centrosomes had much higher densities relative to other regions, we imposed an upper cutoff color value to better illustrate the density variation of non-centrosomal regions. The density difference graph was constructed by the difference between the normalized density of each grid element of APC KD and control cells. A significance test of the density difference was performed by the two-sided t-test to obtain the *p*-value. The inputs for the test were the mean and variance value for each grid of the density map. The output is the *p*-value for each grid. The code was deposited in Github (https://github.com/wxchew/cometdensity/blob/master/2DProjectedCometDensity.ipynb).

### Antibodies and immunolabeling

No untested antibody was used. Rat anti-E-cadherin mAb (ECCD-2 (58)) was a kind gift from Dr. M. Takeichi (RIKEN CDB). The following commercial primary antibodies were used: mouse anti-APC (Ab-1) mAb for western blotting, clone FE9 (OP44; Lot# D00138185; EMD Millipore/Calbiochem, Billerica, MA, USA); mouse anti-Aurora A mAb, clone 35C1 (ab13824; Lot# GR121501-2; Abcam, Cambridge, UK); rabbit anti-Aurora A (phospho T288) polyclonal antibody (pAb), (ab195748, Lot# GR212301-1; Abcam); rabbit phospho-Aurora A (Thr288)/Aurora B (Thr232)/Aurora C (Thr198) (D13A11) mAb (#2914; Lot# 3; CST, Danvers, MA, USA); mouse anri-PLK1 mAb, clone 36-298 (ab17057; Lot# GR136555-1; Abcam); rabbit anti-PLK1 (phospho T210) mAb, clone EPNCIR167 (ab155095; Lot# YJ092901CS; Abcam); rabbit anti-AXIN1 mAb, clone C76H11 (#2087; Lot# 4; CST); purified human anti-Centromere Protein IgG (CREST) (15-235; Lot# 441.13BK.20; Antibodies Incorporated, Davis, CA, USA); rabbit anti-β-catenin pAb (C2206; Lot# 059K4754; Sigma-Aldrich); mouse anti-β-catenin mAb, clone 14 (610154; Lot# 5113978; BD Transduction Lab.); rabbit anti-Ki67 pAb (ab15580; Lot# GR264768-1; abcam); mouse anti-α-tubulin mAb, clone DM1A (T9026; Lot# 078K4763; Sigma-Aldrich); rat anti-tubulin mAb, clone YL1/2 (ab6160; Lot# GR57687-1; Abcam); rabbit anti-γ-tubulin pAb (ab11320; Lot# 628882; Abcam); mouse anti-γ-tubulin mAb, clone GTU-88 (T6557; Lot# 079K4861; Sigma-Aldrich); mouse anti-GAPDH antibody-loading control (HRP), clone mAbcam 9484 (ab9482; Lot# GR149747-1; Abcam); rabbit anti-GFP, HRP-conjugated (598-7; Lot# 002; MBL, Aichi, Japan); mouse anti-GST, clone B-14 (sc-138; Lot# K1808; Santa Cruz Biotechnology); mouse anti-penta-His antibody (34660; Lot# 136244018; Qiagen); mouse anti-Myc tag mAb, clone 4A6 (05-724; Lot# DAM1724025; Millipore); mouse anti-Myc tag mAb, clone 4A6, peroxidase-conjugated (16-213; Lot# DAM1646292; Millipore); Hoechst 33342 solution (346-07951; Lot# FN027; Dojin, Japan) was used as a nuclear marker. Immunofluorescent staining of fixed cultured cells and western blotting were performed as described previously (55–57). Intestinal tissues from 16-weeks old *Apc^Δ716^* mice and human adenomas were fixed in formalin and processed for paraffin-embedding and sectioning as described previously (35). Antigen retrieval for β-catenin (610154, BD), E-cadherin (ECCD-2), AURKA, phosphor-AURKA(T288), Ki67 and anti-α-tubulin (YL1/2) immunostaining was performed by boiling sections for 30 min in Tris-EDTA pH 9.0.

### cDNAs, plasmids and RNA interference

The APC cDNA used was described previously (59). Wild-type mouse β-catenin and dominant-stable β-catenin (MMβ-catenin), in which the NH_2_-terminal four Ser/Thr residues recognized by GSK-3β were substituted for Ala (60) were gifts from Dr. A. Nagafuchi (Nara Medical University). A human AXIN1 cDNA were PCR cloned from MCF10A cells. cDNAs for rat AXIN1 were described previously (12,61). The following plasmids were obtained: pWZL Neo Myr Flag AURKA (addgene plasmid # 20427), a gift from William Hahn & Jean Zhao; mEmerald-TPX2-N-10 (addgene plasmid #54285) and mEmerald-Beta-Catenin-20 (mouse) (addgene plasmid #54017), gifts from Michael Davidson. AURKA(T288D) mutation was introduced by PCR engineering. EGFP-fused APC, β-catenin, AXIN1 and AURKA were expressed by inserting these cDNAs into pEGFP-C (Takara/Clontech). Transient transfection of cells with plasmids was performed using Effectene transfection reagent (Qiagen, Germany).

Lentiviral vectors expressing genes of interest under EF1α promoters (pLVSIN-EF1a-IRES-pur, pLVSIN-EF1a-IRES-bsr) were described previously (56). Myc-tagged AURKA(T288D) and MMβ-catenin were generated by PCR engineering and inserted into pLVSIN-EF1a-IRES series vectors. Lentiviral vectors expressing shRNAs against human *APC* under U6 promoters (pLVSIN-U6-pur, bsr) were described previously (62). The target sequences of shRNAs for human *AURKA was* CAGGACCTGTTAAGGCTACAG. The lentiviral plasmids for shRNA against β-catenin pLKO.1 puro shRNA beta-catenin was obtained from addgene (plasmid #18803, a gift from Bob Weinberg).. The target sequences of siRNA for human AXIN1 were AAGCCAGCCACCAAGAGCTTCATAA (Stealth RNAi, invitrogen). Transfection of cells with siRNA was performed using HiPerFect Transfection Reagent (Qiagen).

### Lentiviral gene transfer

Lentiviruses that carried genes of interest were generated in HEK293T(Lenti-X) cells (Takara) using Lentiviral High Titer Packaging Mix (Takara) as described previously (56). Cells were infected for 24 h with the virus in the presence of 10 mg/ml polybrene (Santa Cruz Biotechnology), washed, and allowed to recover for 48 h before selection with appropriate antibiotics. For selection of gene-transduced cells, puromycin (InvivoGen, San Diego, CA, USA) and blasticidin S (InvivoGen) or bleocin (Wako) were used for selection because of the *bsr* resistance gene. HeLa cells transfected with cDNA and shRNA were maintained in medium that contained appropriate concentrations of selection drugs: APC shRNA/pLVSIN-U6-pur, 10–20 μg/ml puromycin; APC shRNA/pLVSIN-U6-bsr, 100 μg/ml blasticidin S; myc-MMβ-catenin/pLVSIN-EF1a-IRES bsr, 1 μg/ml blasticidin S. MCF10A cells transfected with cDNA and shRNA were maintained in medium that contained appropriate concentrations of selection drugs: APC883 + myc-AURKA(T288D)/pLVSIN-EF1a-IRES bsr, 0.05 μg/ml blasticidin S; APC883 + β-catenin shRNA, 1 μg/ml puromycin;. To screen for genes able to stimulate proliferation of APC883 cells because of rescue of mitotic defects and delayed doubling time, APC883 cells were screened after transfection by continuous culture for more than 1 month without drug selection, which allowed unconstrained proliferation. For prolonged culture, the rescued cells were maintained in medium that contained appropriate selection drugs as described above.

### Proteins

For bacterial expression, DNA fragments encoding APC proteins were inserted into pGEX-5X-1 (GE Healthcare Life Sciences, Pittsburgh, PA, USA). For *in vitro* transcription and translation reaction, AURKA cDNA was inserted into pcDNA3.1 (Invitrogen) and pT7-FLAG-1 expression vectors (Sigma-Aldrich) to generate HA-AURKA and FLAG-AURKA, respectively, and proteins were expressed with a TNT Quick Coupled Transcription/Translation System (Promega, Madison, WI, USA) according to the manufacturer’s protocol. Purified human AXIN1-MYC/DDK (TP308349) and TPX2-MYC/DDK (TP305821) proteins were obtained from OriGene Technologies (Rockville, MD, USA). GST-tagged AURORA A, Active (A28-18G) and Inactive (A28-14G) proteins were obtained from SignalChem (Richmond, Canada). Purified XMP215-His protein has been described previously (63).

### In-cell fluorescence colocalization assay, immunoprecipitation and in vitro pull-down assays and cell proliferation assays

For preliminary detection of protein-protein association, target molecules were tagged with GFP (EGFP or Emerald GFP) or RFP (TagRFP-T or mApple) and overexpressed in HEK293T cells, and their colocalization analyzed under fluorescence microscopy (in-cell colocalization assay). For *in vitro* pull-down assays, proteins fused to GST were synthesized in *E. coli* and isolated by absorption to Glutathione Sepharose (GE Health Care). GST and GST-fusion proteins (2 μg) immobilized to beads were incubated for 1 h at 4°C with *in vitro* transcription/translation products (HA-AURKA and FLAG-AURKA) or purified XMAP215-His protein in 500 μl of binding buffer (0.1% NP-40, 50 mM HEPES (pH 7.0), 150 mM NaCl, 50 mM NaF, 5 mM EDTA, 1 mM DTT, 1 μg/ml aprotinin, 1 μg/ml leupeptin, 50 μg/ml PMSF) and then washed extensively three times with binding buffer. Proteins adhering to the beads were resolved by sodium dodecyl sulfate polyacrylamide gel electrophoresis (SDS-PAGE), transferred to a polyvinylidene difluoride membrane filters (Millipore), and analyzed by immunoblotting. For the pull-down experiments, HEK293 cells were transfected with FLAG-tagged AXIN1-DIX(600-862) and extracted in lysis buffer (50 mM Tris HCl pH 7.5, 150 mM NaCl, 5 mM EDTA, 1% Triton X-100, 2 mM Na3VO4, 10 mM NaF, 1 mM DTT). Cell lysates (500 μl) were incubated with 2 μg GST or GST-APC-221-880 bound to Glutathione Sepharose beads for 1 h at 4°C. The beads were then washed three times with wash buffer (20 mM Tris HCl pH 8.0, 150 mM NaCl, 0.5% NP-40) and boiled in SDS sample loading buffer. Protein complexes were separated by SDS-PAGE and detected by immunoblotting.

### In vitro kinase assay

AURKA autophosphorylation was evaluated using the ADP-Glo Kinase Assay (Promega) according to the manufacturer’s instructions. The ADP-Glo Kinase Assay is a luminescent kinase assay that measures ADP formed via a kinase reaction. Briefly, inactive AURKA (0.3 pmol; SignalChem) was mixed with various concentrations (0.5-2 pmol) of purified GST, Axin1 (OriGene Technologies) or TPX2 (SignalChem) for 10 min at RT. The mixture was further incubated with 50 µM ATP in Reaction Buffer A (40 mM Tris, pH 7.5, 20 mM MgCl_2_, 0.1 mg/ml BSA) supplemented with 50 µM dithiothreitol (DTT) in a 25 µl volume for 30 min at RT. The reaction was terminated by adding 25 µl of ADP-Glo Reagent. After incubation for 40 min at RT, 50 µL of the Kinase Detection Reagent was added and incubated for 30 min at RT. The luminescence was measured using a luminometer (Berthold Technologies, Bad Wildbad, Germany). Data from three independent experiments were averaged after normalization against the control value (AURKA only) which was set to 1.0.

### Western blotting

Total cell lysates were prepared in Laemmli SDS-sample buffer supplemented with Protease Inhibitor Cocktail (Nacalai Tesque, Kyoto, Japan) and protein concentrations were quantified by the DC protein assay (Bio-Rad, CA, USA). Protein concentrations were normalized if necessary and boiled for 5 min with 2-mercaptoethanol, loaded the same amount of protein in each lane, separated by SDS-PAGE, and transferred to 0.45 μm Immobilon-P polyvinylidene fluoride membranes (Millipore). Proteins were visualized using Amersham ECL Select Western Blotting Detection Reagents (GE Healthcare) or Chemi-Lumi One Super reagent (Nacalai Tesque) and detected with a chemiluminescence imaging system Davinch-Chemi (CoreBio, Taoyuan, Taiwan) or Amersham Imager 680 (GE Healthcare). For detection of phosphorylated proteins, cell lysates were immediately boiled for 5 min with 2-mercaptoethanol without concentration adjustment.

### Fluorescence microscopy of immunostained specimens and data analysis

The immunostained specimens were observed using an LSM780 confocal microscope with a Plan-APOCHROMAT 63×/1.4 NA oil immersion objective, a Plan-APOCHROMAT 40×/1.3NA oil immersion objective, a Plan-APOCHROMAT 10×/0.45 NA objective, four laser lines (405 nm; Multi-Ar, 458, 488, 514 nm; DPSS, 561 nm; He-Ne, 633 nm), and a highly-sensitive gallium arsenide phosphide (GaAsP) detector (Carl Zeiss) and an LSM880 confocal microscope with a Plan-APOCHROMAT 63×/1.4 NA oil immersion for SR objective, a Plan-APOCHROMAT 40×/1.3 NA oil immersion for SR objective, a Plan-APOCHROMAT 10×/0.45 NA objective, four laser lines (405 nm; Multi-Ar, 458, 488, 514 nm; DPSS, 561 nm; He-Ne, 633 nm), and a GaAsP detector (Carl Zeiss). Detection and quantitative analysis of phosphorylated proteins were performed using cultured cells and paraffin-embedded tissues fixed in 4% paraformaldehyde or formalin, respectively. To compare immunostaining intensities in each cell type cultured on different cover slips, the cover slips were processed under the same conditions and intensity values were obtained as relative values against that of control cells, which was set to 1. To compare immunostaining intensities in different tissue sections, the sections were bound to a single slide glass to be processed under the same staining conditions. For tumor cells analysis in the intestinal tissue specimens, tumor cells in each polyp were specified by their epithelial features shown with E-cadherin or β-catenin immunostaining.

To measure fluorescence on centrosomes, maximum intensity projections of z-stack images were produced, and centrosomal regions were manually selected to obtain total fluorescence values in the region of interest using the ImageJ software. The average fluorescence intensities and phospho-to-total protein ratios were normalized to the control values, which were set to 1.0. In the analysis of different MCF10A derivatives, data from different experiments were standardized and normalized using parental MCF10A cells as a control to compare AURKA total intensity, phospho intensity, and the phospho-to-total ratio. Finally, the values for MCF10A(20di) cells were normalized to 1.0. For mouse intestines, average values from crypt cells were used as standards. For β-catenin-labeling in mouse intestine, values from crypt cells in the neighbourhood of the polyps along the intestinal tract were used as standards. For human tumors, the average values in polyp#2 were normalized to those of polyp#1 without *APC* mutation. Sample sizes indicate the numbers of cells (β-catenin)/centrosomes (AURKA) analysed. To define the percentage of Ki67-positive cells in AURKA inhibitor–treated mouse tumors, paraffin-embedded tissues fixed in formalin were stained for β-catenin, Ki67, and nuclear DNA (with Hoechst 33342), imaged on a confocal microscope, and analysed using the MetaMorph software. The Ki67- and Hoechst 33342–positive areas were determined by intensity thresholding in the crypts and tumor tissues, and the percentage of Ki67-positive areas were calculated in each image obtained from multiple crypt/tumor areas. Sample sizes indicate the numbers of images analysed. To count mitotic cells in mouse intestines, the tissue sections were stained with Hoechst33342 and imaged using an Imager Z1 fluorescence microscope (Carl Zeiss), and total and mitotic cells were manually counted based on the shapes of chromosomes.

### Live imaging and data analysis

Time-lapse phase-contrast live imaging of cells was performed using an IX81-ZDC inverted microscope with a 20× objective lens LUCPLFN 20×PH (Olympus) controlled by MetaMorph software (Molecular Devices, Sunnyvale, CA, USA). The microscope was installed in a temperature-controlled dark box and was equipped with a motorized XYZ stage, a ZDC (Z drift compensator) autofocus system, a cooled CCD camera ORCA-AG (Hamamatsu Photonics, Shizuoka, Japan), and a heat controller (Tokai Hit, Shizuoka, Japan) and gas mixer (Olympus/Tokken, Chiba, Japan) for cell culture. During imaging cells were maintained at 37°C in a 5% CO_2_ atmosphere. Duration of mitosis and doubling time were manually measured. Time-lapse images were edited using ImageJ and QuickTime Pro software (Apple).

### Animals and human colorectal adenoma

Animals were used in accordance with governmental and institutional regulations and protocols were approved by the Animal Care and Use Committee of the RIKEN Kobe Institute, Kanazawa University and the University of Tokyo. All animals were cared for in accordance with the ethical standards of the institutions. *Apc^Δ716^* mice and paraffin-embedded tissue preparation were described previously (64). *Apc^1638T^* mice (backcrossed to C57BL/6) were described previously (34,65). Wild-type C57BL/6J mice were purchased from Charles River Laboratories Japan Inc. (Kanagawa, Japan). For AURKA inhibitor treatment, wild-type and *Apc^Δ716^*mice were dosed orally twice daily for 14 consecutive days for Ki67 staining (two mice in each group) and 28 consecutive days for polyp number counting (ten mice in each group), with vehicle, 20 mg/kg alisertib (CS-0106, Chemscene LLC, Monmouth Junction, NJ, USA) as a solution in 10% 2-HP-β-CD (66). For polyp counts, incised intestinal tracts were briefly stained in 0.1% methylene blue, and counted under a dissecting microscope and were categorized according to size (<1, 1–2 and >2 mm).

Human colorectal polyps were excised by polypectomy at the Nico-tama Coloproctology Clinic with informed consent and the protocol was approved by the Ethical Committee of the Riken Kobe Institute. Samples were de-identified and analyzed anonymously. A small portion of the polyp was used for genomic DNA isolation using a DNeasy Blood & Tissue Kit (Qiagen, Hilden, Germany), and the remaining portion was immediately fixed in formalin and embedded in paraffin. For genomic sequencing, the last exon of *APC*, encoding from the armadillo repeat region to the stop codon, was cloned by PCR as a fragment of 1000∼1700 bp using specific primers, inserted into pCR-Blunt II-TOPO (Thermo Fisher Scientific, Waltham, MA, USA) and analyzed by Sanger sequencing. In this study, a hyperplasia polyp without APC mutation (polyp#1, Asian female, early 60s) and a low-grade tubular adenoma with two truncating mutations (c.2698G>T and c4558G del) (polyp#2, Asian female, late 50s) were used. Images of hematoxylin-eosin (HE) stained sections of a formalin-fixed, paraffin-embedded Swiss roll of the intestine of an *Apc^Δ716^* mouse and of human polyps were collected using a SZX12 stereomicroscope with a DF PLAPO 1.2× PF2 objective (Olympus, Tokyo, Japan) or a Primo Vert Monitor with a Plan-ACHROMAT 4×/0.10 NA objective (Carl Zeiss, Jena, Germany), respectively.

### Mutation and copy number analyses using TCGA data

Exome and SNP array data from The Cancer Genome Atlas (TCGA)-colon adenocarcinomas (COAD) and rectum adenocarcinomas (READ) paired tumor/normal samples were prepared as described previously (67). Briefly, mutation calling from the exome data was performed using the *EBCall* algorithm (68) and absolute copy number profiles were obtained from the SNP array data using the ASCAT algorithm (69). CNA scores (focal CNA score, FCS; broad CNA scores, BCS; global CNA scores, GCS) and arm-level copy number profiles were obtained from the SNP array data using the default settings (including re-segmentation) of CNApp tools (38). In total, 319 COAD and 127 READ TCGA patient samples were analysed. The 446 patient samples were divided into four groups as follows. Class 1: Lesions without a truncating mutation and LOH in the *APC* locus; Class 2: Lesions with one truncating mutation or one LOH in the *APC* locus, which are expected to have a single *APC* mutation and retain a single wildtype copy of *APC*; Class 3: Lesions with two or more (two to three as a result) truncating mutations in the *APC* locus, or one or more (one to three as a result) truncating mutations, and LOH in the *APC* locus, which are expected to have no wildtype *APC*. *CTNNB1*: patients with mutations in β-catenin gene (*CTNNB1*). The microsatellite instable status and clinicopathological parameters of all COAD and READ patient samples were also obtained from TCGA database. The samples were divided into two subgroups, microsatellite stable (MSS), which included microsatellite instable (MSI) low and MSI high in accordance with the MSI status, using TCGAbiolinks (70) and then the presence or absence of an *APC* mutation was compared between each subgroup. Survival curves and CNA frequencies were analysed as described in the available code. These data were analysed by Fisher’s exact test. More details are in the available code.

### Comparative genomic hybridization (CGH) microarray analysis

CGH microarray analysis was performed by Takara Bio using a SurePrint G3 Mouse CGH Microarray Kit, 1×1M (Agilent Technologies, CA, USA). Genomic DNA from *APC^Δ716^* mouse normal intestines (control) and polyps were fragmented, amplified, and then labelled with Cy3 or Cy5 dyes. Dye-swap replication experiments were performed for each polyp sample vs. control DNA. The hybridised DNAs were applied to the microarray and scanned on an Agilent SureScan Microarray Scanner. The results were analysed using Agilent Genomic Workbench 7.0 software (Agilent Technologies. Experimental parameters were set by considering the mosaicism of mutations in tumor cells and the presence of a considerable amount of normal cells in each sample. For preprocessing: Feature Filter, ON; Design Filter, v2; GC correction, ON; Centralization, ON; Intra array, ON. For Aberration Filter: Log_2_Ratio≥0.01; minimum number of probes in region, 5. For analysis, the AMD-2 algorithm with threshold 6 was used. Array CGH profiles derived from two dye-swap experiments were plotted together on the same plot. Synteny between human and mouse chromosomes were analysed on the Ensembl Comparative Genomics/Synteny website. Ideograms of human chromosomes 13 and 20, and mouse chromosomes 2 and 14 shown in Fig. 4B were generated by tracing illustrations in Ideogram View on the NCBI Genome Data Viewer website. Syntenic comparison of human and mouse chromosomes was performed using the Comparative Genomics/Synteny tool on the Ensembl website.

### Genomic DNA isolation and quantitative PCR (qPCR)

Genomic DNA was extracted from mouse tissues using ISOGENOME (Nippon Gene, Tokyo, Japan) in accordance with the manufacturer’s instructions. Five nanograms of genomic DNA was applied to qPCR using PowerUp SYBR Green Master Mix (Applied Biosystems, Foster City, CA, USA) and the QuantStudio 5 Real-time PCR System (Applied Biosystems). The reaction was first incubated at 50°C for 2 min and then at 95°C for 2 min, followed by 40 cycles at 95°C for 15 sec and 60°C for 1 min. *Zfp438* located on mouse chromosome 18A1 was used as the internal reference gene for qPCR because our CGH microarray analysis did not show the presence of genetic alterations in the A1–A2 region of chromosome 18 in intestinal polyps of *Apc*^Δ*716*^ mice. The results were normalized to the values obtained for *Zfp438* and analysed by the 2^−ΔΔCT^ method. The primer sequences were as follows: m*Aurka* 5ʹ-TGGCTCAAACGGAGAGTTGA-3ʹ and 5ʹ- CCTGGAATCCCATACCTGGA-3ʹ; m*Bora*, 5ʹ-TAGGTTGGACGGTCCCTCAC-3ʹ and 5ʹ-GCAGCACAGACATGGAGAGC-3ʹ; m*Tpx2*, 5ʹ-GCATGCGTTTTGTTCCTCTG-3ʹ and 5ʹ-GCTGCATGATCCCAATGCTA-3ʹ; and m*Zfp438*, 5ʹ- GGCTGTTGCTGTGCTGTACC-3ʹ and 5ʹ-CTCCCTGGGCCATCAGTAAG-3ʹ.

### Karyotyping of chromosomes in MCF10A and APC883 cells

The APC833 cell line was generated from the MCF10A cell line, a genetically stable human mammary gland epithelial cell line (71), by introducing a truncation mutation into the APC locus by genome editing. The MCF10A cell line was established in 1990 from human fibrocystic mammary tissue (39). Karyotyping prior to immortalization showed normal diploidy, but later passages showed near-diploidy with some rearrangements interpreted as 48,XX,3p-,6p+,8+,9p+,16+, including a balanced reciprocal translocation t(3;9)(p13;p22) and an unbalanced translocation t(5q;der[9]) (39) In later studies, gain at chromosome 1q was reported (45,72), as observed in this study. Trisomy 20 is recurrent in subsequent studies (40,45,72,73),

In this study, Q-banding (20 cells for each cell type) were performed by Chromosome Science Labo (Hokkaido, Japan). Complete description of karyotyping results is shown in Table S3. In our early analysis, detailed rearrangement of one chromosome 8 was not determined and was described as “dup8(p?)” In later analysis, the structure of the rearranged chromosome 8 was analyzed by FISH using specific probes (8p11, 8p23, 8q13, 8q21, 8q24) and the description was updated to “der(8)dup(8)(q22q24.3)ins(8)(p24.3q22q24.3)”. For the transformed, near-tetraploid APC883 cells, common number imbalances are shown by letter color for karyotype description: cyan, three chromosome 6s; brown, five chromosome 8s; red, six chromosome 20s (Table S3), and by box color for chromosome images: cyan, three chromosome 6s; brown, five chromosome 8s; red, six chromosome 20s. To analyze newly generated chromosome aberrations, types of abnormalities were classified as tetraploidy, gain, loss, deletion and fusion. The fusion includes all types of chromosomes generated by fusions such as translocation, robertsonian translocation and telomere association. To separate subclones MCF10A cells carrying chromosome 20 disomy [MCF10A(20di)] and trisomy [MCF10A(20tri)], clones were isolated by colony picking followed by Q-banding to determine karyotype.

### Statistics and reproducibility

Statistical evaluation was carried out by the Student’s t-test (two sided), Tukey–Kramer method (two sided), Fisher’s exact test, Wilcoxon rank-sum test, or Mann–Whitney U-test (two sided) as indicated. P-values and sample size are shown in each figure. Biochemical experiments were repeated with at least three independent preparations with similar results. For quantification of image data to ensure adequate power to detect significance, sufficient numbers of regions/cells were analysed as indicated in each figure.

## Data and materials availability

Data are available on request from the authors. Materials used in this study are available for distribution after a materials transfer agreements (MTA), except for materials from human research participants. The aCGH data have been deposited in the NCBI Gene Expression Omnibus (GEO; http://www.ncbi.nlm.nih.gov/geo/) and are accessible through GEO series accession number GSE3642. accession GSE63306 (https://www.ncbi.nlm.nih.gov/geo/query/acc.cgi?acc= GSE63306). The “Fig1c.zip” file containes multiple csv files used to generate the plots in Fig. 1C.

## Code availability

Computer codes are available on request from the authors. The code to create the 2D rotational projection of EB1 comet data has been deposited in Github (https://github.com/wxchew/cometdensity/blob/master/2DProjectedCometDensity.ipynb). Codes for TCGA data analysis are available in Code S1. This folder also contains the TCGA data used for the analysis with the codes attached.

## Acknowledgements

We are grateful to Dr. A. Ochiai (NCC-EPOC) for his guidance on tumor analysis. Drs. A. Nagafuchi (Nara Medical Univ), T. Toda (Hiroshima Univ), M. Toya (Waseda Univ), M. Takeichi (Riken BDR), C. Furusawa (Univ. of Tokyo), and Y. Takai (Kobe Univ) for critical reading of the manuscript and valuable suggestions. Our special thanks are also due to Ms. M. Yamaguchi and Mr. Y. Sato (Carl Zeiss Japan), Mr. M. Yamaguchi (Molecular Devices Japan), and Mrs. Y. Imai and M. Emi (Olympus) for excellent technical assistance in operating the microscope systems and image analysis software.

## Funding

Y.M.-K. was supported by the Japan Society for the Promotion of Science– NEXT program (LS128), the Takeda Science Foundation, the Uehara Memorial Foundation, a Grant-in-Aid for Challenging Research (Pioneering) (JSPS KAKENHI Grant Number 20K20379), and Japan Science and Technology Agency Core Research for Evolutional Science and Technology (no. JPMJCR1863), Extramural Collaborative Research Grant of Cancer Research Institute, Kanazawa University, and an intramural grant from RIKEN. The results published here are in part based upon data generated by the TCGA Research Network: http://cancergenome.nih.gov/. Imaging experiments were in part supported by the RIKEN Kobe Light Microscopy Facility. Animal experiments were in part supported by the Laboratory for Animal Resources and Genetic Engineering at the RIKEN BDR.

## Author contributions

Y.K., T.H., K.O., A.H. and T.N. performed experiments and edited the manuscript. Y.W., M.O., S.S., and N.T. performed experiments. Y.K., A.N., and K.M. analyzed TCGA data using informatics techniques. T.K., T.S., M.O., K.F., A.K., P.O W. and H.O. prepared biological samples. K.K. performed image data analysis and statistical data analysis. N.Y., M.M., H.Y., C.W.X., S.A., K.T., W.R.L., B-C.C. and E.B. performed imaging experiments, image processing and data analysis. R.S. and R.F. prepared biological samples and edited the manuscript. M.O., M.M.T and T.A gave conceptual and technical advice. Y.M.-K. conceived and managed the project, performed experiments, and wrote the manuscript.

## Competing interests

Portions of the technology described herein are covered by U.S. Patent 7,894,136 issued to Eric Betzig (EB) and assigned to Lattice Light, LLC of Ashburn, VA, U.S. Patents 8,711,211 and 9,477,074 issued to EB and assigned to HHMI, U.S. Patent application 13/844,405 filed by EB and Kai Wang (KW) and assigned to HHMI, and U.S. Patent 9,500,846 issued to EB and KW and assigned to HHMI.

## Supplementary Materials

**Fig. S1.**
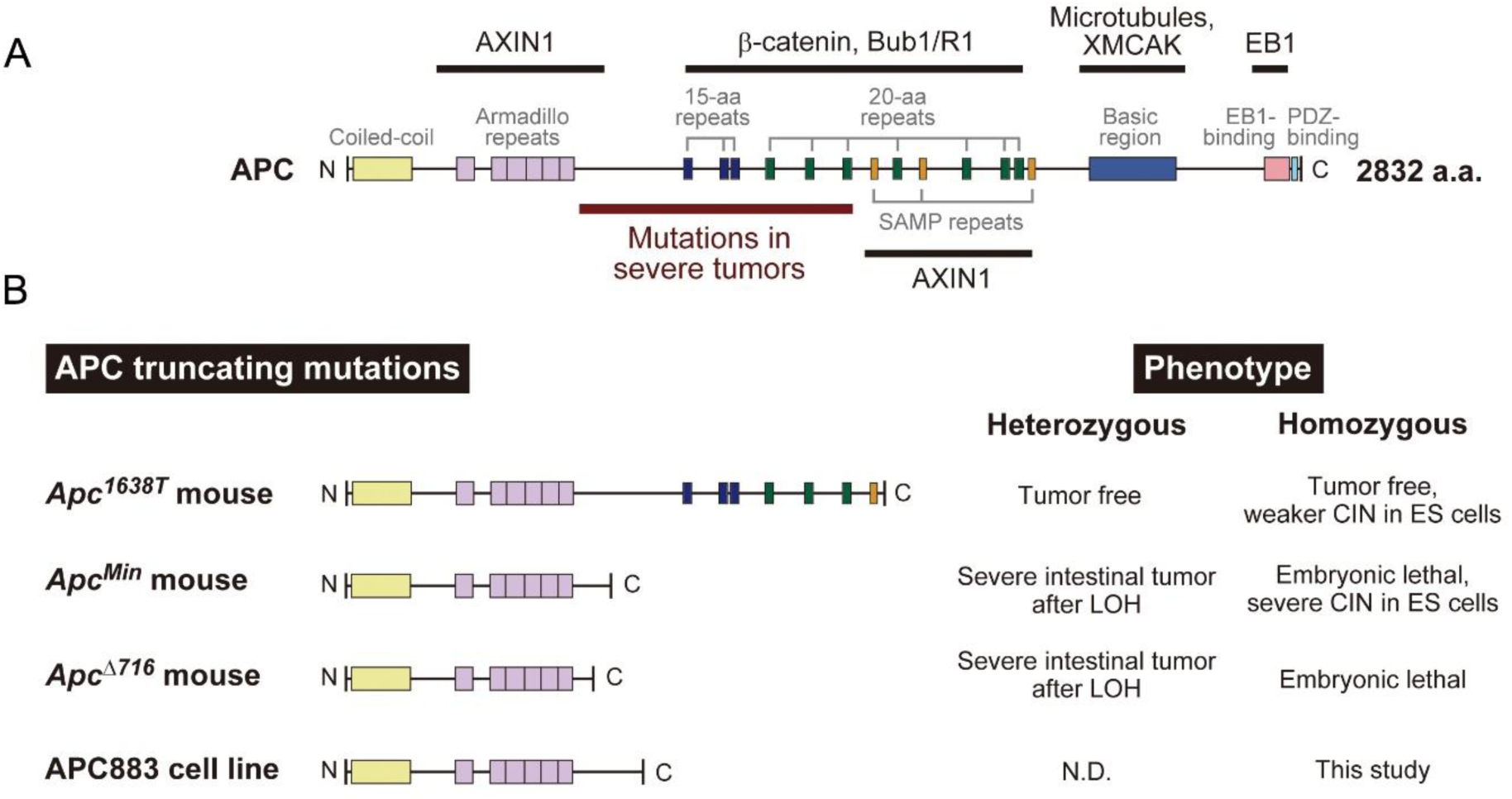
Structure and function of APC. (A) Domain structures of full length APC and major binding partners (1–7). The *APC* gene is responsible for many sporadic cases of gastrointestinal cancers and familial adenomatous polyposis, a dominantly inherited colorectal tumor predisposition that results from germline mutations in *APC* (8–14). Tumors start developing upon mutation of both alleles. In general, *APC* mutations that result in stable proteins truncated in the region between the armadillo repeat and the first SAMP motif are associated with severe polyposis phenotypes and, in the human colon, are associated with a very malignant colorectal cancer type (reviewed in (4,15)). (B) Truncated *APC* mutants. Phenotypes of cells and mice are shown on the right. Heterozygous *Apc* mutations, such as *Apc^Min^* and *Apc^Δ716^*, lead to severe polyposis after loss of the wildtype allele (>∼30 polyps), whereas their homozygosity results in embryonic lethality (16–18). Homozygous *Apc^1638T^* mice with a truncating mutation just after the first SAMP motif, which retains the β-catenin destruction ability, are viable and do not develop any tumors, thereby demonstrating that this N-terminal fragment that retains the first SAMP motif is sufficient for the tumor-suppressing function (19). Nevertheless, *Apc^1638T^* homozygous ES cells have a CIN phenotype, although it is weaker than that in *Apc^Min^* homozygous ES cells (20). MCF10A-derived cell line APC883 with a homozygous *APC* truncation was generated by genome editing.

**Fig. S2.**
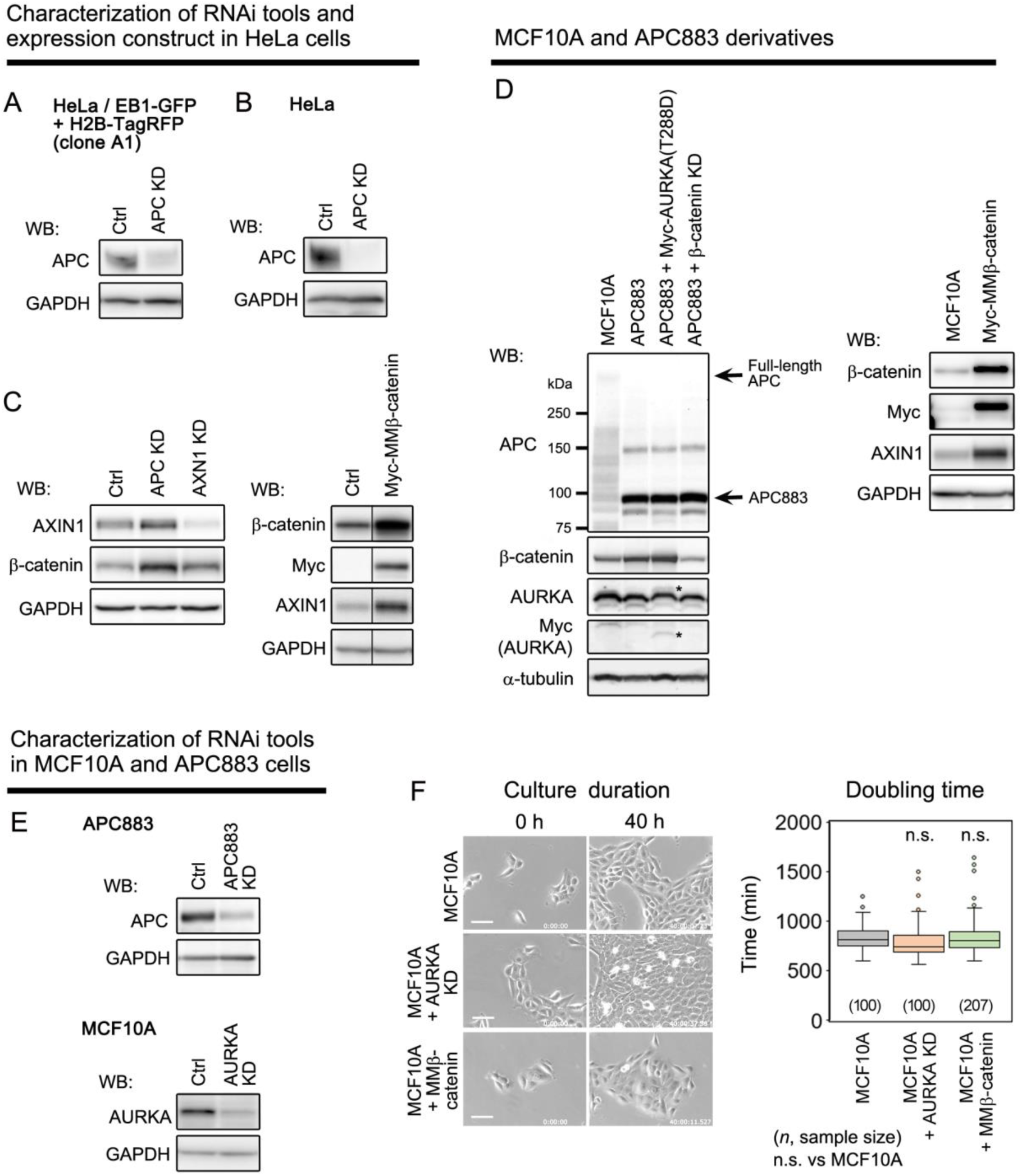
Characterization of RNAi tools, expression constructs, and cell lines used in this study. (**A, B**) Western blot (WB) analysis of HeLa cells and HeLa cells that expressed EB1-GFP and H2B-TagRFP (clone A1) (21) before and after constitutive transfection with shRNA against APC (APC KD). APC knockdown efficiency was ∼70%. (**C**) Characterization of RNAi tools and expression construct in HeLa cells as indicated using the indicated antibodies. (**D**) Characterization of MCF10A cells, APC883 cells, and APC883 cell-derived transfectants as indicated using the indicated antibodies. Full-length APC and truncated APC883 protein are indicated. For Myc-AURKA (T288D), endogenous and exogenous Myc-tagged proteins are shown. (**E**) Characterization of RNAi tools in MCF10A and APC883 cells as indicated. (**F**) Doubling time of parental MCF10A cells, AURKA knockdown MCF10A cells and MMβ-catenin expressing MCF10A cells. n.s. (Student’s t-test showed not significant, *P*>0.05).

**Fig. S3.**
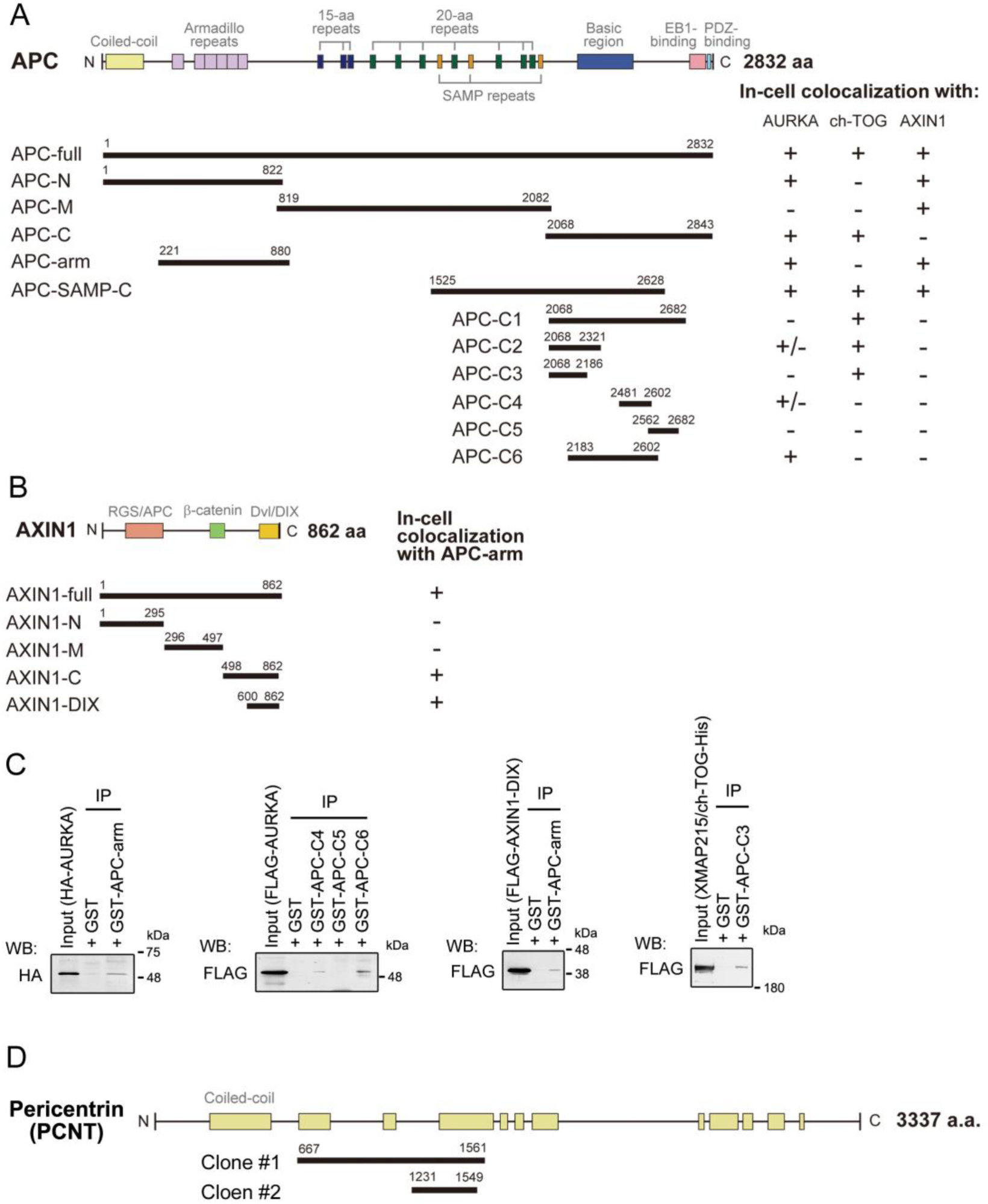
(**A, B**) Domain structures of APC and AXIN1, and the analyzed fragments. Amino acid numbering is based on human APC transcript variant 3 (NM_000038) and human AXIN1 transcript variant 1 (NM_003502). Associations of APC with AURKA and AXIN1 were assessed by in-cell colocalisation analysis using a series of deletion mutants. The results are shown on the right. (**C**) Direct binding between APC fragments and AURKA/AXIN1/ch-TOG predicted in (A) was confirmed by in vitro pull-down assays using purified proteins. The APC arm and APC-C6 bound to AURKA. The APC arm also bound to the C-terminal DIX domain of AXIN1. XMAP215 (22), the *Xenopus* homologue of ch-TOG, bound to APC-C3. We used Xenopus XMAP215 because we could not purify full length human ch-TOG from E. coli. The C-terminal one-third bound to the APC arm. See also Table S2. (**D**) Yeast two-hybrid screening using the APC arm region as bait identified pericentrin. The two identified pericentrin clones are shown.

**Fig. S4.**
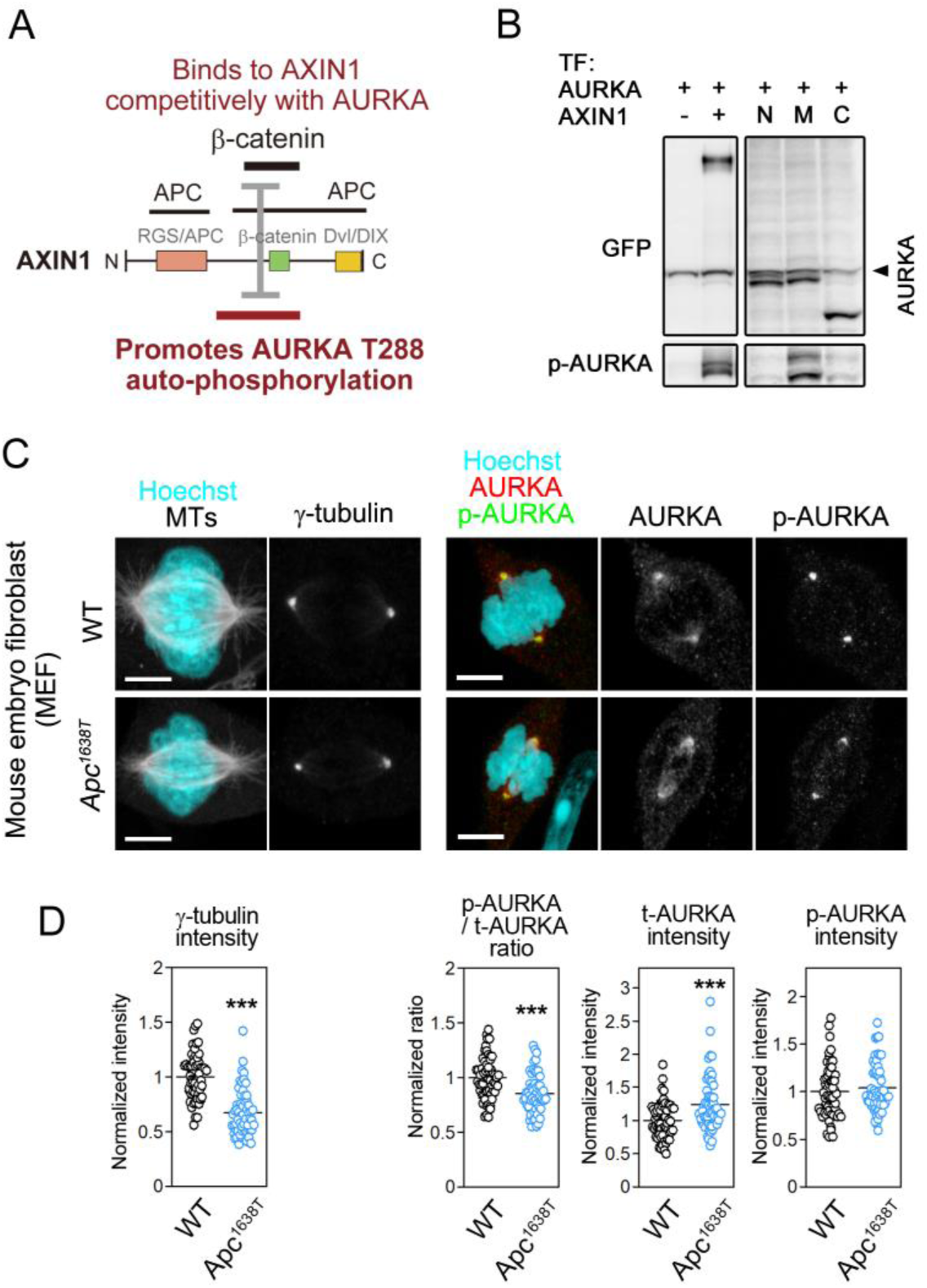
Functions of AXIN1 and *Apc^1637T^* gene expression products. (**A**) AXIN1 structure and AURKA-activating region that also binds to β-catenin (23). (**B**) Effects of AXIN1 on AURKA phosphorylation were analyzed using the HEK293T cell overexpression assay system. HEK293T cells were transfected with GFP-fused AURKA and full length AXIN1 or fragments shown in Fig. S3B, lysed, and then subjected to western blot analysis using anti-GFP and anti-p-T288 AURKA antibodies. The left panel is the same as Fig. 2E. See also Table S2. (**C, D**) Analysis of primary MEFs from wildtype (WT) and *Apc^1638T^* mice. Immunostaining of microtubules (MTs), γ-tubulin, AURKA, and p-T288 AURKA (C). Immunostaining intensity of γ-tubulin, the autophosphorylation ratio (p-AURKA/t-AURKA ratio), total AURKA protein levels (t-AURKA), and autophosphorylated AURKA levels (p-AURKA) were normalized to those of the wildtype, which were set to 1.0 (**D**). Scale bars: 5 μm. \*\*\**P* < 0.001 (Student’s t-test).

**Fig. S5.**
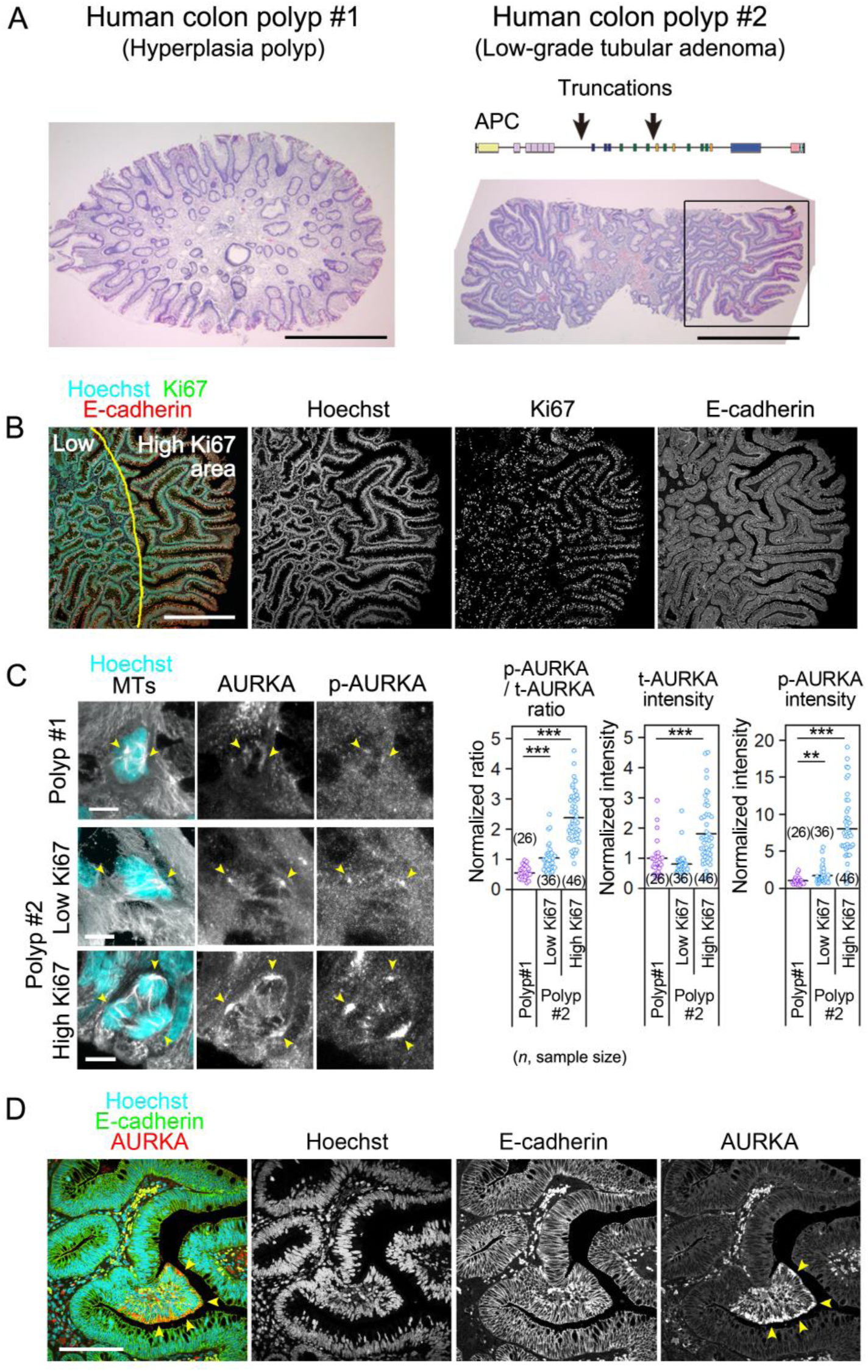
Immunofluorescence analysis of paraffin-embedded sections of *APC*-mutant human tumor tissues. (**A**) HE-stained sections of a human hyperplasia polyp without an APC mutation (polyp #1) (left) and a low grade tubular adenoma with two APC truncation mutations (polyp #2) (right). Two mutant APC sequences detected in polyp #2 are shown (arrows). For details, see Materials and Methods. (B) Ki67 proliferation index in a low grade tubular adenoma with two APC truncation mutations (polyp #2). On the basis of the Ki67 intensity, the area was roughly divided into a highly proliferative region (outer, high Ki67 area) and moderately proliferative region (inner, low Ki67 area) as indicated by the yellow line. (C) Representative images of mitotic cells in polyp #1 and low and high Ki67 areas of polyp #2 (left). The centrosomes are indicated by yellow arrowheads. Quantitative analysis of immunofluorescence signals of AURKA in polyp #1 and polyp #2 (right) at the centrosomes. (D) Immunostaining of polyp #2 for E-cadherin and AURKA with nuclear DNA stained by Hoechst 33342, showing appearance of a patchy region with extremely high AURKA expression levels (yellow arow heads). Scale bars: 200 μm (A), 100 μm (C, E), 5 μm (B). \*\**P* < 0.01; \*\*\**P* < 0.001 (Tukey–Kramer test).

**Fig. S6.**
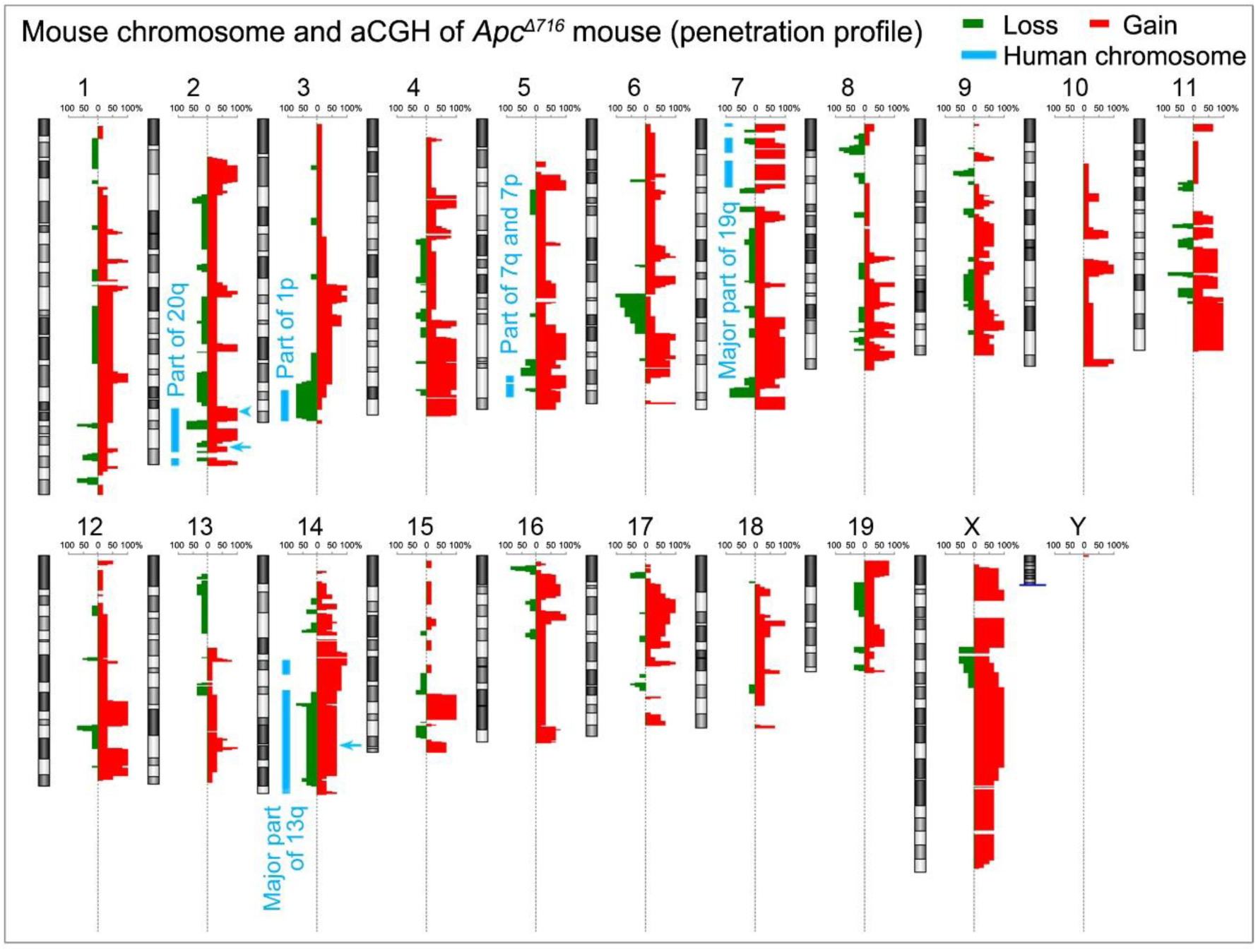
aCGH analysis of mouse tumors and syntenic comparison with human TCGA analysis. aCGH analysis of chromosomes in three independent *Apc^Δ716^* mouse polyps. A penetration profile of copy number changes from duplicate analyzes is shown. Chromosomal regions showing cross-species overlaps showing similar gain/loss pattern in two or more samples are indicated by cyan lines. Arrow and arrowhead on chromosome 2 indicate the loci of *Aurka* and *Tpx2*, respectively. Arrow on chromosome 14 indicates the locus of Bora.

**Fig. S7.**
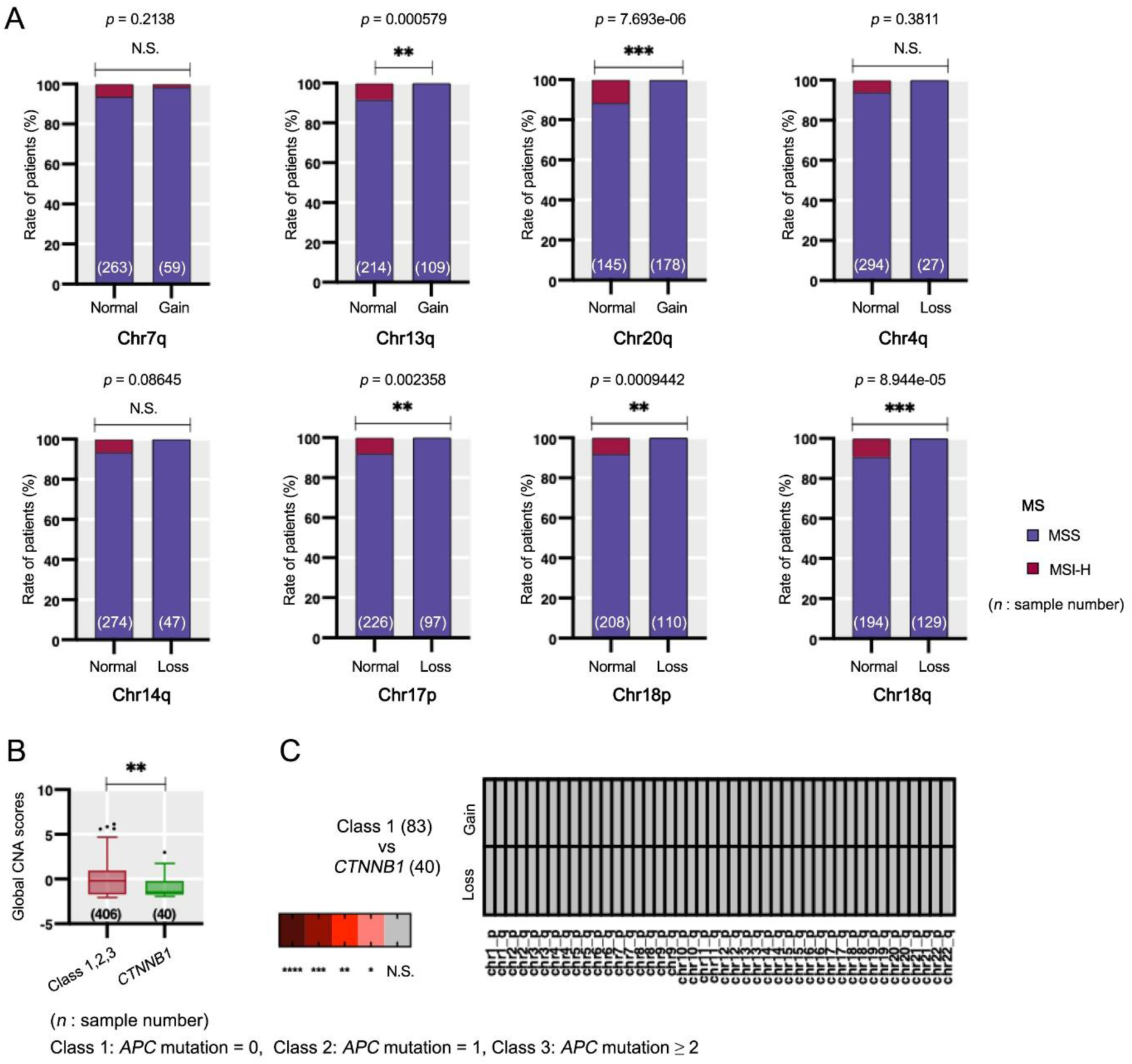
TCGA data analysis for MSI status. (**A**) Frequency of microsatellite stable (MSS) and microsatellite instable (MSI) in copy number changes in chromosomes with the most frequent alterations. Fisher’s extract test, \*\**P* < 0.01; \*\*\**P* < 0.001. (**B**) Global CNA score (GCS) distribution in *CTNNB1* (*β-catenin*) wildtype group (Classes 1–3) and *CTNNB1* mutant group. Class 1: *APC* mutation = 0; Class 2: *APC* mutation = 1; Class 3: *APC* mutation ≥2. Mann-Whitney test significance is shown as \*\**P* < 0.01. (**C**) Class1 and CTNNB1 mutant were without significant difference. \**P* < 0.05; \*\**P* < 0.01; \*\*\**P* < 0.001 \*\*\*\**P* < 0.0001 (Fisher’s extract test).

**Fig. S8.**
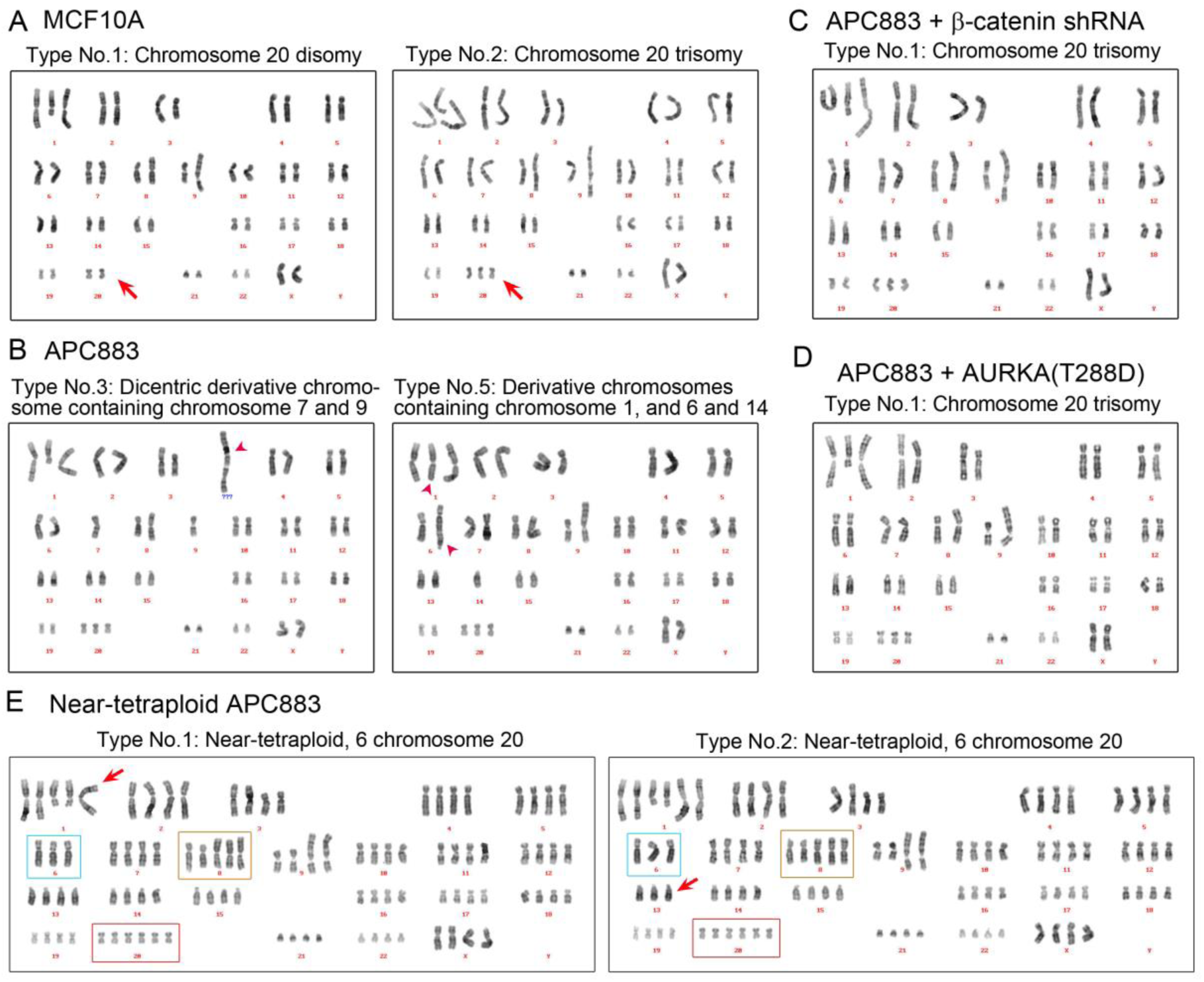
Karyotypes of MCF10A and APC883 cells. Representative karyotype images of each cell line. (20 cells were analyzed in each cell line). For complete karyotype descriptions of all analyzed cells, see Table S3. The karyotypes were determined by Q-banding with Hoechst/quinacrine staining. Note that commonly observed abnormalities in chromosome 1 (deletion and partial duplication), chromosome 3 (deletion and translocation), chromosome 8 (duplication), and chromosome 9 (translocation) have been observed previously (24,25). (**A–D**) Karyotype of parental MCF10A cells, APC883 cells, and APC883 derivatives. Major karyotypes in each cell type and representative examples with chromosome aberrations are shown. Chromosome 20 is indicated by red arrows. The parental MCF10A cell line has two major populations with chromosome 20 disomy or chromosome 20 trisomy (types 1 and 2, respectively). All APC883 cells and APC883-derive cells observed have chromosome 20 trisomy, the same as MCF10A type No. 2. Newly derived abnormal chromosomes are indicated by red arrowheads. (**E**) Representative karyotypes of transformed APC883 cells appeared after >5 months of passaging are shown. In this transformed subpopulation, all analyzed cells had a near tetraploid karyotype with some numerical imbalances. Common numerical imbalances are shown by the letter color for the karyotype description: cyan, three chromosome 6s; brown, five chromosome 8s; red, six chromosome 20s. Examples of randomly occurring imbalances are indicated by arrows.

### Legend for Tables S1 to S3, Movies S1 to S5, and Code S1 (separate files)

**Table S1.**
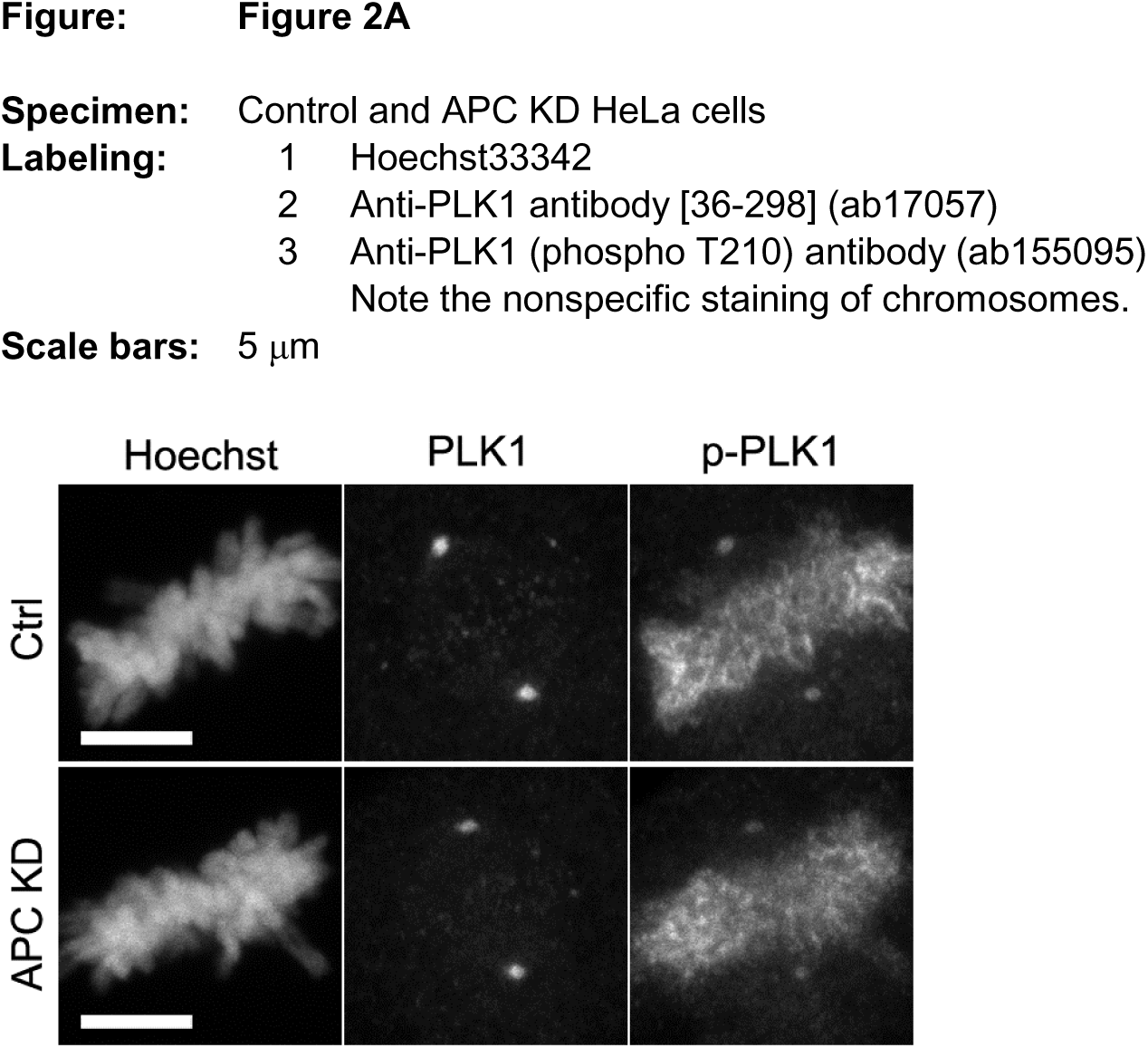
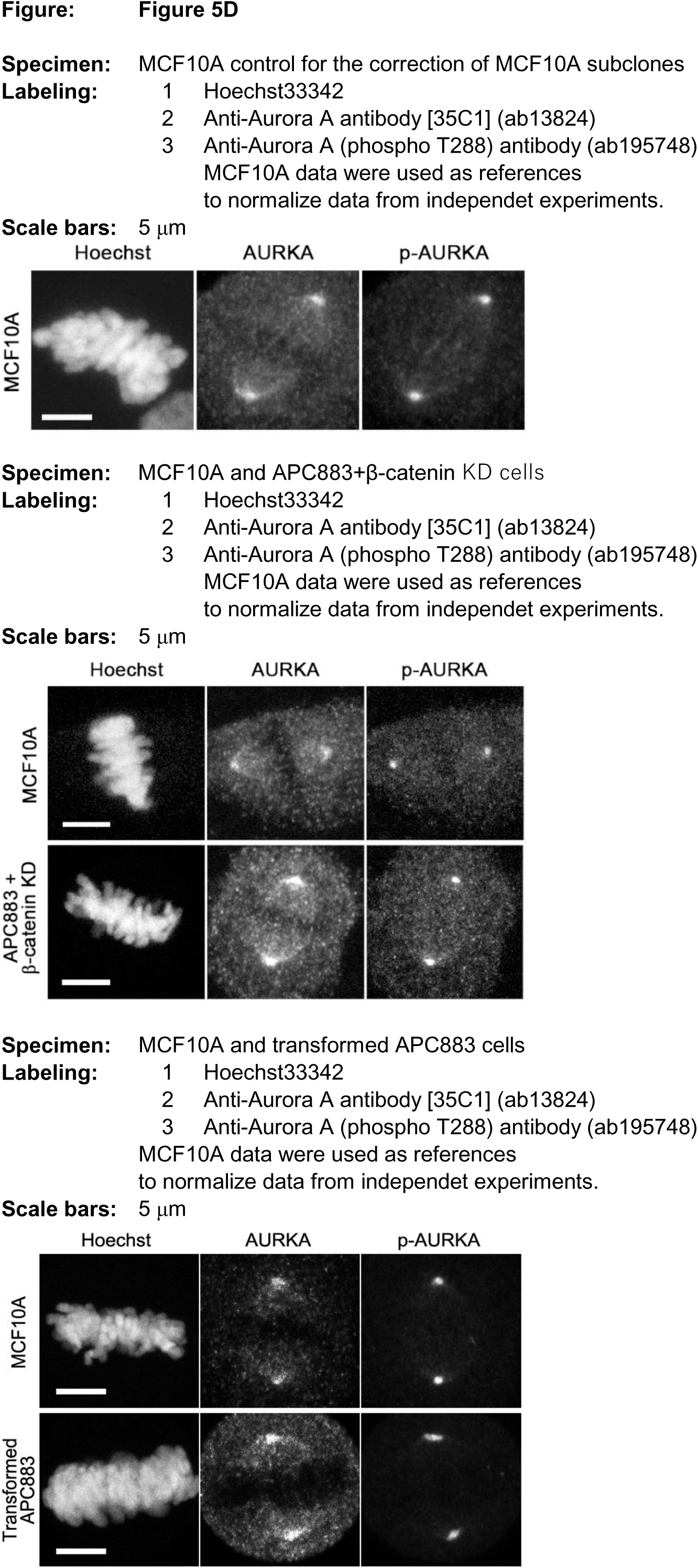
Representative images used in the analysis, which could not be presented in the main body.

**Table S2.**
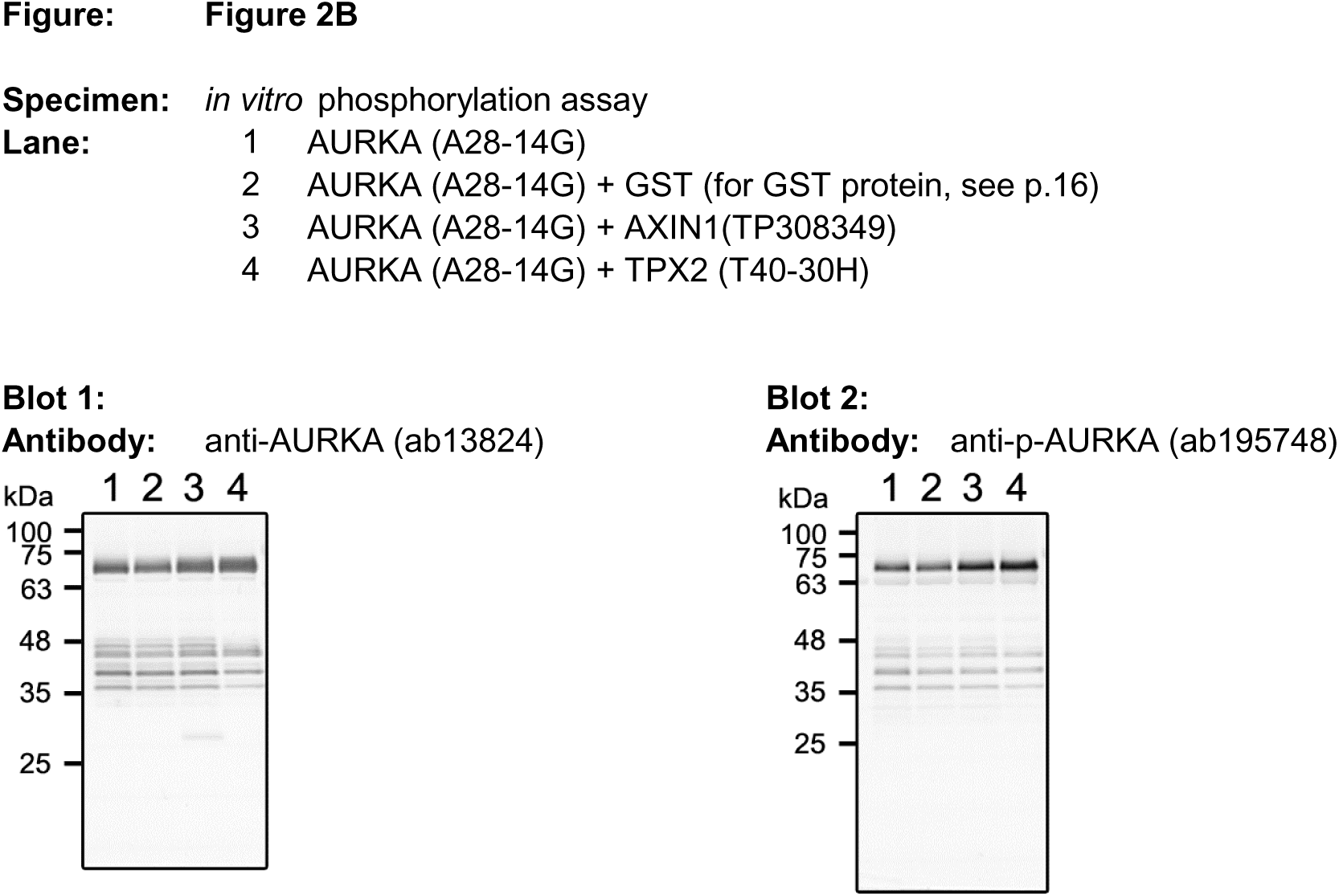
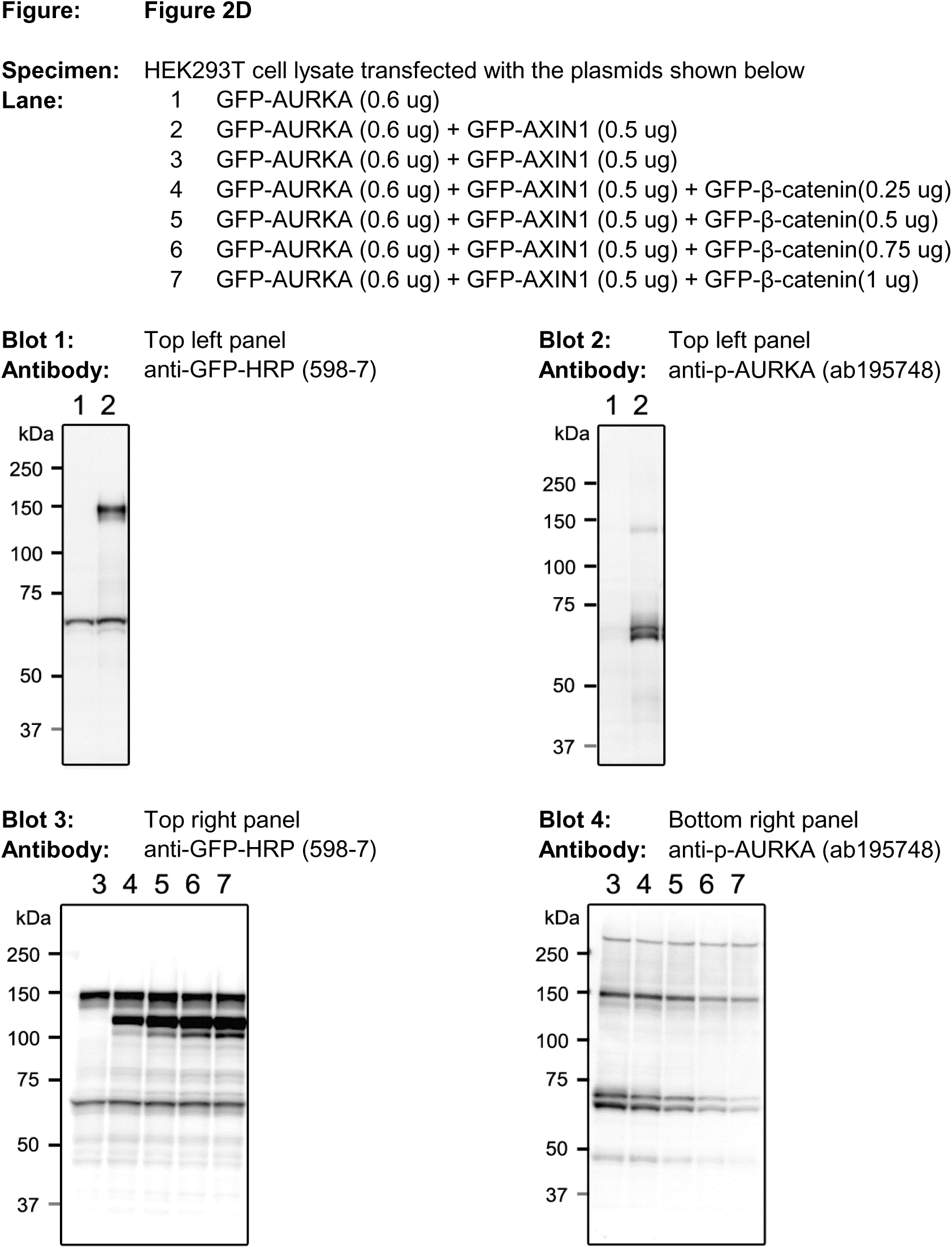
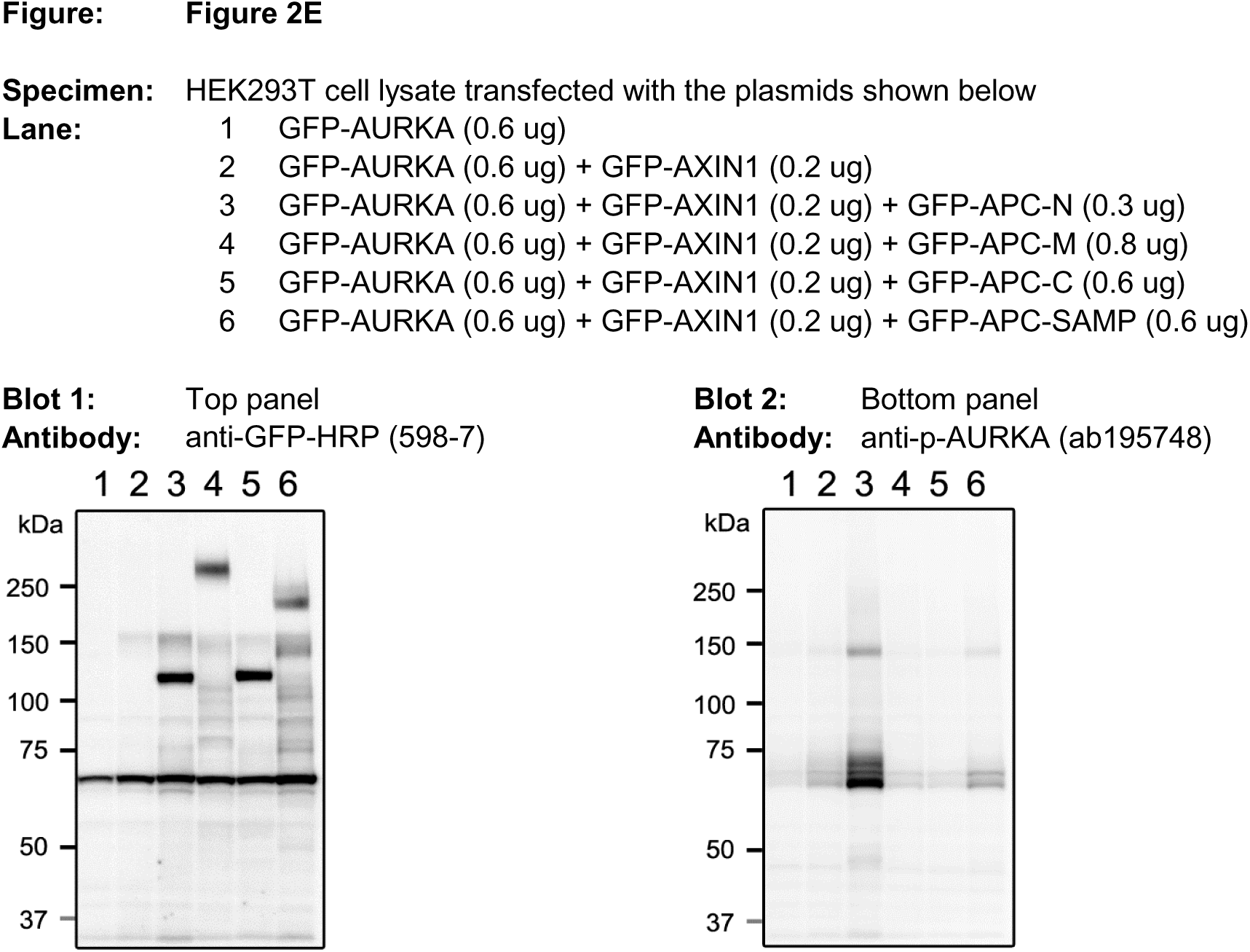
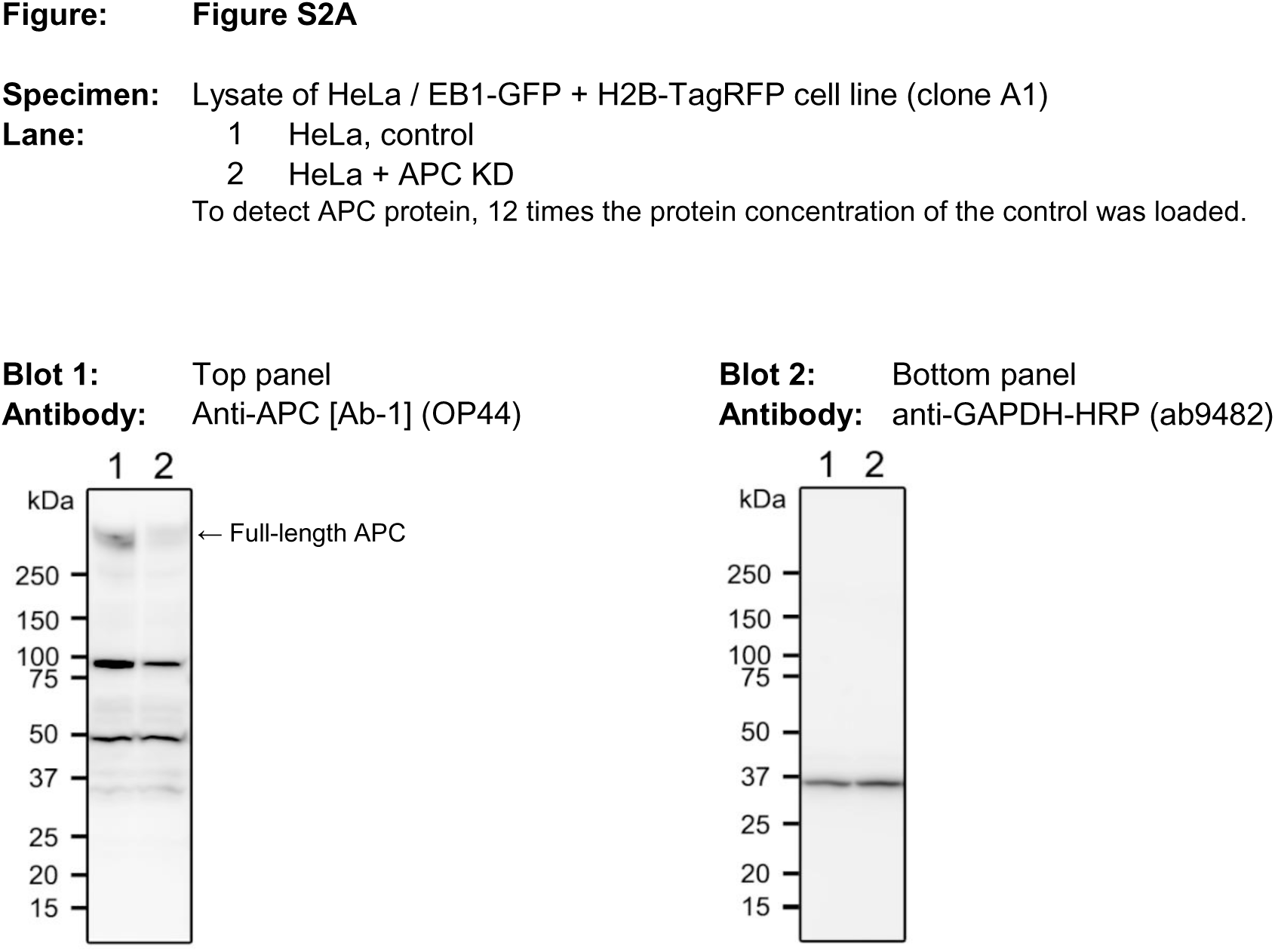
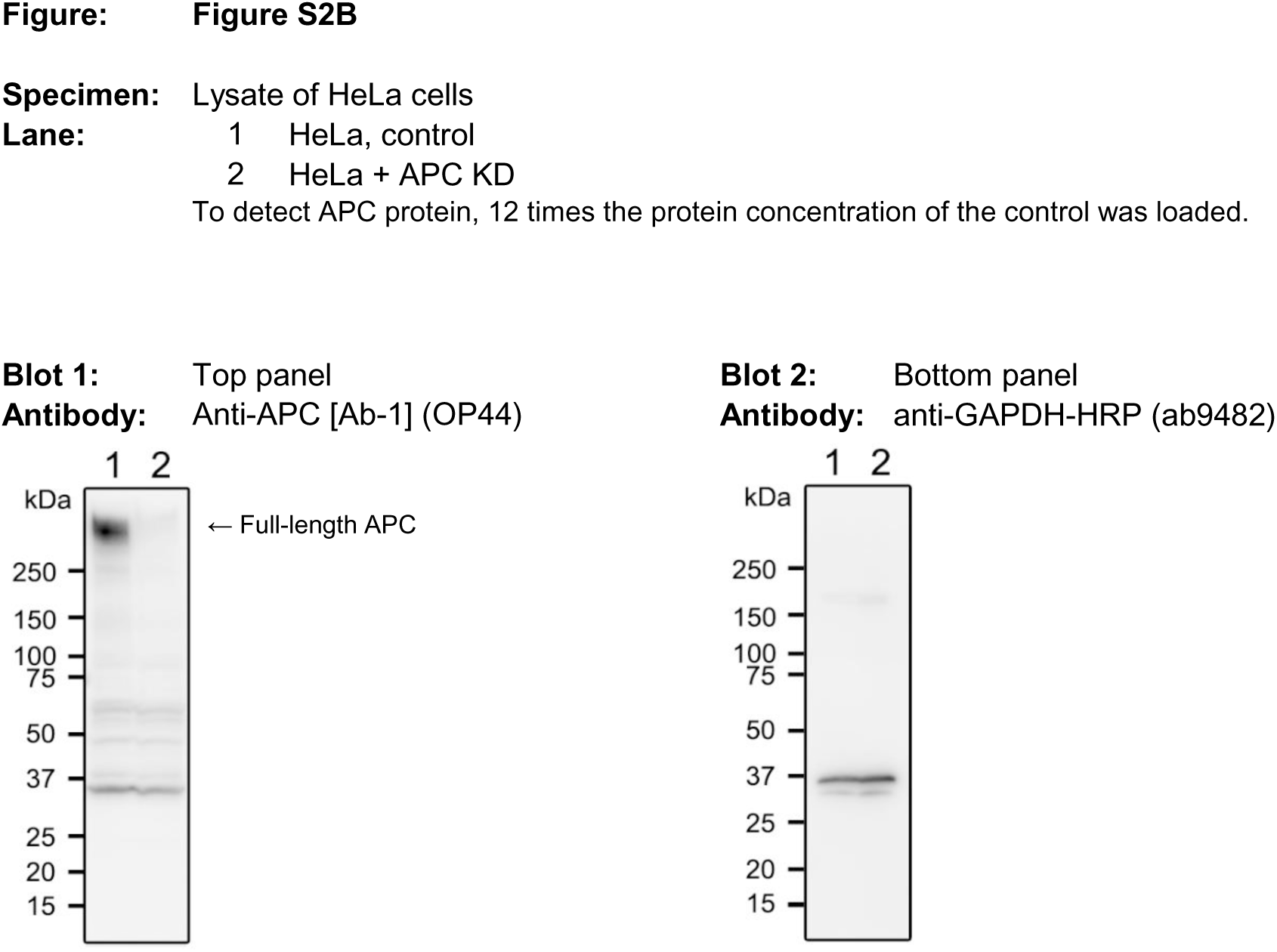
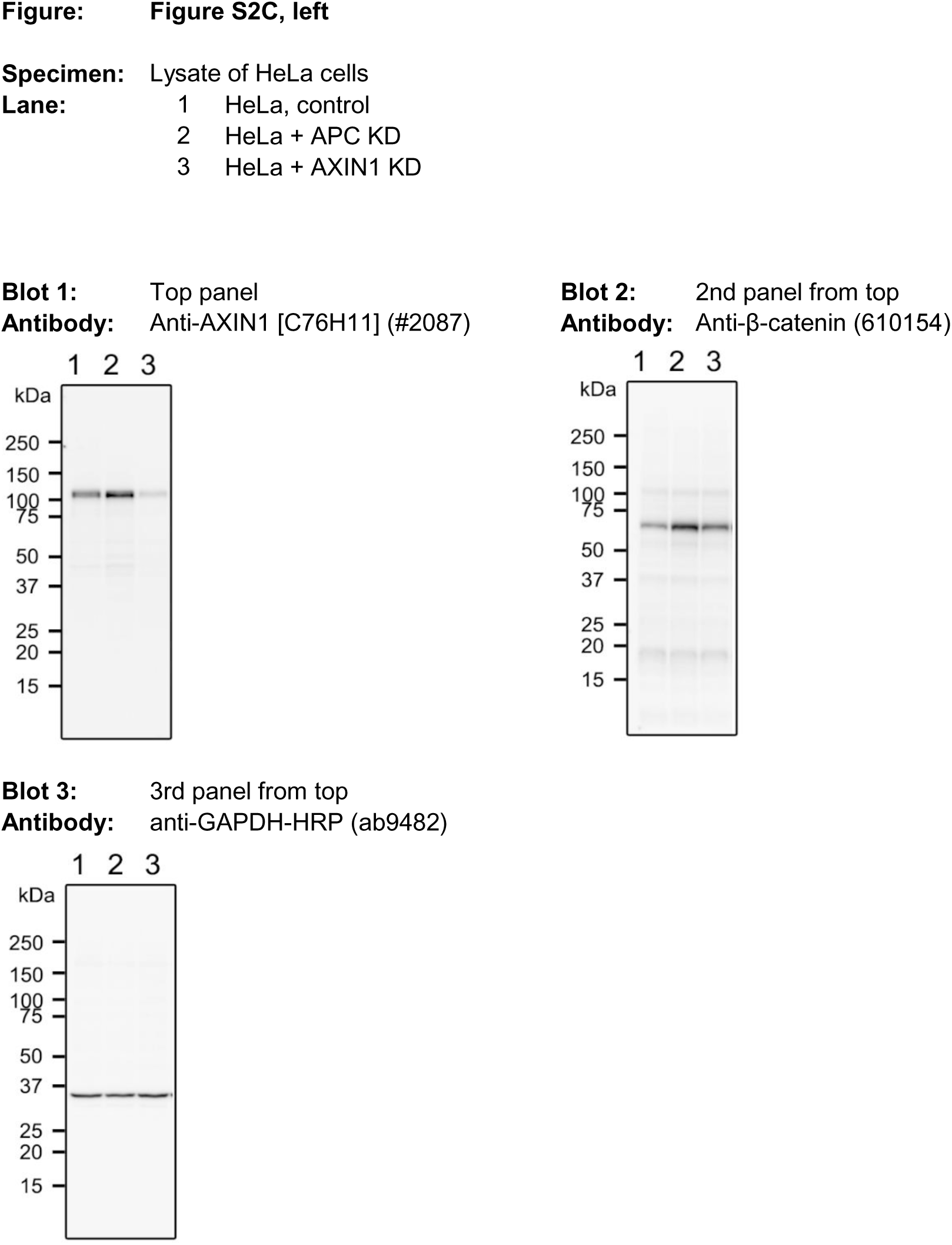
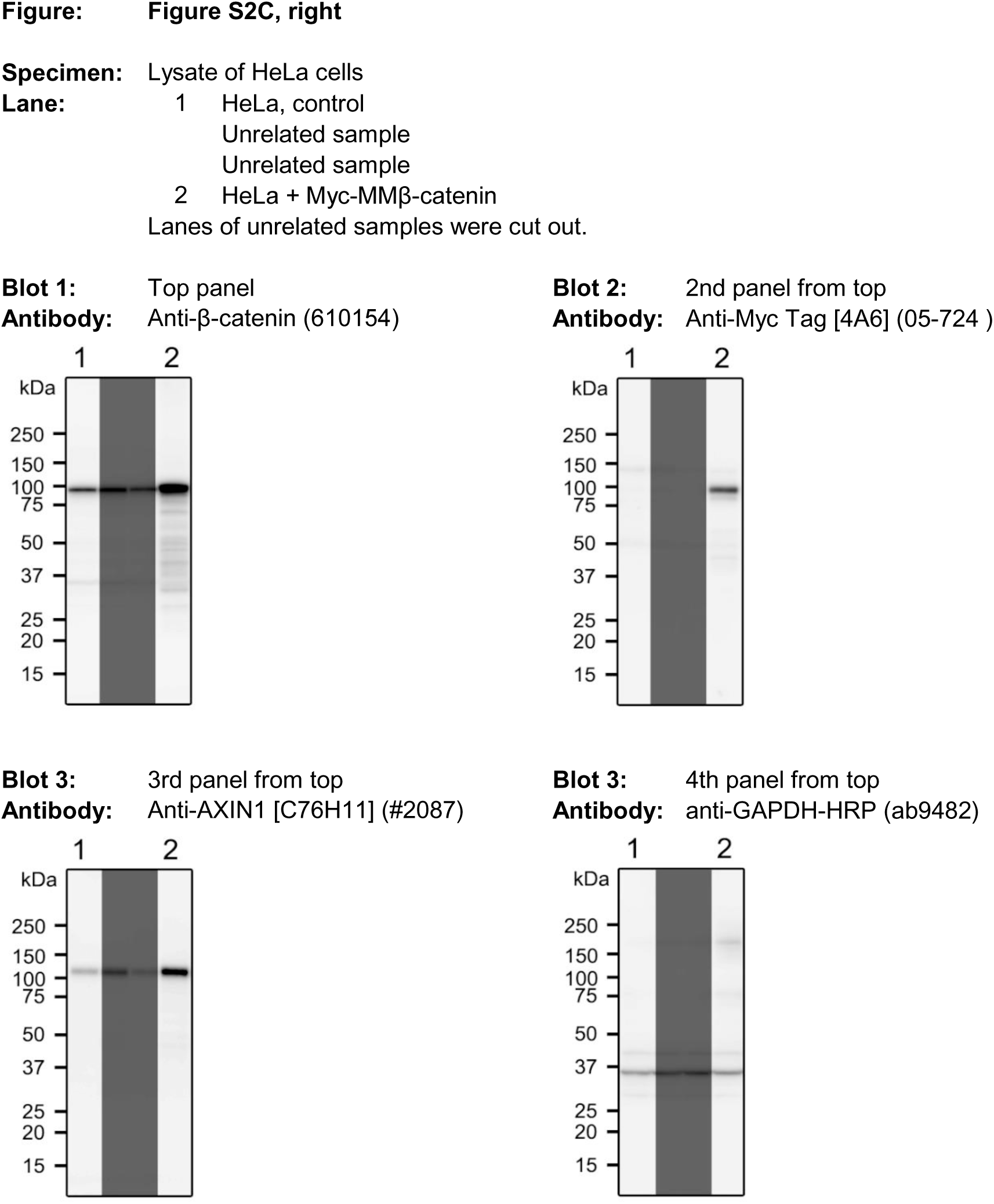
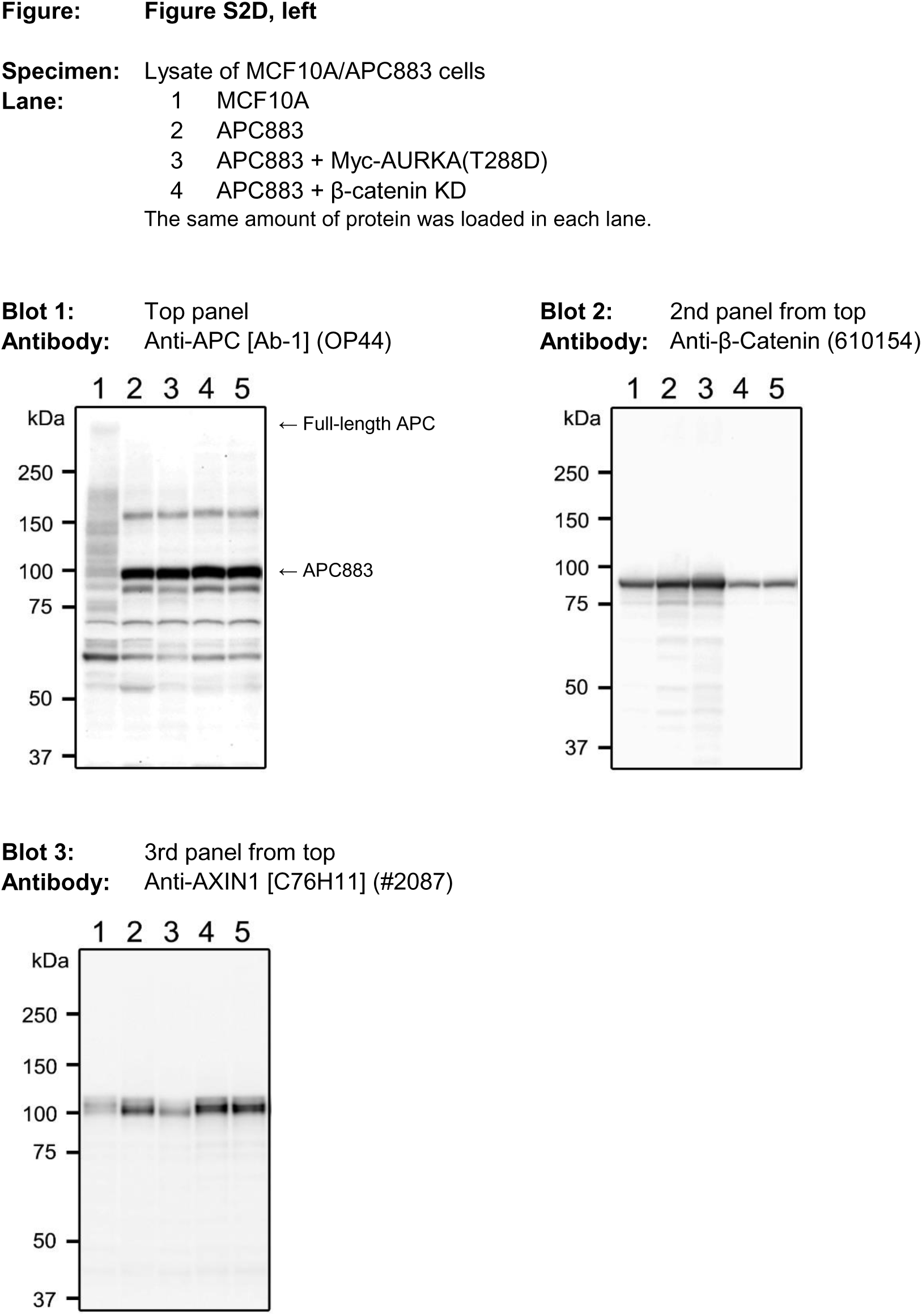
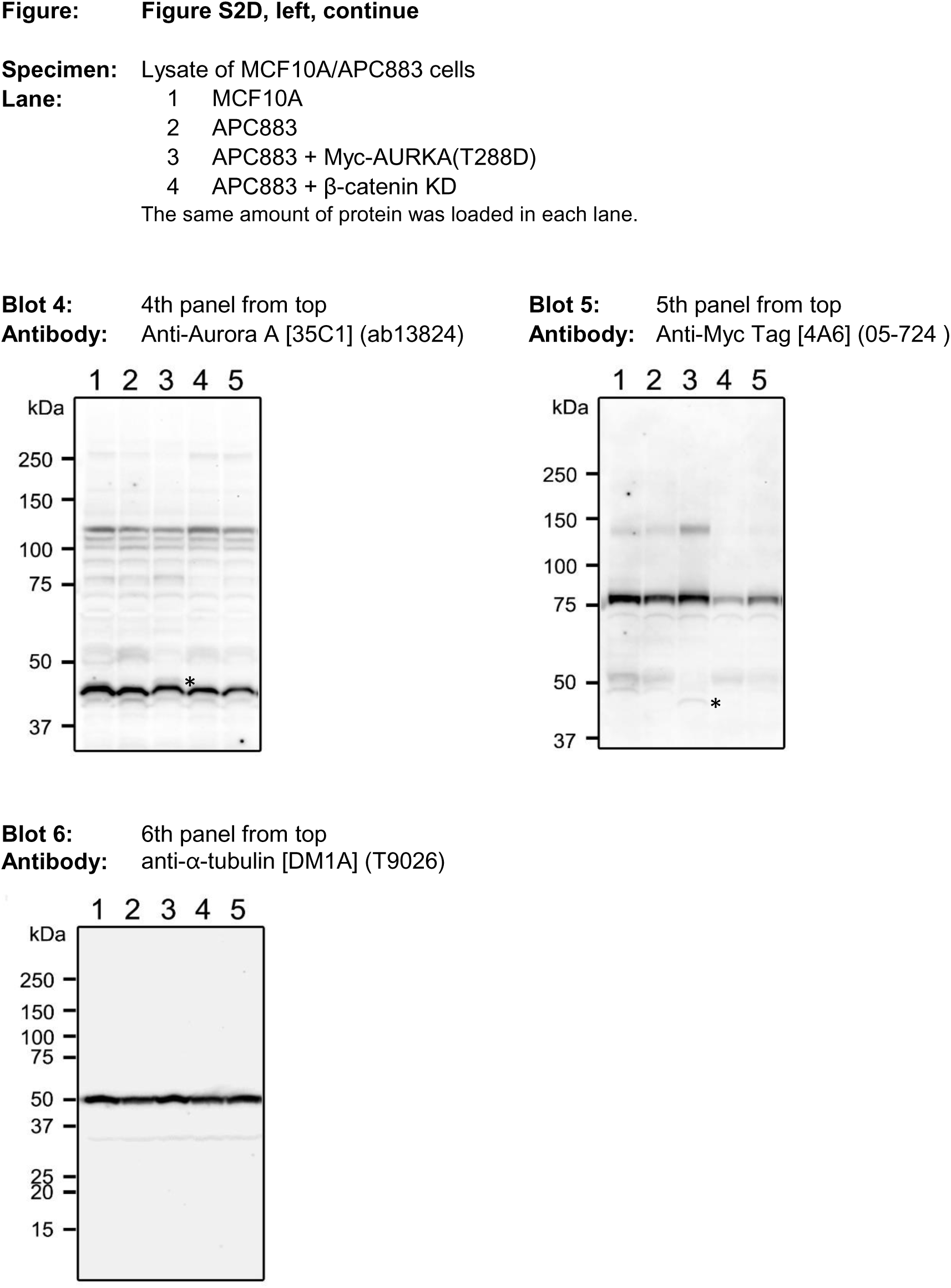
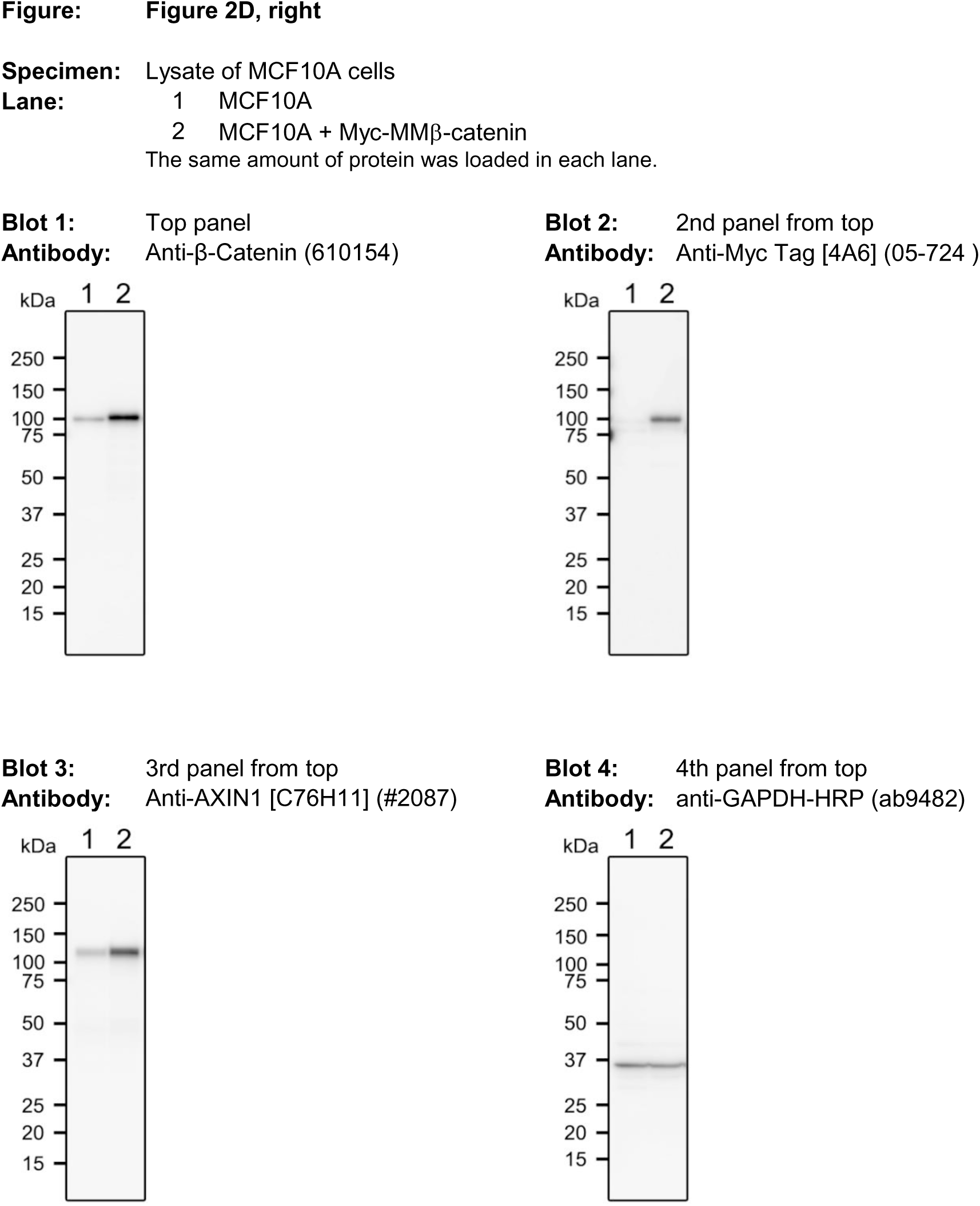
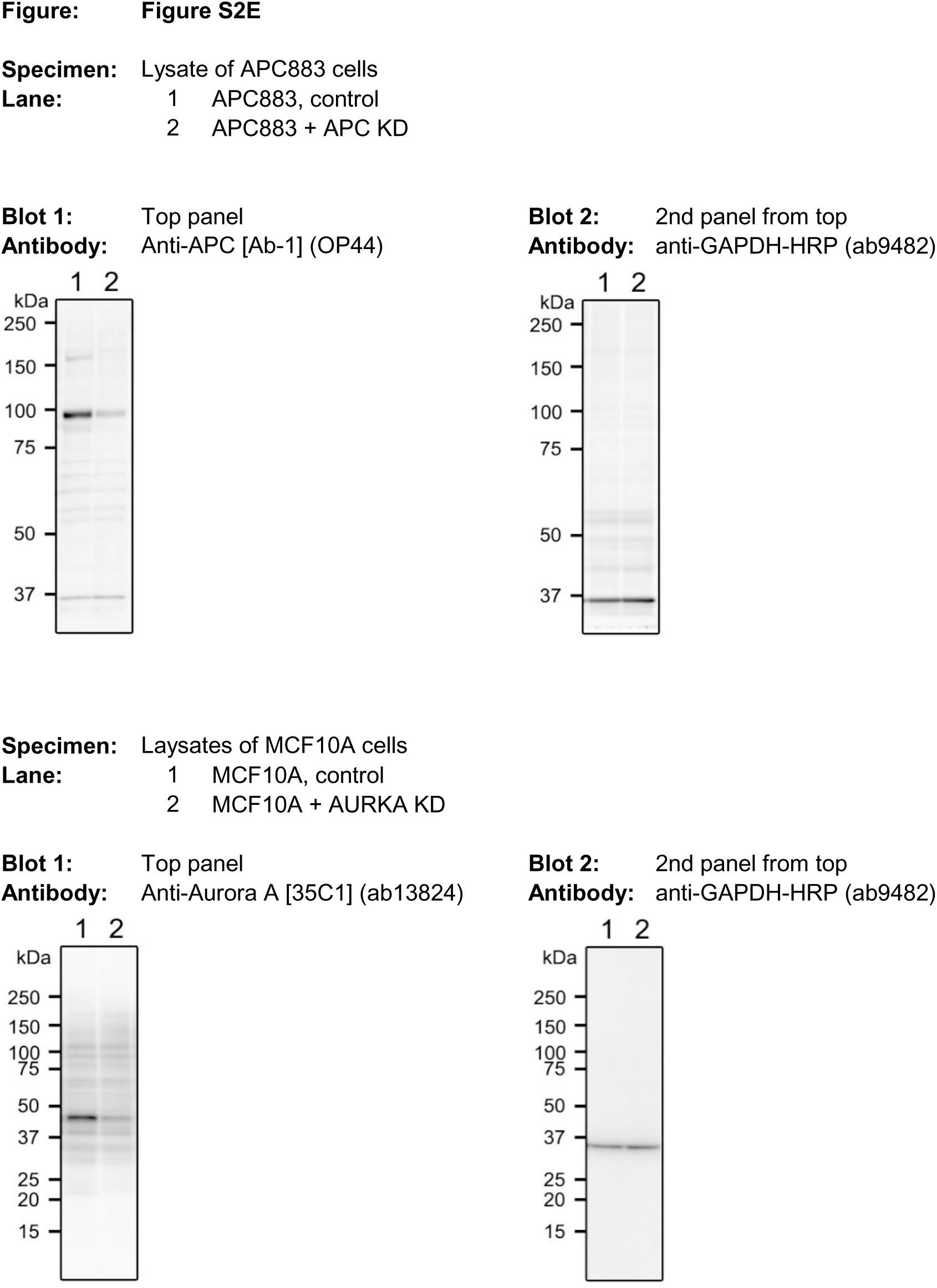
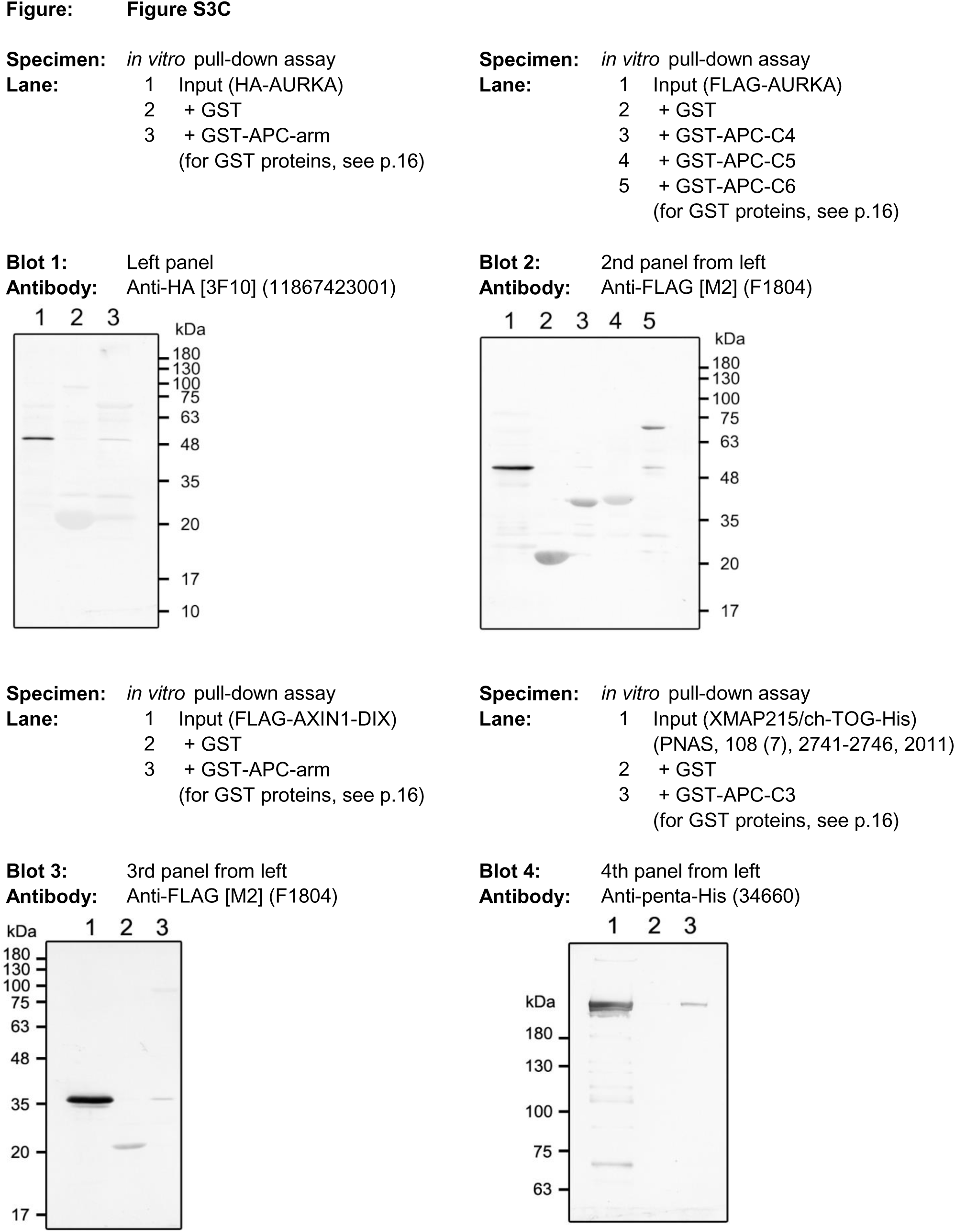
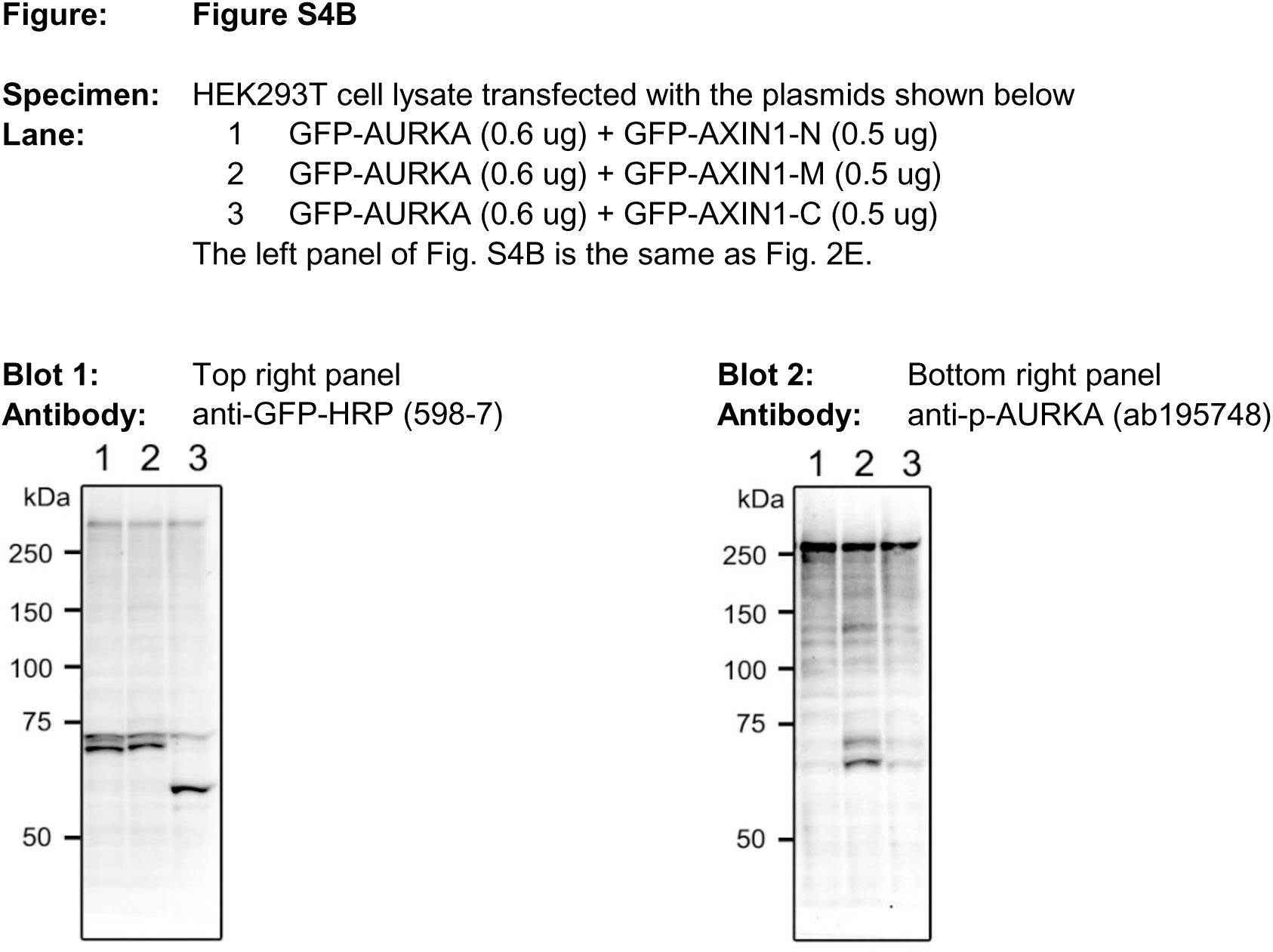
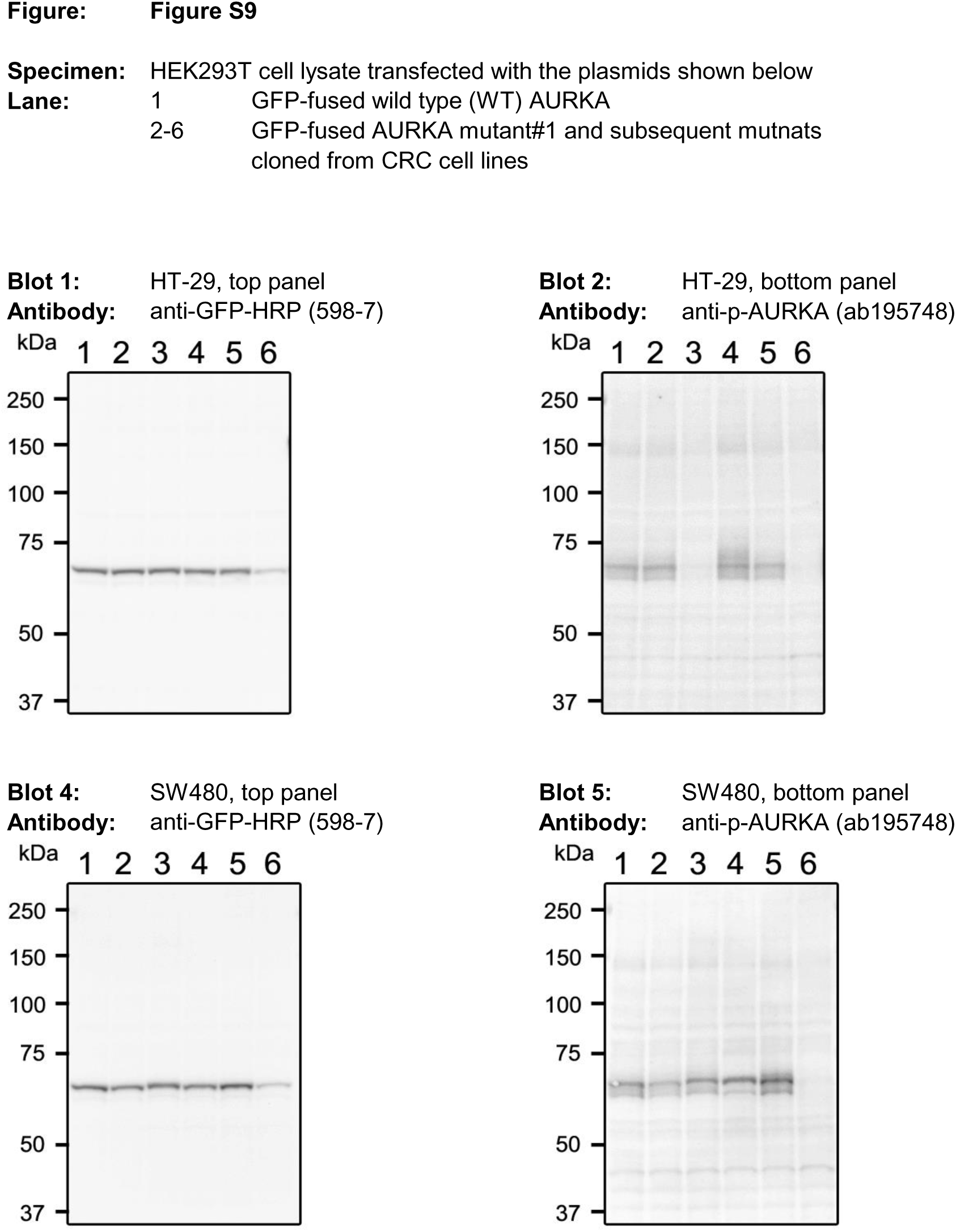
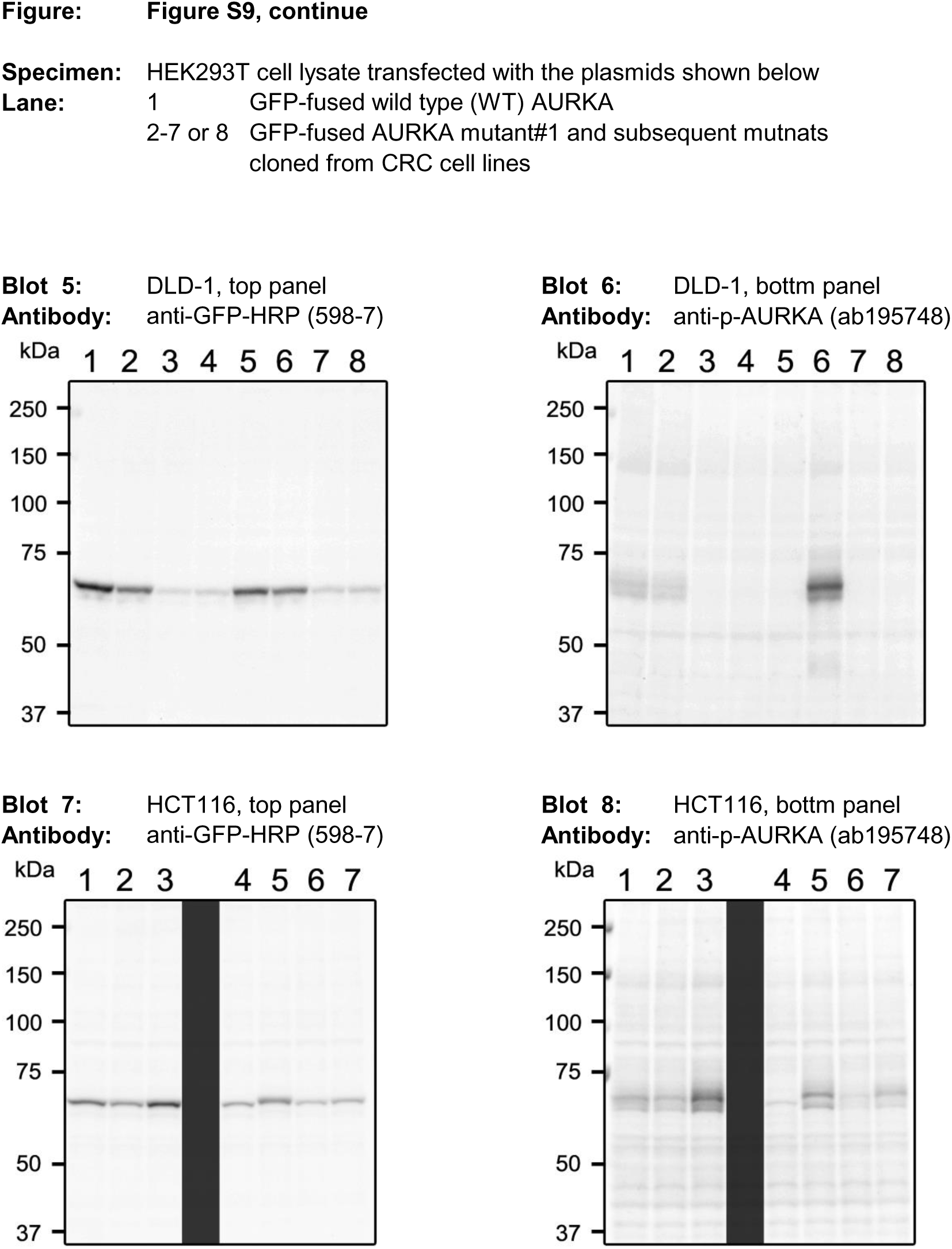
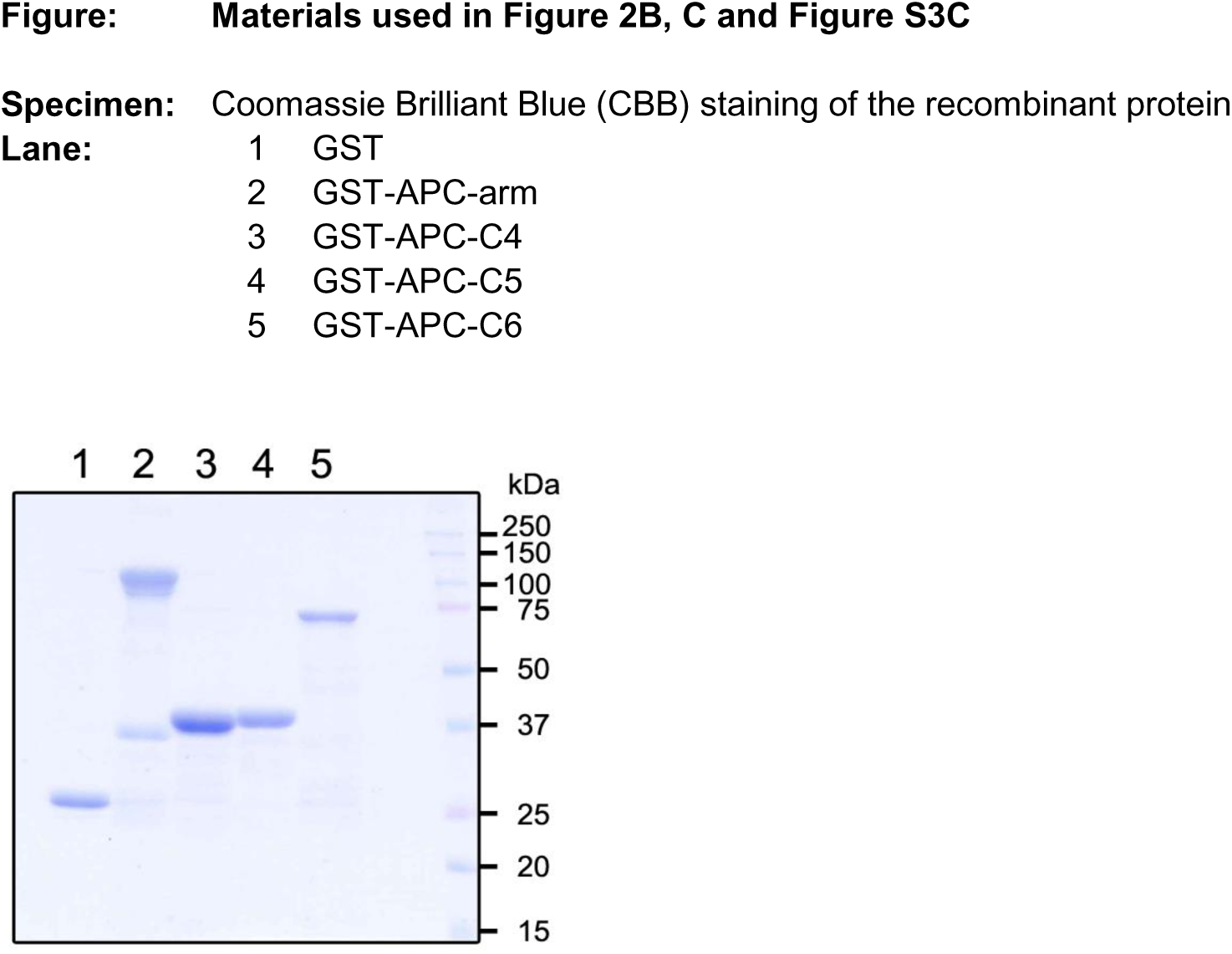
Uncropped versions of all gels and blots.

**Table S3.**
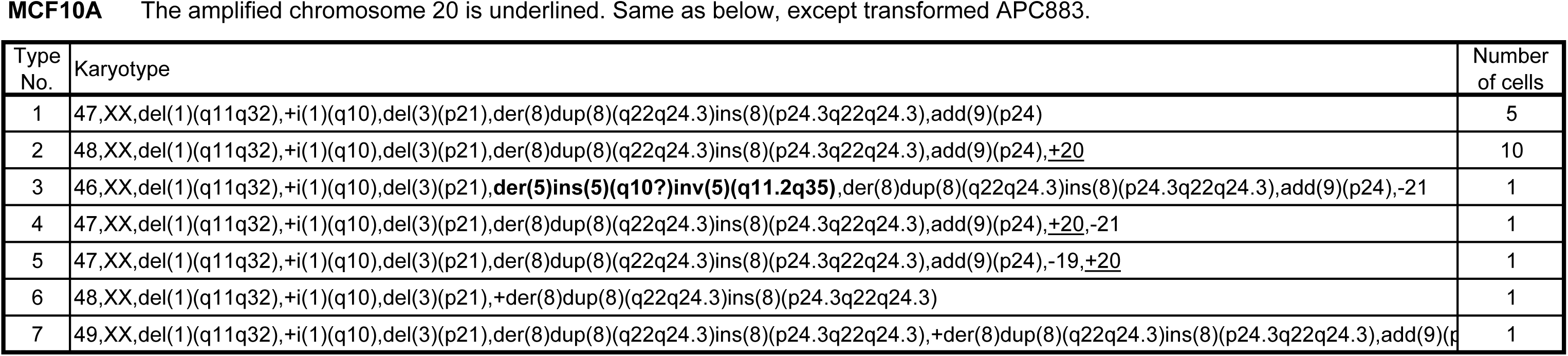
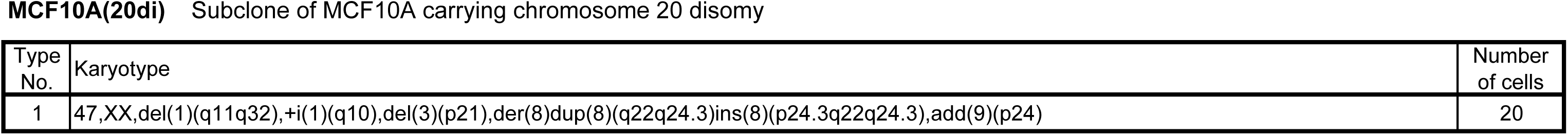
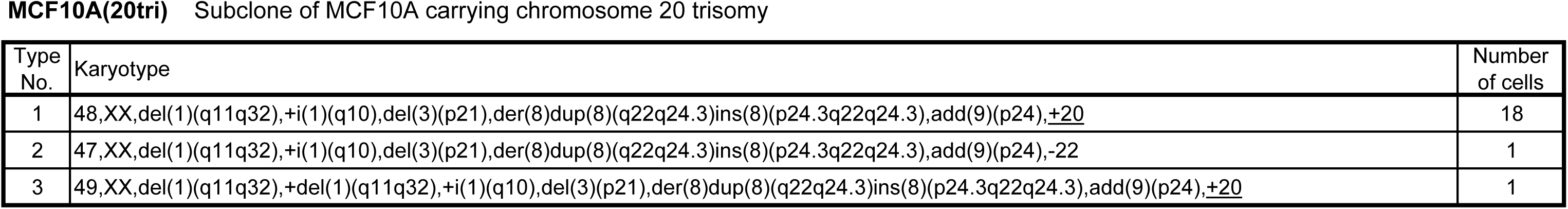
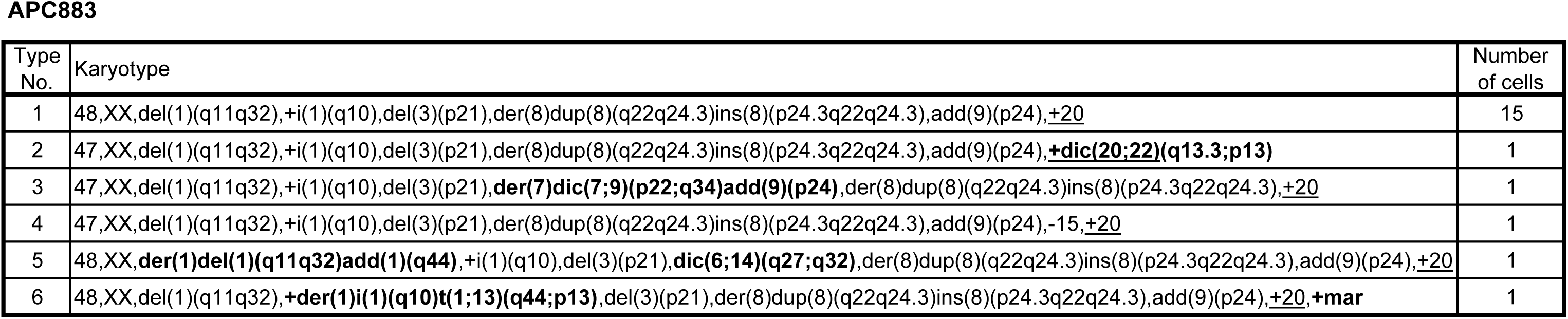
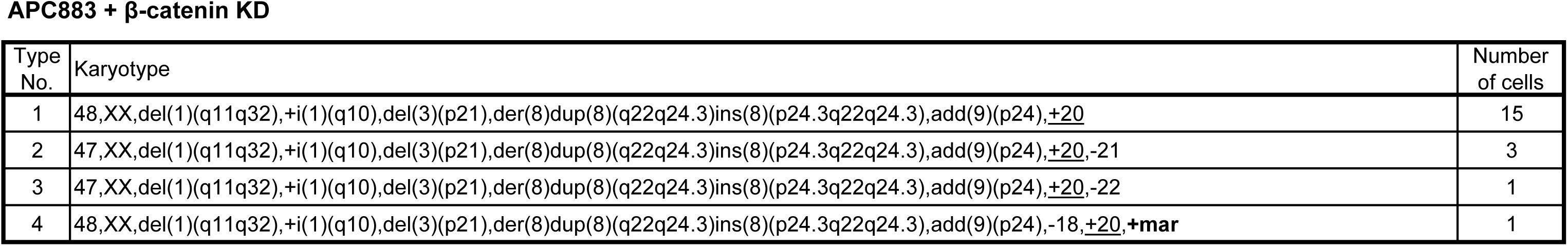
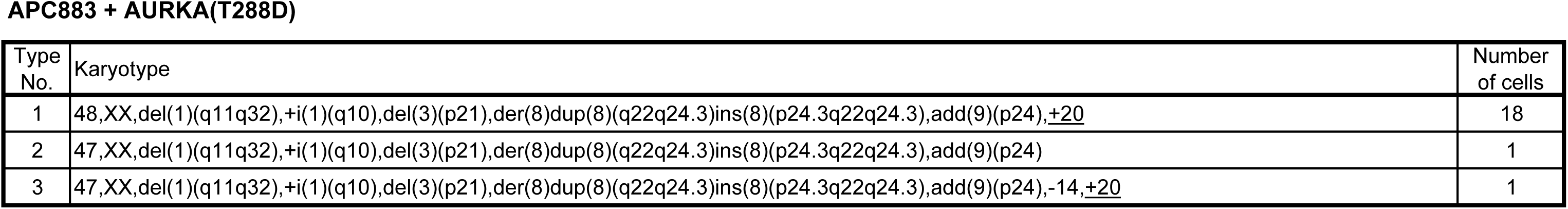
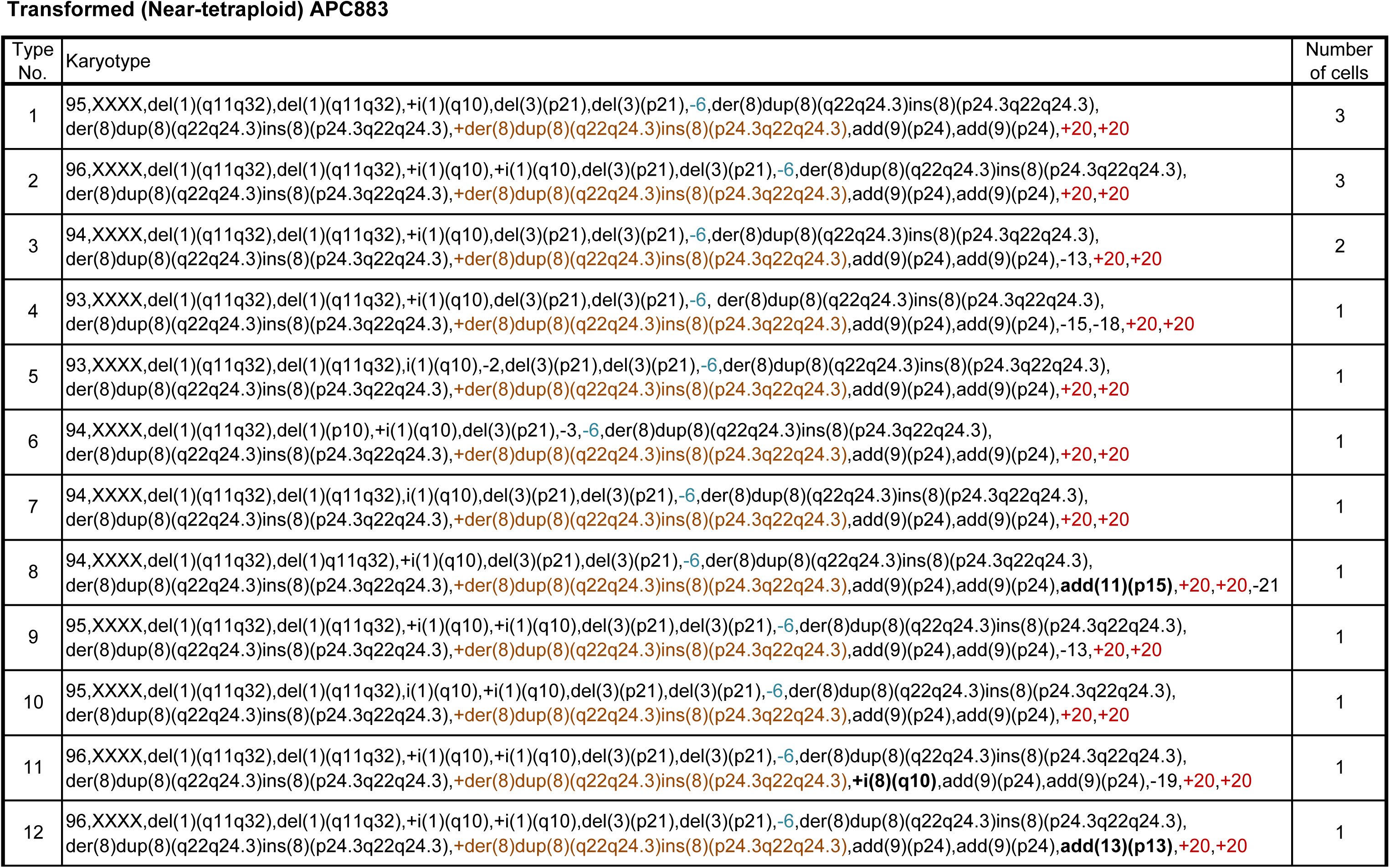
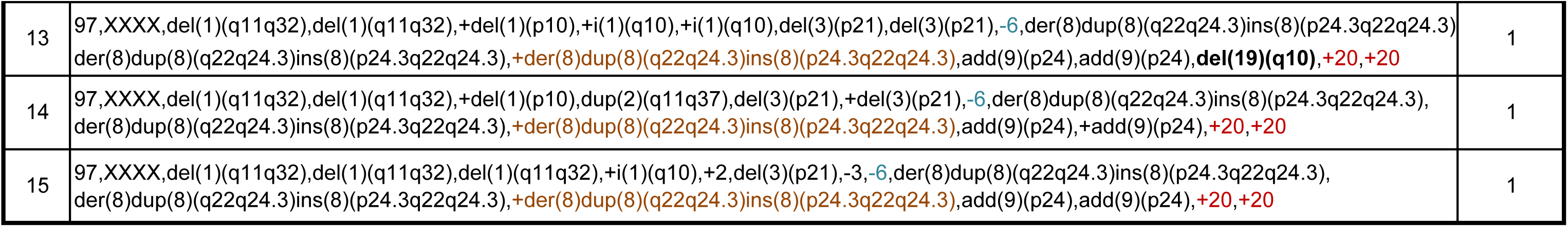
Full descriptions of the karyotypes of cell lines MCF10A and APC883, their subclones, and transfectants. Newly generated chromosomes, which included truncation and chromosome rearrangements, such as translocation, duplication, and telomere association, are shown in bold. For the spontaneously transformed, near tetraploid APC883 cells, the common number imbalance of chromosomes 6, 8, and 20 is indicated by color codes identical to the box color in Fig. S8E: cyan, three chromosome 6s; brown, five chromosome 8s; red, six chromosome 20s.

**Movie S1.**

Three-dimensional timelapse images of a mitotic HeLa cell that expressed EB1-GFP and H2B-TagRFP collected by lattice light sheet microscopy at 10-s intervals are represented in a 3D space. Time scale: h.m.s.

**Movie S2.**

Three-dimensional timelapse images of an APC-depleted mitotic HeLa cell that expressed EB1-GFP and H2B-TagRFP collected by lattice light sheet microscopy at 10-s intervals are represented in a 3D space. Time scale: h.m.s.

**Movie S3.**

Three-dimensional timelapse images of control and APC-depleted mitotic HeLa cells shown in Movies S1 and S2 were projected onto a 2D plane and combined for comparison. Time scale: h.m.s.

**Movie S4.**

Representative phase contrast timelapse videos of MCF10A(20di), MCF10A(20tri), APC883, and APC883+β-catenin KD, APC883+AURKA(T288D), APC883+APC KD, APC883+AURKA KD and transformed APC883 cells that spontaneously appeared after >5 months of passaging, shown in Fig. 5A, E. Timelapse images at 10-min intervals for 50 h are displayed from videos that recorded at 5-min intervals. Time scale: h:m:s.ms.

**Movie S5.**

Representative phase contrast timelapse videos of MCF10A, MCF10A+AURKA KD and MCF10A+MMβ-catenin cells, shown in Fig. S2F. Timelapse images at 10-min intervals for 50 h are displayed from videos that recorded at 5-min intervals. Time scale: h:m:s.ms.

**Code S1.**

Codes for TCGA data analysis. This folder also contains the data used to create the diagram with the attached codes.

## Notes

### Summary of Updates

We have conducted a comprehensive whole cancer genome analysis, revealing specific chromosomal rearrangement patterns associated with APC-mutant cancers. This analysis provides new insights into the genomic alterations that accompany APC loss. Additionally, we have extensively revised the interpretation of experimental results and the discussion. These updates clarify the biological implications of our findings and refine the conclusions drawn from the data.

