## Supplementary material for "Adaptive fitness in *APC*-mutant cells requires chromosome rearrangements": Fig1C: functionlist.py

```
import numpy as np
import sys,math,os
import matplotlib.pyplot as plt
from mpl_toolkits.mplot3d import axes3d
from mpl_toolkits.axes_grid1 import make_axes_locatable, axes_size
import matplotlib as mpl
import matplotlib.gridspec as grd
from matplotlib.ticker import FormatStrFormatter
import matplotlib as mpl
from matplotlib import ticker

mpl.rcParams.update({'font.size': 10})
mpl.rcParams.update({'axes.titleweight': 'medium'})
mpl.rcParams.update({'axes.titlesize': 15})
mpl.rcParams.update({'axes.labelweight': 'medium'})
mpl.rcParams.update({'axes.labelsize': 12})
asp = 'auto'	#aspect ratio
inter = 'spline16'	#interpolation
cmap = plt.get_cmap('hot')


def initialize(fn,types): 
    """load data and sort them according to track_id and time"""
    data = np.loadtxt("data/"+types+'/'+fn, delimiter=',', skiprows=4, dtype=np.str_, usecols=[0,1,2,6,7])
    data = data[np.all(data != '', axis=1)]
    pre_filename = "data/"+fn
    np.savetxt(pre_filename, data, delimiter=",", fmt="%s")
    dtype = [('x', float), ('y', float), ('z', float), ('time', int),
    ('track_id', int)]    
    data = np.loadtxt(pre_filename, dtype=dtype, delimiter=',')
    data=np.sort(data, order=['track_id', 'time'])
    return data

def plot_spot(dat,m,clr,sz): 
    """3D scatter plot of all spots
    (coordinates,marker,markercolor,markersize)
    """
    fig = plt.figure(1)
    ax =fig.gca(projection='3d')
    ax.scatter(dat['x'],dat['y'],dat['z'],marker=m,c=clr,s=sz,edgecolor='None')
    return ax

def importspots(types,channel,cell,plot):
    """import spots data and plot"""
    filename1 = types+str(channel)+'spot'+str(cell)+".csv" 
    filename2 = types+str(channel)+'cen'+str(cell)+".csv" 
    spot = initialize(filename1,types) #EB1 spots
    cent = initialize(filename2,types) #centrosome spots
    if plot:
        ax=plot_spot(spot,'.','b',1) 
        ax=plot_spot(cent,'o','r',50) 
        ax.set_title(types+' channel '+str(channel)+'cell'+str(cell))
    return spot,cent


def rotate(point,c1,c2): 
    """rotational projection of spots data
    (spots coordinate,centrosome1 coordinate,centrosome2 coordinate)
    """
    axis = c2-c1 #centrosome axis
    plane1 = [axis[1],-axis[0],0] #vector perpendicular to plane
    plane2 = np.cross(plane1,axis) #vector along the plane but perpendicular to plane1
    cs = point-c1
    dot1 = np.dot(axis,cs)
    axism = np.linalg.norm(axis)
    ss = cs-np.multiply(dot1/axism**2.,axis) #vector perpendicular to axis and goes through given point
    dotp1 = np.dot(plane1,ss)
    dotp2 = np.dot(plane2,ss)
    theta = math.acos(abs(dotp2)/np.linalg.norm(ss)/np.linalg.norm(plane2))
    if (dotp2<0):   theta = theta+np.pi
    if (dotp1*dotp2>=0): theta = -theta
    rotated = np.dot(rotation_matrix(axis,theta), point)
    return rotated

def rotation_matrix(axis, theta):
    """Return the rotation matrix associated with counterclockwise rotation about the given axis by theta radians using euler-rodrigues formula """
    axis = np.asarray(axis)
    theta = np.asarray(theta)
    axis = axis/math.sqrt(np.dot(axis, axis))
    a = math.cos(theta/2.0)
    b, c, d = -axis*math.sin(theta/2.0)
    aa, bb, cc, dd = a*a, b*b, c*c, d*d
    bc, ad, ac, ab, bd, cd = b*c, a*d, a*c, a*b, b*d, c*d
    return np.array([[aa+bb-cc-dd, 2*(bc+ad), 2*(bd-ac)],
                     [2*(bc-ad), aa+cc-bb-dd, 2*(cd+ab)],
                     [2*(bd+ac), 2*(cd-ab), aa+dd-bb-cc]])

def ave_cent(cent): 
    """time average centrosome position"""
    tid = np.unique(cent['track_id'])
    ls = len(cent[cent['track_id']==tid[0]]),len(cent[cent['track_id']==tid[1]])
    if (ls[0]!=ls[1]): #check whether number of centrosomes data are the same
        idx = ls.index(min(ls))
        minls = ls[idx]
        #check whether time frame is continuous or not
        if ([minls in cent[cent['track_id']==tid[idx]]['time']]):
            rows = minls
        else:
            print( 'centrosome not continuous in time, missing frame')
    else:
        leng = max(cent['time'])
        if (cent[cent['time']==leng].shape[0]==2):
            rows = leng
        else:
            print( 'centrosome not continuous in time, missing frame')
    c1 = np.column_stack((cent['x'][0:rows],cent['y'][0:rows],cent['z'][0:rows]))
    c2 = np.column_stack((cent['x'][rows:rows*2],cent['y'][rows:rows*2],cent['z'][rows:rows*2]))
    return np.mean(c1,axis=0),np.mean(c2,axis=0)


def rotatespots(dat,cent,plot): 
    """given spots and centrosome, rotate to 2D plane """
    c1,c2 = ave_cent(cent)
    cp = c1
    c1 = c1-cp 
    c2 = c2-cp
    c2xy =[c2[0],c2[1],0]
    c2yz =[0,c2[1],c2[2]]
    alpha = ((-1)**((c2[0]*c2[1]<0)))*math.acos(abs(np.dot(c2xy,[0,1,0]))/np.linalg.norm(c2xy))
    beta = ((-1)**((c2[1]*c2[2]<0)))*math.acos(abs(np.dot(c2,[0,0,1]))/np.linalg.norm(c2))
    rm =  np.dot(rotation_matrix([1,0,0],beta+np.pi*0.5), rotation_matrix([0,0,1],alpha)) 
    c2 = np.dot(rm, c2)#rotate c2 to y axis
    postn = np.empty((0,2),float)
    dur = max(dat['time'])
    for p in range(len(dat)): 
        mat = np.array([dat['x'][p]-cp[0], dat['y'][p]-cp[1], dat['z'][p]-cp[2]]) #translate spot
        s = np.dot(rm,mat)#rotate spot to y 
        rotated = rotate(s,c1,c2)# rotate to plane
        postn = np.r_[postn,np.column_stack(([rotated[1]],[rotated[2]]))]
    if (c2[1]<0):
        c2[1]=-c2[1]
        postn[:,0]=-postn[:,0]
    X,Y=postn.T
    if plot:
        f,ax = plt.subplots()
        ax.scatter(c1[1],c1[2],s=100, c='c')
        ax.scatter(c2[1],c2[2],s=100, c='c')
        ax.scatter(postn[:,0],postn[:,1],s=5,marker='.', c='b',edgecolor='None')
        ax.set_aspect('equal')
        plt.tight_layout()
        ax.set_xticks(np.arange(int(min(X)-1),int(max(X))+2,1))
        ax.set_yticks(np.arange(int(min(Y)-1),int(max(Y))+2,1))
        ax.set_ylim(min(Y),max(Y))
        ax.set_xlim(min(X),max(X))
        ax.set_xlabel('x')
        ax.grid(which='both')
    return X,Y,c1[1:],c2[1:],dur

def plotfig(x,y,c1,c2,num,plot): 
    """plot spots with rescale x axis """
    xx = x
    xmin = min(xx[xx<0])
    xmax = max(x)
    ymin = c1[1]
    ymax = 10
    xedges = np.linspace(xmin,xmax,num)
    yedges = np.linspace(ymin,ymax,num)    
    if plot:
        ax = plt.subplot()
        ax.scatter(x,y,s=5,marker='.',c='c',edgecolor='None')	
        ax.scatter(c1[0],c1[1],s=50, c='c')
        ax.scatter(c2[0],c2[1],s=50, c='c')
        ax.set_aspect(asp)
        ax.set_ylim(ymin,ymax)
        ax.set_xlim(xmin,xmax)    
        major_ticksx = np.linspace(xmin,xmax,4)
        major_ticksy = np.linspace(ymin,ymax,4)
        ax.set_xticks(major_ticksx)
        ax.set_xticks(xedges, minor=True)
        ax.set_yticks(major_ticksy)
        ax.set_yticks(yedges, minor=True)
        ax.grid(which='both')
        ax.set_title('rescale x-axis with grid')
    return xedges,yedges


def numberofspot(x,y,c1,c2,xed,yed,plot): 
    """return number of spot in each grid 
    (xed and yed are the min and max edges of the spots data)   """
    inter = 'spline16'
 #   xx=-(x[x>0.5]-1) #half
#    x[x>0.5] = xx #half
    H, xedges, yedges = np.histogram2d(x,y, bins=[xed, yed],normed=False)
    myextent  =[xedges[0],xedges[-1],yedges[0],yedges[-1]]
    if plot:
        ax = plt.subplot()
        p = ax.imshow(H.T,origin='low',extent=myextent,interpolation=inter,cmap=cmap,aspect=asp)
        ax.scatter(c1[0],c1[1],s=50, c='c')
        ax.scatter(c2[0],c2[1],s=50, c='c')
#        ax.set_xlim(-0.5,0.5)
        ax.set_xlim(-0.5,1.5)
        ax.set_axis_bgcolor('black')
        divider = make_axes_locatable(ax)
        cax = divider.append_axes("right", size="10%", pad=0.05)
        fig=plt.gcf()
        cb = fig.colorbar(p,cax=cax)
        cb.set_label("number of points")
        ax.set_title('heat map of number of spots')
    return H


def Ndensity(H,xedges,yedges,c1,c2,dur,num,plot): 
    """return normalized density from number of spots """
    #bin volume
    const = np.pi*((yedges[1]-yedges[0])**2)*abs(xedges[1]-xedges[0])
    vol = [const*(2*i+1) for i in range(num-1)]
    #number density= spots number /bin's 3D volume
    ND = np.array([ H.T[i]/vol[i] for i in range(num-1)])
    #NDpt = number density /duration
#    NDpt = np.array([ND[i]/dur for i in range(num-1)])
    NDpt = ND/dur
    tot = NDpt.sum()
    #normalized density= NDpt / sum of NDpt
    normH = NDpt/tot#np.array([NDpt[i]/tot for i in range(num-1)])
    myextent  =[xedges[0],xedges[-1],yedges[0],yedges[-1]]
    vmax = normH[list(yedges>1.).count(0)::].max()#
    if plot:
        ax = plt.subplot()
        p = ax.imshow(normH,origin='low',extent=myextent,interpolation=inter,cmap=cmap,aspect=asp,vmin=0,vmax=vmax)
        ax.scatter(c1[0],c1[1],s=50, c='c')
#        ax.scatter(c2[0],c2[1],s=50, c='c')
#        ax.set_xlim(-0.5,1.5)
        ax.set_xlim(-0.5,0.5)
        ax.set_axis_bgcolor('black')
        divider = make_axes_locatable(ax)
        cax = divider.append_axes("right", size="10%", pad=0.05)
        fig=plt.gcf()
        cb = fig.colorbar(p,cax=cax)
        cb.set_label("normalized density")
    return normH
```
