## Supplementary material for "Adaptive fitness in *APC*-mutant cells requires chromosome rearrangements": Fig1C: sig_test.py

```
import numpy as np
import sys,math,os
from functionlist import * #(refer to functionlist.py for detail)

#from spot to density to norm desntiy
typess = ['control','apc']
channels = [1,2]
apccell = [[1,2,3],[1,2,3,4,5]]
controlcell = [[1,2,3],[1,2,3,4,5,6,7]]
spotdata = []
dts=[0.755,1.51]#time duration of each frame
totduration=[]

num = 40 #number of bin in each axis
ed=[-0.733, 1.629, 0, 10] #[xmin,xmax,ymin,ymax] of edges
#density = [[],[]]
#rescale centrosome positions:
c1 = [0,0]
c2 = [1,0]
xedges = np.linspace(ed[0],ed[1],num)
yedges = np.linspace(ed[2],ed[3],num)

for types in typess:
    print('types=',types)
    xs = np.array([])
    ys = np.array([])
    if types=='control': cellist=controlcell
    else: cellist=apccell
    totdur=0
    for ch in channels:
        print('channel=',ch)
        cells= cellist[ch-1]
        for cell in cells:
            spot,cent=importspots(types,ch,cell,False)
            sx,sy,C1,C2,tf=rotatespots(spot,cent,False)
            dur = tf*dts[ch-1]
            print('cell',cell,' duration=',dur            )
            xs = np.r_[xs,sx/C2[0]]
            ys = np.r_[ys,sy]  
            totdur+=dur
            totduration.append(totdur)
            H=numberofspot(xs,ys,c1,c2,xedges,yedges,False)    
            density = Ndensity(H,xedges,yedges,c1,c2,totdur,num,False)#/1e-3#rescaling factor for plotting
            print(density.shape)
            if (ch ==1 and cell ==1):
                Density =density
            else:
                Density = np.dstack([Density,density])
    print(Density.shape)
    if types =='control':
        control_density_mean = np.mean(Density,axis=2)
        control_density_var = np.var(Density,axis=2)
    elif types =='apc':
        apc_density_mean = np.mean(Density,axis=2)
        apc_density_var = np.var(Density,axis=2)
    
#t test for each grid, input are mean and variance of each grid
#two-sided t test are performed using function from 
#https://docs.scipy.org/doc/scipy/reference/generated/scipy.stats.ttest_ind_from_stats.html#scipy.stats.ttest_ind_from_stats
from scipy.stats import ttest_ind_from_stats
PV = np.ones((39,39))
for i in range(39):
    for j in range(39):
        mea1 = control_density_mean[i,j]
        mea2 = apc_density_mean[i,j]
        var1 = control_density_var[i,j]
        var2 = apc_density_var[i,j]
        stat,pvalue = ttest_ind_from_stats(mean1=mea1, std1=np.sqrt(var1), nobs1=10,
                     mean2=mea2, std2=np.sqrt(var2), nobs2=8)
        if np.isnan(pvalue) == False: PV[i,j] = pvalue

PVf = PV.flatten()
plt.hist(PVf)
nogrid = len(PVf)
PVft = PVf[PVf!=1]
nogriddiff = len(PVft)#number of grid that show difference between control and apc
print('Number of grid with p<0.1 {:.1f}%'.format(len(PVft[PVft<0.1])*100/nogriddiff))
print('Number of grid with p<0.01 {:.1f}%'.format(len(PVft[PVft<0.01])*100/nogriddiff))
```
